# Systematic Integration of Structural and Functional Data into Multi-Scale Models of Mouse Primary Visual Cortex

**DOI:** 10.1101/662189

**Authors:** Yazan N. Billeh, Binghuang Cai, Sergey L. Gratiy, Kael Dai, Ramakrishnan Iyer, Nathan W. Gouwens, Reza Abbasi-Asl, Xiaoxuan Jia, Joshua H. Siegle, Shawn R. Olsen, Christof Koch, Stefan Mihalas, Anton Arkhipov

## Abstract

Structural rules underlying functional properties of cortical circuits are poorly understood. To explore these rules systematically, we integrated information from extensive literature curation and large-scale experimental surveys into a data-driven, biologically realistic model of the mouse primary visual cortex. The model was constructed at two levels of granularity, using either biophysically-detailed or point-neurons, with identical network connectivity. Both variants were compared to each other and to experimental recordings of neural activity during presentation of visual stimuli to awake mice. While constructing and tuning these networks to recapitulate experimental data, we identified a set of rules governing cell-class specific connectivity and synaptic strengths. These structural constraints constitute hypotheses that can be tested experimentally. Despite their distinct single cell abstraction, spatially extended or point-models, both perform similarly at the level of firing rate distributions. All data and models are freely available as a resource for the community.

## Introduction

Mechanisms connecting structural properties of cortical circuits to patterns of neural activity are poorly understood. Elucidating such mechanisms requires systematic data collection, sophisticated analyses, and modeling to “understand” this data. Such an understanding is always relative to a particular domain of interest – be it modeling the physics of highly excitable brain tissue composed of a myriad of heterogeneous neurons (Koch, 1999; Einevoll *et al*., 2013), mimicking the computations that lead to a particular set of firing rates (Yamins and DiCarlo, 2016), or diagnosing and ultimately curing psychiatric and neurological brain diseases. The first option – biologically realistic modeling – appears necessary to disentangle the extreme biological complexity of the cortex (Harris and Mrsic-Flogel, 2013; Harris and Shepherd, 2015; Amunts *et al*., 2016; Koch and Jones, 2016; Martin and Chun, 2016; Chevée and Brown, 2018; Einevoll *et al*., 2019).

Simulating cortical circuits has a long history (e.g. (Wehmeier *et al*., 1989; Zemel and Sejnowski, 1998; Troyer *et al*., 1998; Krukowski and Miller, 2001; Traub *et al*., 2005; Zhu, Shelley and Shapley, 2009; Potjans and Diesmann, 2014; Markram *et al*., 2015; Arkhipov *et al*., 2018; Joglekar *et al*., 2018; Schmidt *et al*., 2018; Antolík *et al*., 2019; Schwalger and Chizhov, 2019)), with models incrementally building upon their predecessors. The simulations described here are a further instance of this evolution toward digital simulacra that predict new experiments, are insightful, and ever more faithful to the vast complexity of cortical tissue, in particular its heterogeneous neuronal cell classes, connections, and *in vivo* activity.

We developed data-driven models of the mouse primary visual cortex (area V1), containing ∼230,000 neurons, to simulate *in silico* visual physiology studies with arbitrary visual stimuli (Fig. 1A). The choice to focus on the mouse visual system was given by the availability of substantial amounts of high-quality data on structural and *in vivo* functional properties of different cell classes, especially from the standardized experimental pipelines at the Allen Institute. However, the primary aim for developing these models is to provide a computational platform to study cortical structure and computation in general, under constraints of realistic biological complexity. Our models can be used as templates to develop models of other cortical areas in rodents or other species. We developed two V1 variants at two different levels of granularity: using biophysically detailed spatially extended compartmental models of 17 different cells classes (Gouwens *et al*., 2018) or Generalized Leaky Integrate and Fire (GLIF) point-neuron models of these 17 classes (Teeter et al., 2018). The latter variant can be run on a single CPU, making it broadly accessible to all neuroscientists. Both variants are constrained by experimental measurements and reproduce multiple observations from in-house Neuropixels high-density electrical recordings *in vivo* (Siegle *et al*., 2019).

**Figure 1:**
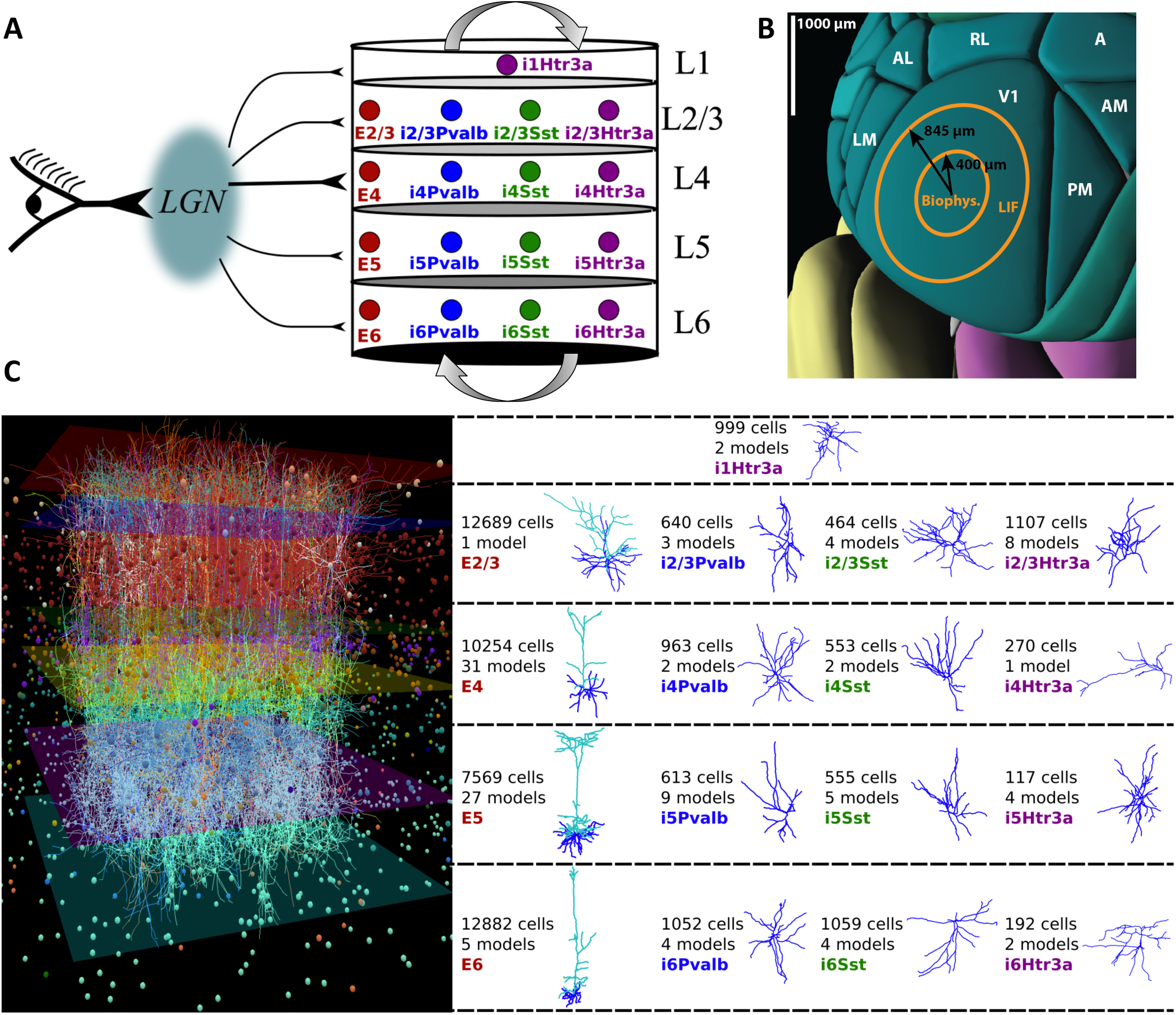
Overview of the V1 models. **A:** The models consist of one excitatory class and three inhibitory classes (Pvalb, Sst, Htr3a) in each of four layers - L2/3, L4, L5 and L6; L1 has a single Htr3a inhibitory class, for a total of 17 cell classes. Visual stimuli are conveyed by projections from the LGN (see Figs. 2 and 3). **B:** Mouse posterior cortex (from the Allen Brain Explorer), illustrating V1 and higher visual cortical areas, and the region covered by the models (400 μm radius for the core; 845 μm radius with surrounding annulus). **C:** Left: Visualization of the biophysically detailed network (1% of neurons shown). Right: Exemplary dendritic morphologies for each class. The total number of neurons per class as well as the number of neuron models that comprise every class are indicated. Every model has its own uniquely reconstructed morphology and electrical parameters. Models are selected from the Allen Cell Types Database (see Methods).

To demonstrate their potential, we here describe three predictions that emerged in the process of building and testing these models. First, the strength (but not the probability) of synaptic connections from excitatory to non-parvalbumin inhibitory neurons is determined by how similarly tuned pre- and post-synaptic neurons are to sensory stimuli. Second, the synaptic strength in connected pairs of excitatory cortical neurons depends on both the similarity of the stimulus tuning of the two neurons and the similarity in the phase of their responses. Third, as a consequence of the anisotropic mapping from the visual space to cortical space (e.g., a 10° movement in the visual space requires ∼250 μm movement within cortex for elevation, but only ∼140 μm for azimuth) (Schuett, Bonhoeffer and Hübener, 2002; Kalatsky and Stryker, 2003), there will be an asymmetry between neurons preferring vertical vs. horizontal directions of motion that is compensated for by an asymmetric circuit architecture.

Our models use the Brain Modeling ToolKit (BMTK, github.com/AllenInstitute/bmtk) (Gratiy *et al*., 2018) that facilitates building large-scale networks with both NEURON (Hines and Carnevale, 1997) and NEST (Gewaltig and Diesmann, 2007). The model architecture and its outputs (e.g., transmembrane voltage across dendrites and soma, spiking activity) are saved using the standardized SONATA format (github.com/AllenInstitute/sonata, (Dai *et al*., 2019)). All models, code, and meta-data resources are publicly available for download via the Allen Institute for Brain Science’s web portal (brain-map.org/explore/models/mv1-all-layers). As an open public resource, these models will be useful for making direct predictions as well as complementing other experimental and modeling endeavors. The *in vivo* extracellular recordings used for comparison are recorded from a standardized pipeline that is freely available from the Allen Institute (Siegle *et al*., 2019;brain-map.org/explore/circuits/visual-coding-neuropixels).

## Results

### Populating Diverse Cortical Cell Classes With Neuronal Models

Our biophysical and GLIF variants of V1 use the same connectivity graph (i.e., each neuron of ∼230,000 nodes in one variant has an exact counterpart in the other, with the same coordinates, presynaptic sources, and postsynaptic targets). The first step in building this network is to instantiate and distribute neurons in a data-driven manner. The models span a 845 μm radius of cortex (Fig. 1B). For the biophysically detailed variant, the “core” (400 μm radius) is composed of spatially extended neurons, with passive dendrites, surrounded by an annulus of leaky-integrate-and-fire neurons, to avoid boundary artifacts (as described previously, (Arkhipov *et al*., 2018)). In both the GLIF and biophysical variants, the focus of our analysis is on the network within this central core.

Neuron models are reconstructed from slice electrophysiology and are publicly available from the Allen Cell Types Database (Gouwens *et al*., 2018; Teeter *et al*., 2018, celltypes.brain-map.org). Although the most recent transcriptomic and electrophysiological/morphological *in vitro* surveys suggest ∼50-100 neuronal classes in V1 (Tasic *et al*., 2018; Gouwens *et al*., 2019), the currently available neuronal models, connectivity data, and *in vivo* recordings offer lower cell class resolution. Thus, we adopted a coarser classification of 17 classes (Fig. 1A, C).

Specifically, inhibitory neurons are drawn from Htr3a neurons in layer 1 (L1) and from three classes of interneurons in layers 2/3 to 6 - Paravalbumin- (Pvalb), Somatostatin- (Sst), and Htr3a positive cells (Lee *et al*., 2010; Tremblay, Lee and Rudy, 2016). We note that the commonly studied Vasoactive Intestinal Polypeptide (VIP) interneurons are a subclass of Htr3a in L2/3 to L6 (Tremblay, Lee and Rudy, 2016). Since VIP is the subclass most extensively characterized experimentally within the Htr3a class, we resorted to using VIP studies to constrain the Htr3a class in our models (see below). One class of excitatory neurons are each present in L2/3, L4, L5, and L6 (E2/3, E4, E5, and E6).

In total, the two V1 variants contain 17 cell classes, represented by 112 unique individual neuron models for the biophysical and 111 for the GLIF network, copied and distributed in layers according to the best data available (see Methods). Cell densities (including inhibitory subclasses) across layers are estimated from anatomical data (Schüz and Palm, 1989; Lee *et al*., 2010), with an 85%:15% fraction for excitatory and inhibitory neurons. The final network contains 230,924 cells, of which 51,978 are in the core (see Methods).

We determined synaptic connectivity using three design iterations. In the first, discussed immediately below, we determined the feed-forward geniculate input into cortex. In the second iteration, we introduced massive synaptic recurrency within cortex, which depended on the difference in stimulus tuning among pairs of cortical cells. In the third and final step, we refined the recurrent functional connectivity with respect to the stimulus tuning properties as well as the phase difference between responses of the connected cells.

## Thalamic Input To The V1 Models

The Lateral Geniculate Nucleus (LGN) of the thalamus mediates retinal input to V1. We created an LGN module that generates action potentials for arbitrary visual stimuli, as described below.

### Creating LGN Units

The LGN module is composed of spatio-temporally separable filter units (released publicly via the Brain Modeling ToolKit, github.com/AllenInstitute/bmtk) fitted to electrophysiology recordings from mouse LGN (Durand *et al*., 2016). In a substantial elaboration over our previous work (Arkhipov *et al*., 2018), we developed filters for four classes of experimentally observed functional responses (Piscopo *et al*., 2013; Durand *et al*., 2016): sustained ON, sustained OFF, transient OFF, and ON/OFF (the latter is related to the DS/OS class of (Piscopo *et al*., 2013)). These four filter groups are further subdivided according to their maximal response to drifting gratings of different temporal frequencies (TF) (Fig. 2A). We average the experimentally recorded responses for each class to create linear-nonlinear filter models that process any spatio-temporal input and compute a firing rate output (see Methods, example in Fig. 2B). These filters are distributed in visual space according to occurrence ratios of the LGN cell classes (Durand *et al*., 2016), translating any visual stimulus into firing rates. The firing rate output is converted, via a Poisson process, into spike times (see Methods).

**Figure 2:**
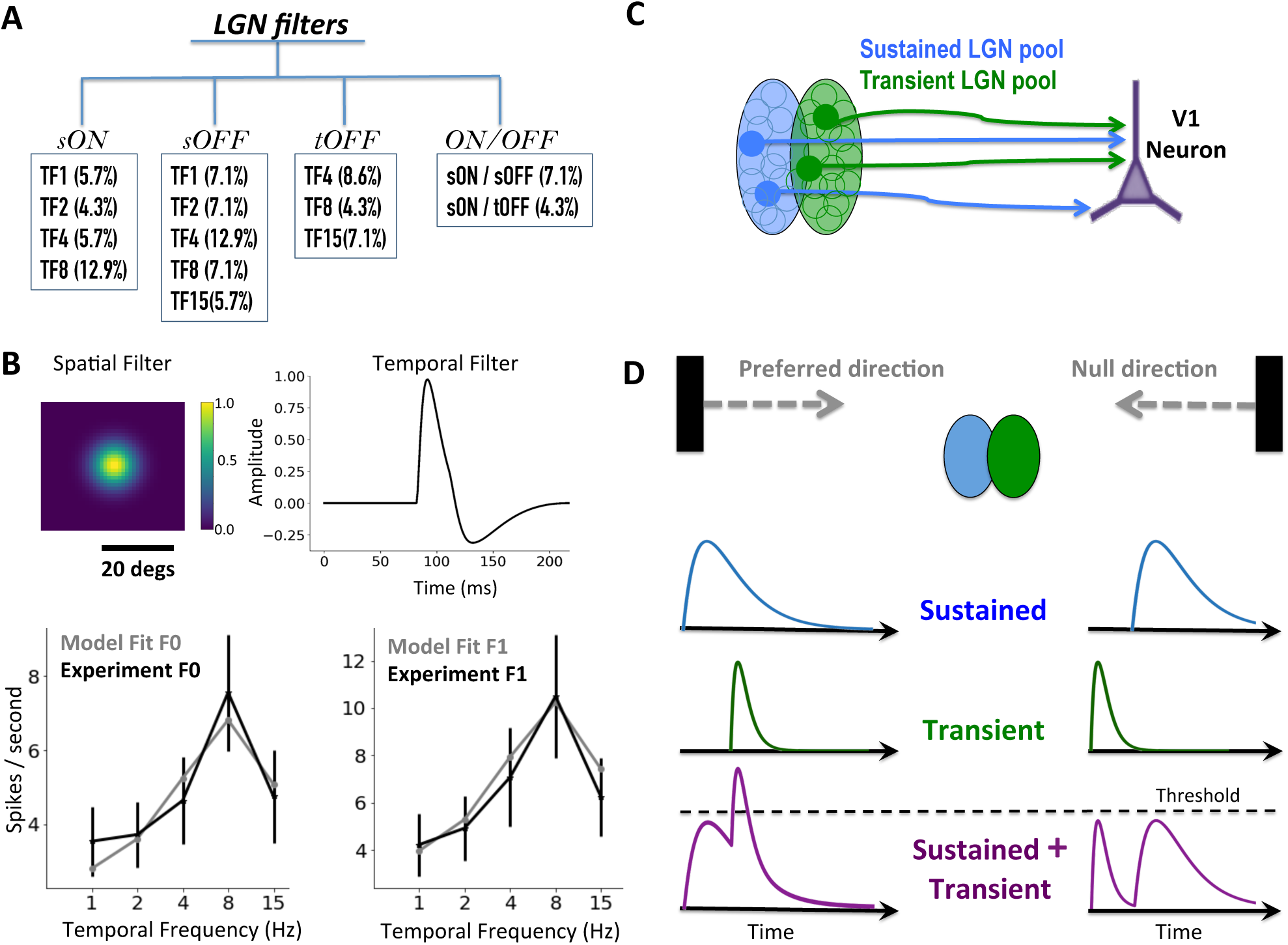
Filter models for enabling arbitrary visual input to be applied via the lateral geniculate nucleus (LGN). **A:** LGN cell classes fit from electrophysiological recordings (Durand *et al*., 2016) using spatiotemporally separable filters. Every major class has sub-classes that respond maximally to specific temporal frequencies (TF). The numbers in parentheses indicate the rate of occurrence in our model. **B:** Example filter for the sON-TF8 class. *Top:* the spatial and temporal components of the filter. *Bottom*: plots of the F0 (cycle averaged mean rate response) and F1 components (modulation of the response at the input stimulus frequency) of the data and the model fit (see Methods) in response to drifting gratings (mean ± s.e.m). **C:** Schematic of a candidate pool of LGN cells, separated into sustained and transient subfields, projecting to a cortical cell with matching retinotopic positions. **D:** Schematic illustrating the direction selectivity mechanism. When a bar moves from left to right (“preferred” direction), the responses from the sustained and transient components overlap and exceed a threshold, while movement in the opposite (“null”) direction prevents overlapping responses.

### Direction Selective Input Into V1 Cells

Our main emphasis is on how the structure of cortical circuits is reflected in their *in vivo* function. In visual cortex, a major functional property is direction selectivity (Niell and Stryker, 2008; Durand *et al*., 2016); its genesis is a central question in the field. We sought to recapitulate physiological levels of direction selectivity and, in doing so, investigate the underlying structure, starting from feedforward thalamocortical inputs, subsequently adding recurrent connectivity (see below). Direction selectivity is only one of several other metrics that we evaluated. Our freely available models permit one to test many other metrics and ask many more questions about structure-function relationships in cortical circuits.

Although some direction-selectivity is observed in the LGN (Marshel *et al*., 2012; Piscopo *et al*., 2013; Scholl *et al*., 2013; Zhao *et al*., 2013; Sun *et al*., 2016), recent work indicates that direction selectivity is produced *de novo* in V1 from convergence of spatio-temporally asymmetric LGN inputs (Lien and Scanziani, 2018). Based on this, we assume that LGN innervation into V1 neurons has two subfields, one with slow (sustained) and the other with fast (transient) kinetics (Fig. 2C). These produce an asymmetry in responses to opposite directions of motion (Fig. 2D). A simplified theoretical framework demonstrates (see Methods, Fig. S1) that sufficiently high orientation and direction selectivity indices (OSI and DSI; see Methods), which are the metrics of how selective individual neurons are for orientation and direction of stimuli, can be achieved with such input subfields (Lien and Scanziani, 2013, 2018). Interestingly, this analytic treatment predicts reversal of preferred direction as the spatial frequency of grating increases, which we confirmed experimentally to be a ubiquitous phenomenon in the mouse visual system (Billeh *et. al* 2019). This mechanism has analogy to aliasing found in the fly (Hassenstein and Reichardt, 1956; Barlow and Levick, 1965; Van Santen and Sperling, 1984; Borst and Egelhaaf, 1989) and parallels the OFF pathway motion detection system in fly T5 neurons (Serbe *et al*., 2016; Arenz *et al*., 2017).

### Creating And Testing Thalamocortical Connectivity

We instantiated individual filters to represent the diverse LGN responses (Fig. 2A), placing 17,400 LGN units in visual space. LGN axons project to all layers of V1 (Kloc and Maffei, 2014; Morgenstern, Bourg and Petreanu, 2016), selectively innervating excitatory neurons and Pvalb interneurons in L2/3-L6, as well as non-Pvalb interneurons in L1 (Ji *et al*., 2015). We targeted LGN inputs to V1 neuron classes accordingly and then established connections to individual neurons using the following three-step procedure (see Methods).

The first step selects the LGN units projecting to a particular V1 neuron, leveraging the fact that the spatiotemporally asymmetric architecture yields direction and orientation selectivity (Lien and Scanziani, 2013, 2018). For each V1 neuron, we determined the visual center, size, and directionality (a pre-assigned preferred angle of stimulus motion) of elliptical subfields from which LGN filters will be sampled, according to the neuron’s class and position in the cortical plane (Fig. 3A). We then identified LGN receptive fields (RFs, parameterized during filter construction) that overlap with these elliptical subfields of the V1 neuron. One subfield always samples from transient OFF LGN filters and the other from sustained ON or OFF (see Methods).

**Figure 3:**
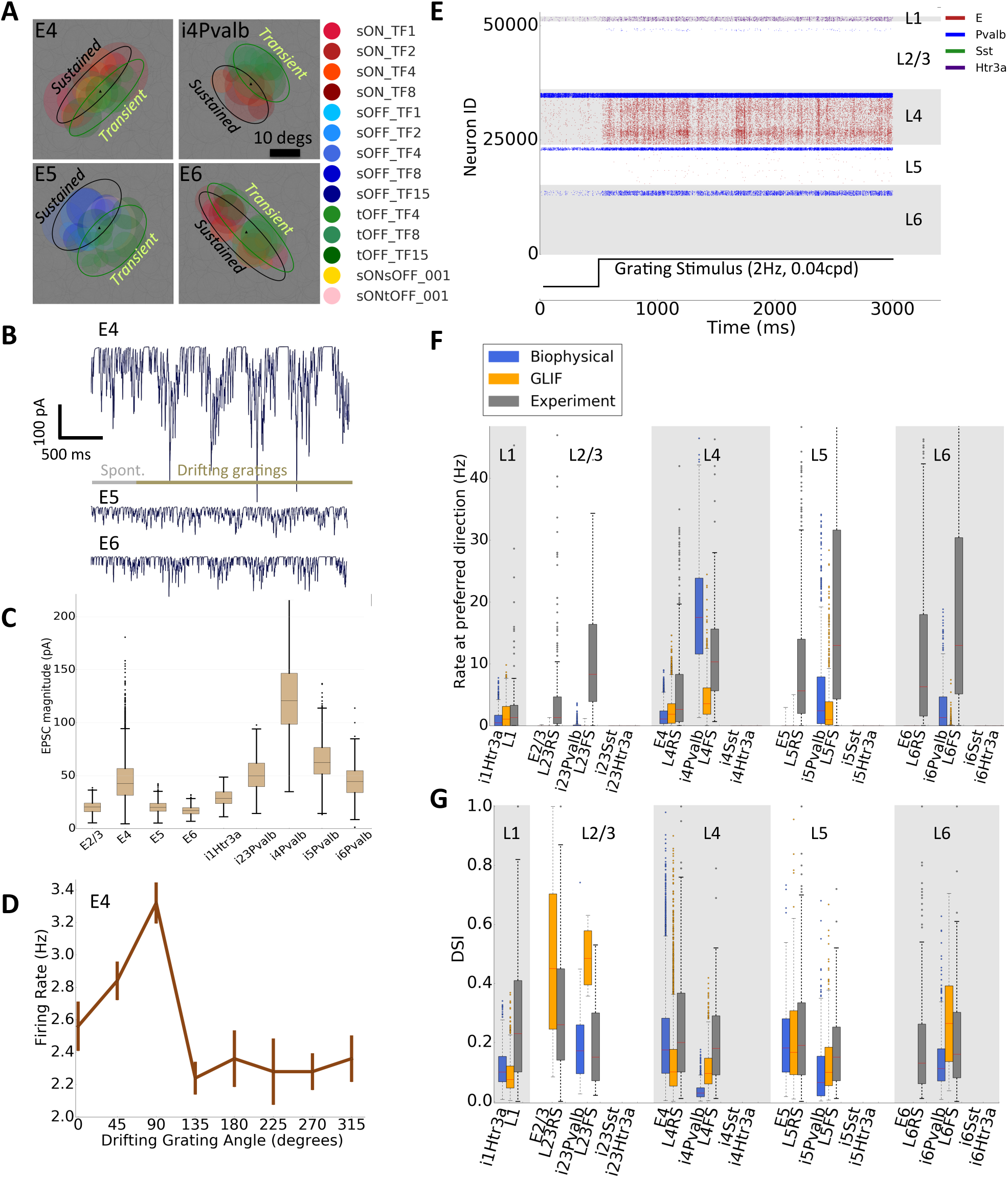
Thalamic inputs to the cortical models. **A:** LGN filters connecting to four different cortical (V1) neurons. Black triangles and colored circles indicate the centers of the receptive fields of V1 and presynaptic LGN neurons. Gray circles indicate all other LGN filters. The elliptical subfields used to select the projecting LGN filters are shown. **B-G**: Responses of the biophysical V1 model to LGN input without any intracortical connections nor background activity. **B:** Postsynaptic currents in V1 neurons responding to 500 ms of gray screen followed by a drifting grating. The mean current is matched to experimental measurements and is largest for layer 4 neurons. **C:** Boxplots of postsynaptic currents for every neuron class (for preferred drifting grating), after matching to target values (median indicated with a bar; boxes span 25^th^ till 75^th^ percentile and whiskers extend a maximum of 1.5 the interquartile range). **D:** Example tuning curve of an E4 neuron (the neuron’s firing rate in response to visual stimulation with a grating drifting in different directions, from 0° to 315°, shown for each direction as mean ± s.e.m over 10 trials). **E:** Example raster plot, same stimulus as in B. Neuron classes with large EPSC current values (boxplots in C) exhibit substantial spiking activity. **F:** Boxplots characterizing firing rates of neuronal classes. For reference, experimental data from *in vivo* extracellular electrophysiology recordings from awake mice (i.e., fully connected cortical circuit) are shown. **G:** Characterization of the direction selectivity index (DSI) from responsive neurons (see Methods). Some DSI values are high as these simulations are purely feedforward and thus exhibit low firing rates.

The second step, only applicable to the biophysical model, determines the number and placement of synapses on V1 neurons, using data on LGN axonal density in V1 (Morgenstern, Bourg and Petreanu, 2016) and estimates of synapse numbers per neuron (Schoonover et al., 2014; Bopp et al., 2017). The effect of dendritic placement on the somatic charge conductance is shown in Fig. S2.

The third and final step establishes the strength of the thalamocortical synapses, constrained by experimental current measurements (Lien and Scanziani, 2013; Ji et al., 2015). The strength is scaled to match the target mean current (Fig. 3B, C) in response to a drifting grating (see Methods). Layer 4 is the main primary input target of the thalamocortical projections, and therefore the currents are largest in this layer (Fig. 3B, C).

To test the outcome of this procedure, we carried out simulations of the entire network without recurrent connections using drifting gratings. Individual neurons are direction selective (Fig. 3D), consistent with experimental measurements of LGN input currents (Lien and Scanziani, 2013, 2018). At the network level (example raster in Fig. 3E), the average firing rates, DSI, and OSI due to LGN-only input are calculated (Figs. 3F, 3G, and S3 respectively). For reference, data from *in vivo* extracellular Neuropixels recordings from awake mice within a standard pipeline (Siegle *et al*., 2019) are included in Fig. 3F (data available by the Allen Institute, brain-map.org/explore/circuits/visual-coding-neuropixels) and are used throughout the manuscript as a benchmark (examples in Fig. S4). Note that experimental data are robustly classified into regular-spiking (RS) and fast-spiking groups (FS), roughly corresponding to excitatory and Pvalb inhibitory neurons (although small contributions from non-Pvalb inhibitory neurons are likely present in both groups). Hence, here and throughout the Results section, we compare model excitatory and Pvalb neurons with experimental RS and FS cells, respectively. Further, we use gray throughout the manuscript to indicate experimental data, blue for the biophysical network model, and orange for the GLIF network model unless otherwise stated.

Finally we define a similarity score, **S**, between distributions of a metric of interest, to compare the population of excitatory and Pvalb neurons in experiments and models (one minus the Kolmogorov–Smirnov distance, see Methods). If the distribution in the simulated population is close to the experimental one, the similarity measure will be close to unity; **S** close to zero indicates quite different distributions (Fig. S5). As expected, in the absence of intra-cortical amplification, **S** is low for firing rates (E-biophysical = 0.18, E-GLIF = 0.19, Pvalb-biophysical = 0.62, Pvalb-GLIF = 0.37), for OSI (E-biophysical = 0.24, E-GLIF = 0.25, Pvalb-biophysical = 0.49, Pvalb-GLIF = 0.57), and for DSI (E-biophysical = 0.24, E-GLIF = 0.24, Pvalb-biophysical = 0.58, Pvalb-GLIF = 0.63). We also note that the two model resolutions compare well to one another (e.g., **S** values: E-rates = 0.96, E-OSI = 0.95, E-DSI = 0.96).

Finally, a background pool, mimicking the influence of the rest of the brain on the modeled region provides inputs from a single Poisson source firing at a constant rate of 1 kHz to all V1 cells. The weights of this background were adjusted with the recurrent connectivity in place, to ensure that the baseline firing rates of all neurons match experiments (see below). While more sophisticated models of background can be implemented (e.g., Arkhipov *et al*., 2018), this simple one sufficed for our purpose.

## Creating the recurrent connectivity in the V1 network

We now turn to the considerably more complex problem of determining cortico-cortical connections.

Cortical circuits feature extensive recurrent connections which amplify thalamocortical inputs and shape cortical computations (Douglas, Martin and Whitteridge, 1989; Douglas *et al*., 1995; Douglas and Martin, 2007; Lien and Scanziani, 2013; Arkhipov *et al*., 2018). Despite many studies (e.g., (Cauli *et al*., 1997; Dantzker and Callaway, 2000; Beierlein and Connors, 2002; Thomson *et al*., 2002; Beierlein, Gibson and Connors, 2003; Mercer *et al*., 2005; Song *et al*., 2005; West *et al*., 2005; Yoshimura, Dantzker and Callaway, 2005; Lefort *et al*., 2009; Hofer *et al*., 2011; Ko *et al*., 2011; Levy and Reyes, 2012; Olsen *et al*., 2012; Pfeffer *et al*., 2013; Vélez-Fort *et al*., 2014; Bortone, Olsen and Scanziani, 2014; Cossell *et al*., 2015; Jiang *et al*., 2015)), data on the exact patterns and magnitude of V1 recurrent connectivity remains sparse, and no resource exists that comprehensively characterizes all connections under standardized conditions. We set out to construct recurrent connections in a data-driven manner via extensive curation of the literature supplemented by Allen Institute data (Seeman *et al*., 2018) when available. This resulted in four key resources (Fig. 4) containing estimates of (1) connection probability, (2) synaptic amplitude or strengths, (3) axonal delays, and (4) dendritic targeting of synapses. These resources are provided to the community (brain-map.org/explore/models/mv1-all-layers) with every estimate and assumption documented in interactive files. Our cortical network contains specific instantiations of these connectivity rules.

**Figure 4:**
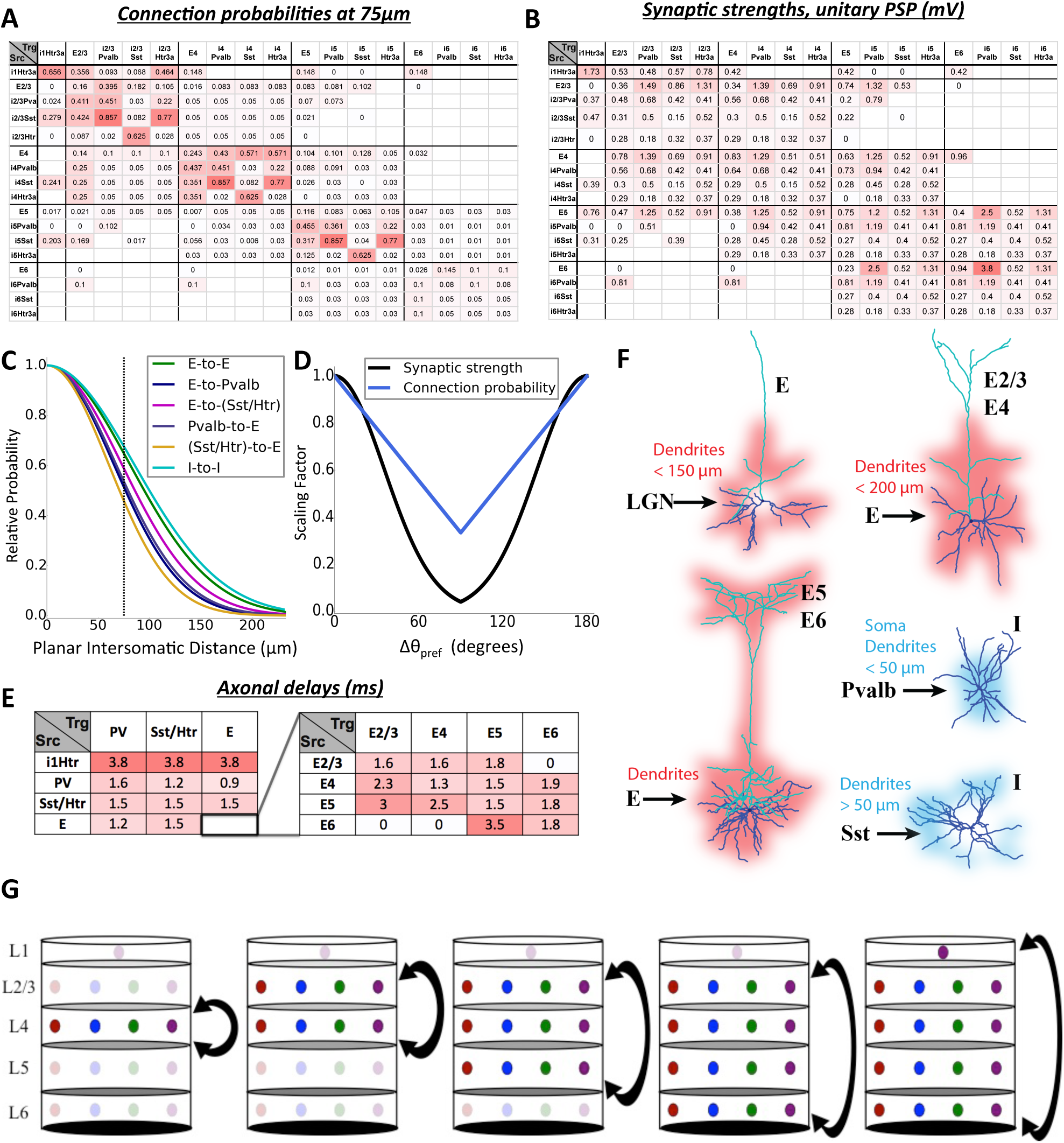
Summary of recurrent connectivity rules used for both variants. **A:** Probability of connection at an intersomatic distance of 75 μm. **B:** Strength of connections (somatic unitary post-synaptic potential (PSP)). **C:** The distance-dependent connection probability profiles for different classes of connections. **D:** The functional rules for connection probability (applied only to E-to-E connections) and synaptic strengths (applied to all connection classes) as a function of the difference in preferred direction angle between the source and target neurons. **E:** Axonal delays for connections between classes. **F:** Example schematics of dendritic targeting rules. For detailed descriptions, see Methods. **G:** Schematic illustrating the layer-by-layer optimization procedure after incorporating all known data for recurrent connections. In the first stage, all recurrent connections except those within L4 were set to zero, and the L4 weights were optimized. Then, L2/3 connections (within layer and between L2/3 and L4) were added and optimized. The procedure was repeated, adding one layer at a time, until all layers were connected and optimized. Intermediate simulation outputs for the different stages of optimization are shown in Fig. S6.

Unfortunately, data do not exist for many connection classes in mouse V1; therefore we used other sources of information in the following order of preference as a guiding principle: mouse visual cortex, followed by mouse non-visual or rat visual cortex, then rat non-visual cortical measurements. Additional entries were filled using assumptions of similarity and/or the rat somatosensory cortex model (Markram *et al*., 2015; Reimann *et al*., 2015). 89 out of the total 289 entries remained undetermined (empty cells in Fig. 4A, B) and were set to zero due to lack of data (see Methods).

Fig. 4A reports connection probability values at 75 μm planar intersomatic distance, used as parameters for Gaussian distance-dependent connectivity rules for different source-target class pairs (Fig. 4C). Excitatory-to-excitatory (E-to-E) connections in L2/3 of mouse V1 also exhibit “like-to-like” preferences (Ko *et al*., 2011; Cossell *et al*., 2015; Wertz *et al*., 2015; Lee *et al*., 2016) – that is, cells preferring similar stimuli are preferentially connected. We here assume that such like-to-like rules are ubiquitous among all E-to-E connections, both within and across layers. These rules are illustrated in Fig. 4D (see Methods), based on the preferred direction of motion angle assigned (*θ_pref_*) to each neuron. No such rules were applied for E-to-I, I-to-E, and I-to-I connection probabilities, following experimental observations (Bock *et al*., 2011; Fino and Yuste, 2011; Packer and Yuste, 2011; Znamenskiy *et al*., 2018).

Recent experiments indicate that, besides connection probability, the amplitude (strength) of E-to-E synaptic connections in L2/3 also exhibit a like-to-like dependence (Cossell *et al*., 2015; Lee *et al*., 2016). In earlier work, we found these to be even more important for response tuning than connection probability rules (Schaub *et al*., 2015; Arkhipov *et al*., 2018). A similar like-to-like rule for synaptic strength (but not connection probability) has been reported for I-to-E connections (Znamenskiy *et al*., 2018). Thus, we assume that all synaptic strength classes (Fig. 4B) are modulated by such a rule (Fig. 4D). At this point, all like-to-like connection probability and synaptic strength profiles were symmetric with respect to the opposite preferred directions (i.e., orientation-dependent but not direction-dependent).

Notably, some of the first predictions from our models came from this data-driven building stage. One important rationale for imposing the like-to-like synaptic weights rule for all connection classes is that the Sst and Htr3a classes receive little to no LGN input (Fig. 3; (Ji *et al*., 2015)), yet exhibit orientation and direction tuning (Liu *et al*., 2009; Kerlin *et al*., 2010; Ma *et al*., 2010). We assumed that these classes become tuned due to like-to-like inputs from excitatory neurons, and, indeed, our simulations implementing these rules exhibit substantial orientation and direction selectivity for these interneuron classes. We confirm this prediction by removing the like-to-like synaptic weight rule to Sst and Htr3a neurons in our final model: indeed, both cell classes lose their orientation and direction selectivity properties (see below).

The third resource contains synaptic delays between different neuronal classes. Given that measurements of these properties were particularly sparse, our final table is of coarser resolution (Fig. 4E). The fourth resource, applicable to the biophysical model only, is a set of dendritic targeting rules for each connection class (examples illustrated in Fig. 4F). Experimental data for this (typically, from electron microscopy) are only available for a relatively small number of scenarios, and we used what was available from internal data and the literature (see Methods).

### Optimization of synaptic weights

Although our data-driven approach systematically integrates a large body of available data, these data are still incomplete and are obtained under disparate conditions. It is therefore not surprising that after construction, our models need to be tuned to obtain physiologically realistic spiking patterns and avoid run-away excitation or epileptic-like activity. While efficient optimization methods for recurrent spiking networks have been described (e.g., (Sussillo and Abbott, 2009; Nicola and Clopath, 2017)), their performance has not yet achieved the level required for optimization of the computationally expensive and highly heterogeneous networks we constructed. We therefore use a heuristic optimization approach with identical criteria applied to biophysical and GLIF variants.

Following (Arkhipov *et al*., 2018), we employed three criteria: both (i) spontaneous firing rates as well as (ii) peak firing rates in response to a single trial of a drifting grating (0.5 s long) should match experimental data, and (iii) the models should not exhibit epileptic activity. The optimization applied to synaptic weights only, via grid searches along weights of connections between neuronal classes, using uniform scaling of the selected weight class. The LGN-to-L4 weights were fixed as they were matched directly to experimental recordings *in vivo* (Lien and Scanziani, 2013) (Fig. 3), whereas the net current inputs from LGN to other layers could vary (within strict bounds) since the corresponding experimental data were obtained *in vitro* (Ji *et al*., 2015). Optimizing a fully recurrent network in one step was very challenging; instead, we followed a stepwise, layer-by-layer procedure (Fig. 4G). We first optimized the recurrent weights within L4, with all recurrent connections outside L4 removed (Fig. S6). Then we added L2/3 recurrent connections and optimized the weights in both L4 and L2/3. This approach was repeated by adding L5, then L6, and finally L1 (Figs. 4G, S6; see Methods for details). The order of adding layers was selected based on the canonical cortical microcircuit (Douglas, Martin and Whitteridge, 1989; Douglas *et al*., 1995; Douglas and Martin, 2007). We expect that using such biological insights to guide optimizations of large-scale biological models, perhaps as a strategy accompanying algorithmic methods, may increase the speed and likelihood of convergence.

After optimization, a typical response to a drifting grating exhibits irregular activity, with the strongest spiking among neurons tuned to the drifting grating’s direction of motion (Fig. 5A). The firing rates for both V1 variants across all neuronal classes are similar to those measured *in vivo* (Fig. 5B, **S** values: E-biophysical = 0.76, E-GLIF = 0.75, Pvalb-biophysical = 0.85, Pvalb-GLIF = 0.83). Excitatory neurons show improved, relative to LGN only simulations (Fig. 3G), yet unsatisfactory orientation tuning (Fig. S7, **S** values: E-biophysical = 0.50, E-GLIF = 0.56, Pvalb-biophysical = 0.27, Pvalb-GLIF = 0.42). Similarly, the direction selectivity match is also poor, particularly for Pvalb interneuons (Figs. 5C, 5D, **S** values: E-biophysical 0.53, E-GLIF = 0.54, Pvalb-biophysical = 0.29, Pvalb-GLIF = 0.32). Given that all neuron classes fire in these recurrently connected networks, we compare the models to time-based dynamic metrics. We find our models match well with experimental data for signal correlations (pairwise correlation of mean responses to the drifting grating directions), noise correlations (spike-count correlations to repeated drifting grating presentations), and correlations of signal and noise correlations (correlation of the signal and noise correlations for a group of neurons), discussed in Fig. S8 (see Methods). Finally, we once again observe strong similarity between both variants (see Discussion; **S** values: E-rates = 0.90, E-OSI = 0.89, E-DSI = 0.95).

**Figure 5:**
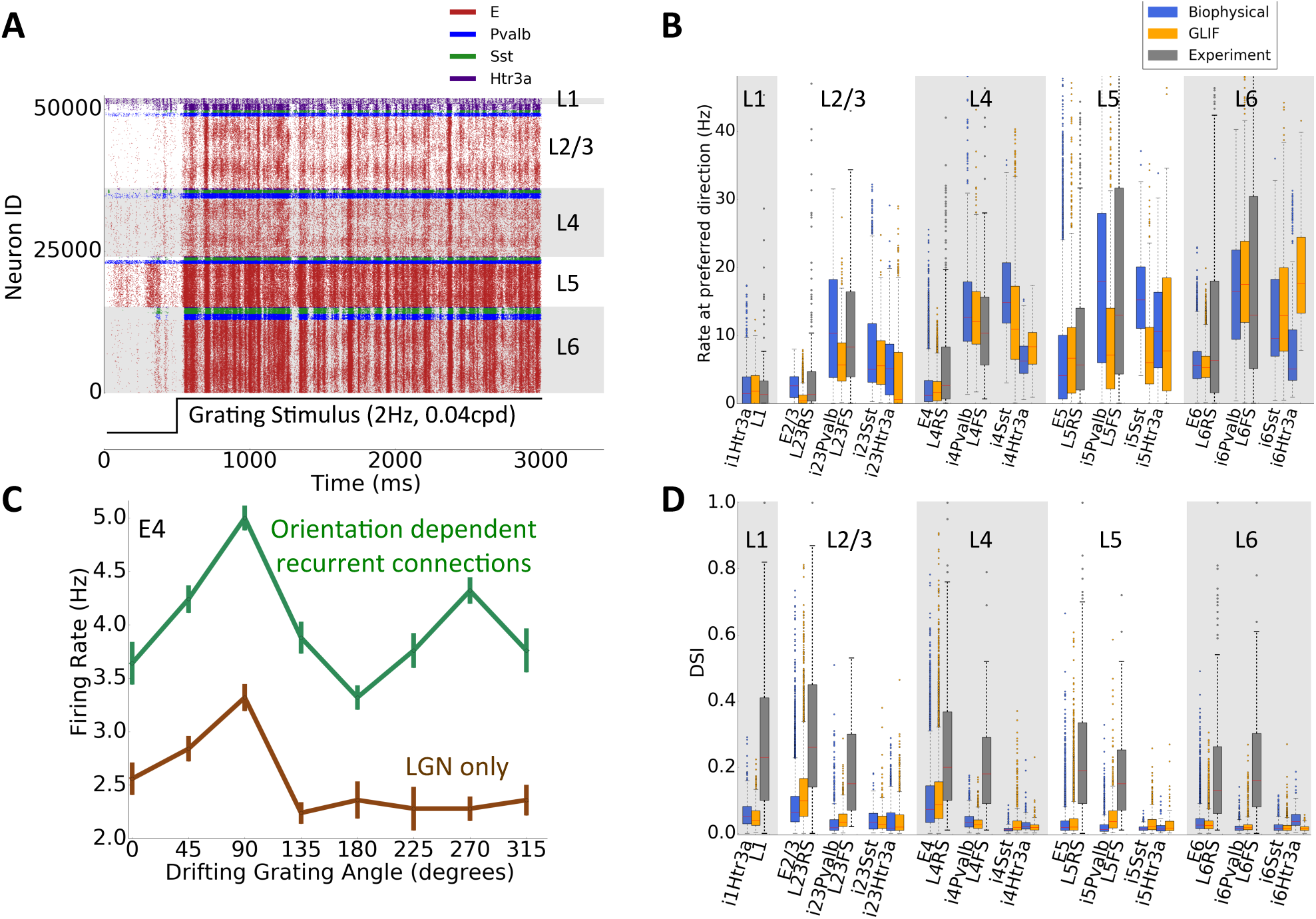
Initial simulation results from the biophysical and GLIF recurrent V1 models. **A:** Raster plot in response to a drifting grating (biophysical model). Within each cell class, the cell IDs are sorted according to the cells’ preferred angles. **B:** Peak firing rate boxplots for both V1 models and *in vivo* extracellular electrophysiology recordings. **C:** Example tuning curves (mean ± s.e.m) for both the biophysical recurrent model and LGN-only model (same neuron as Fig. 3D). **D:** DSI boxplots for both V1 variants and *in vivo* measurements.

Although these results are an improvement relative to the simulations without recurrent connections (Fig. 3), they fail at orientation and direction selectivity. While the structure of our network was constrained by data relatively well at the level of cell classes, existing data provide much fewer constraints on the functional rules of individual recurrent connections, which relate tuning of neurons to their connectivity and synaptic properties. We therefore reasoned that these functional rules require further adjustment in the models to improve their performance with respect to orientation and direction selectivity and explored this hypothesis as described below.

### Refined synaptic functional connections amplify direction selectivity

Up to this point, the recurrent like-to-like connectivity rules in our models were “orientation-based”, *i.e*., the probability and weights were symmetric with respect to Δθ=90°, where Δθ is the difference between the preferred angles of the two neurons (Fig. 4D). In other words, for two neurons that prefer opposite directions (Δθ=180°), the probability of them being connected and their synapse strength were the same as for two neurons that prefer the same direction (Δθ=0°). This can be contrasted with “direction-based” asymmetric rules, where a pair of neurons preferring opposite directions of motion are treated differently from a pair preferring the same direction (Fig. 6A). We reasoned that low levels of direction selectivity in our V1 models are due to the absence of such direction-based rules, since the orientation-based rules enhance neurons’ responses to their anti-preferred direction due to inputs from the oppositely tuned neurons. However, the models are also grounded in data, which show symmetric, orientation-based like-to-like rules for probability of E-to-E connections and no like-to-like rules for I-to-E connections (Fino and Yuste, 2011; Ko *et al*., 2011; Packer and Yuste, 2011; Lee *et al*., 2016; Znamenskiy *et al*., 2018) (although the data is mostly limited to connection classes in L2/3). We assumed that all E-to-E connection probabilities obey the orientation-based rule. Therefore, the only remaining flexibility is in the functional rules specifying synaptic weights, rather than connection probabilities, of any connections formed.

**Figure 6:**
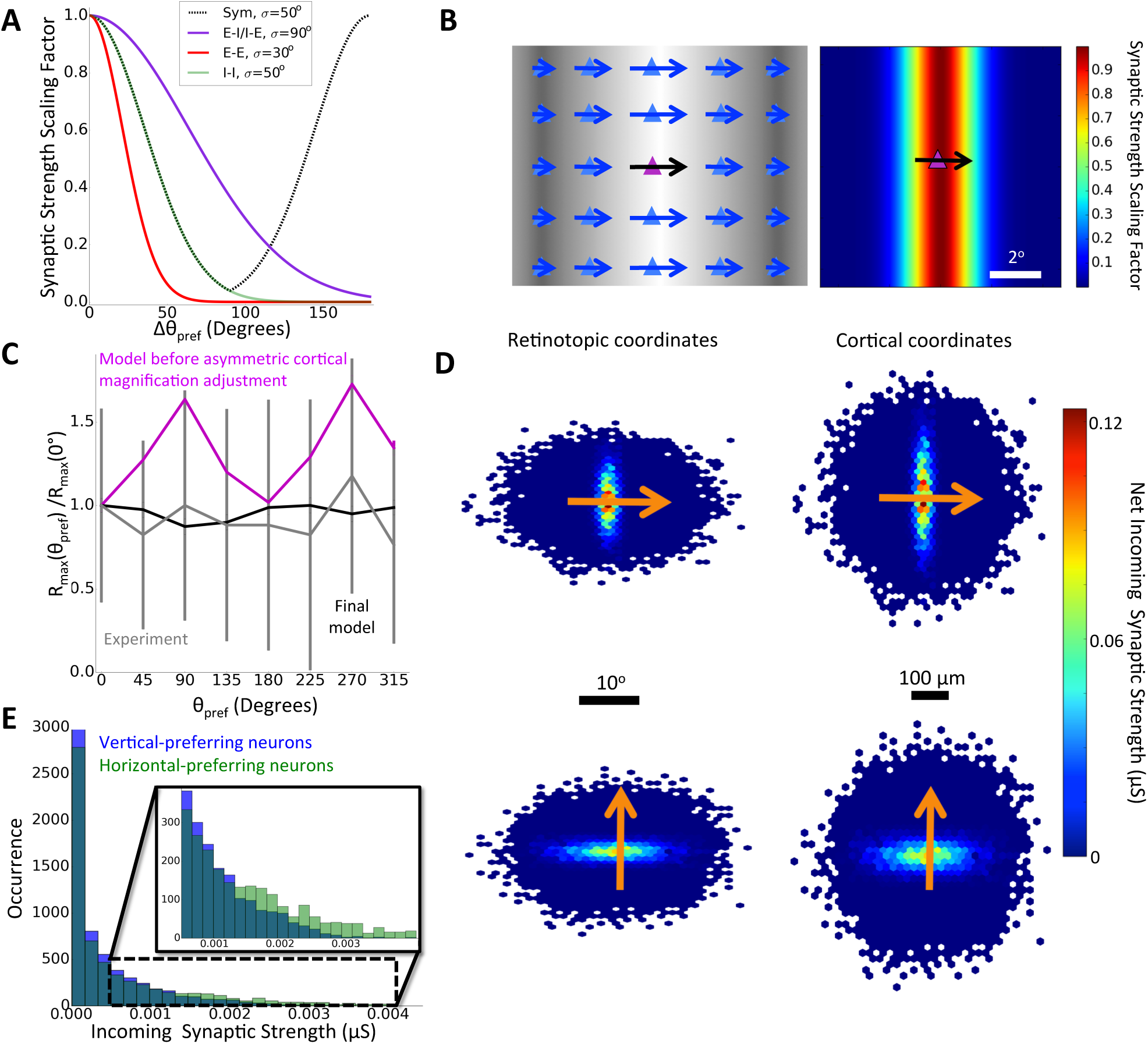
Refined synaptic functional connections. **A:** The original symmetric or “orientation-based” (dotted black, “Sym”; Fig. 4D) and the refined, asymmetric or “direction-based” (colors) synaptic strength profiles as a function of the difference between the preferred angles in two connected neurons. The like-to-like rule for E-to-E connection probabilities remains orientation-based (Fig. 4D) and no like-to-like rules are applied to other connection probabilities. **B:** The phase-based rule for synaptic strengths of E-to-E connections. *Left*: Schematic example of neurons preferring 0° direction, as they respond to a 0° drifting grating (background shows phase alignment with the drifting grating). Arrow lengths indicate magnitude of response. Neurons aligned vertically with the center neuron have a matching phase. *Right*: stronger weights are assigned to phase-matched than phase-unmatched neurons (the heat-map illustrates the scaling factor applied in the models). **C:** Log firing rates of excitatory neurons in response to their preferred drifting grating direction (median ± s.e.m), for the biophysical model. Applying the rules from (A) and (B) results in a firing rate bias for vertical- vs. horizontal-preferring neurons due to differential cortical magnification (magenta); the bias is not observed experimentally (gray). The bias disappears after additional direction-dependent scaling is applied to synaptic weights according to the target neuron’s assigned preferred angle (black). **D:** Net synaptic inputs for horizontal- and vertical-preferring E4 biophysical neurons (rules in (A) and (B) and the additional direction-dependent scaling), in retinotopic (left) and cortical (right) coordinates (averages over 100 neurons after aligning their centers). The connection rules are also included here (distance dependence and “orientation dependent” like-to-like rules). The distance dependence determines the length of these stripes. **E:** Histogram of incoming synaptic weights onto E4 neurons based on their preferred orientation. Horizontal-preferring neurons have a heavier tail than vertical-preferring neurons.

Less is known about functional rules for synaptic strength. Available data from L2/3 (Cossell *et al*., 2015; Znamenskiy *et al*., 2018) indicate that synaptic amplitude correlates with similarity of responses, for both E- to-E and I-to-E connected pairs. However, similarity of preferred direction alone is a poor predictor of synaptic strength for E-to-E connections, whereas similarity of receptive fields (ON-OFF overlap) is a better predictor (Cossell *et al*., 2015). Furthermore, *in vivo* patch-clamp measurements in L4 indicate that excitatory neurons responding to a drifting grating in phase with each other are preferentially connected (Lien and Scanziani, 2013). Motivated by these observations, we introduce two modifications to the synaptic strength rules: (1) a direction-of-motion-based like-to-like Gaussian profile applied to all connection classes (Fig. 6A) and (2), for the E-to-E classes only, a decrease of the synaptic strength with distance in retinotopic visual space between the source and target neurons, projected on the target neuron’s preferred direction (Fig. 6B). No other distance dependency in synaptic strength was introduced. Rule (2) confines the sources of sufficiently strong connections to a stripe perpendicular to the target neuron’s preferred direction, biasing the inputs to come primarily from neurons that respond in phase with the target neuron when stimulated by a drifting grating or a moving edge (Fig. 6B). These assumptions are consistent with theory based on optimal Bayesian synaptic connectivity for integrating visual stimuli (Iyer and Mihalas, 2017). We incorporated rules (1) and (2) at the synaptic strength level while the connection probabilities are governed by a distance dependent rule and an “orientation dependent” like-to-like connectivity rule.

We tested 8 specific choices of rules (1) and (2), sampling multiple selections of parameters for each choice (over 100 variants in total), primarily employing the GLIF V1 model for this purpose (Fig. S9), before converging on a final set (Fig. 6A). With a sufficiently narrow Gaussian curve characterizing the direction-based dependence on Δθ (Fig. 6A), substantial improvement in the levels of DSI are obtained across all layers (Fig. S10). This allows us to predict that like-to-like rules (1) and (2) above may apply across all layers in the mouse V1, potentially with cell-class specific parameters – in fact, in our models we use relatively narrow rule (1) profiles for E-to-E connections, since excitatory populations typically exhibit high DSIs, and wider profiles for other connections (Fig. 6A). Given that multiple different values of parameters for rules (1) and (2) result in networks with robust levels of direction selectivity, we cannot reliably choose a single “optimal” parameter set. The set (Fig. 6A) we use for subsequent simulations should be considered a representative example among possible solutions. In the absence of direct experimental measurements, we simply note that application of rules (1) and (2) with sufficiently narrow profiles (e.g., a Gaussian with standard deviation of 30° for rule (1) in E-to-E connections) enables amplification of direction selectivity by the recurrent connections, consistent with available data.

In testing the connectivity, we notice that rules (1) and (2) are not sufficient by themselves as they introduce a firing rate bias depending on the neuron’s preferred direction of motion. Vertical-preferring neurons exhibit higher peak firing rates than do horizontal-preferring neurons, but such a bias is *not* present in experimental data (Fig. 6C). The root cause of this is the experimentally observed asymmetric retinotopic magnification mapping in cortex (Schuett, Bonhoeffer and Hübener, 2002; Kalatsky and Stryker, 2003), which is implemented in our models (see Methods). Specifically, moving along the horizontal direction in the cortical retinotopic map (azimuth) by 100 μm corresponds to ∼7° in the visual space, whereas along the vertical direction (elevation) 100 μm corresponds to ∼4°. Consequently, the stripe from rule (2) (Fig. 6B) is wider in cortical space for vertical than for horizontal preferring neurons, thus providing stronger net inputs from presynaptic V1 neurons (Fig. S11). Since such a firing rate bias is not empirically observed (Fig. 6C), some mechanisms must adjust for the horizontal-vertical mismatch of translating retinotopy to connectivity. Multiple mechanisms are plausible, including, e.g., different distance dependence of connectivity rules for vertical- vs. horizontal-preferring neurons, different strengths of LGN inputs, or different strengths of recurrent connections. We implement the latter, in a simple linear fashion where horizontal-preferring target neurons receive synapses scaled by 0.5×(7+4)/4=1.38 and vertical neurons scaled by 0.5×(7+4)/7=0.79, with a linear interpolation in-between (see Methods). This approach fixes the firing rate bias and synaptic weight bias (Figs. 6C, S11).

In the final model, horizontal- and vertical-preferring cells receive, on average, equal excitatory synaptic input, sourced from the same size of stripes in retinotopic space, but different widths in physical, cortical space (the stripe for horizontal-preferring cells is almost half the width of the stripe for vertical-preferring cells; Fig. 6D – note also the distance dependence contribution that results in a finite stripe length relative to Fig. 6B). As a consequence (Fig. 6E), the distribution of incoming excitatory weights has a heavier tail for horizontal-than vertical-preferring neurons, an observation that could be tested in the future.

With these finalized models, we simulated drifting gratings (Fig. 7A, B; **S** scores for firing rate: E-biophysical = 0.72, E-GLIF = 0.72, Pvalb-biophysical = 0.80, Pvalb-GLIF = 0.76). Note the emergence of horizontal patches of excitatory neurons (Fig. 7A) due to pronounced direction-selectivity not previously present (Fig. 5A). For excitatory cells, the OSI distributions approximately match experimental recordings (Fig. S12; **S** scores: E-biophysical = 0.75, E-GLIF = 0.72, Pvalb-biophysical = 0.33, Pvalb-GLIF = 0.44), indicating that the new direction- and phase-based rules are not detrimental to orientation selectivity. Most importantly, the match of DSI to experimental values (Fig. 7C, D; **S** scores: E-biophysical = 0.92, E-GLIF = 0.87, Pvalb-biophysical = 0.71, Pvalb-GLIF = 0.83) is much improved for all cell classes, compared to the models with purely orientation-based rules (Fig. 5C, D). The Sst and Htr3a interneurons showed near-zero DSI in Fig. 5D, but now exhibit DSIs equal or higher than those of Pvalb interneurons, consistent with published observations (Kerlin *et al*., 2010; Ma *et al*., 2010). Thus, these new rules successfully enable direction selectivity in distinct populations of neurons, while obeying diverse empirical constraints.

**Figure 7:**
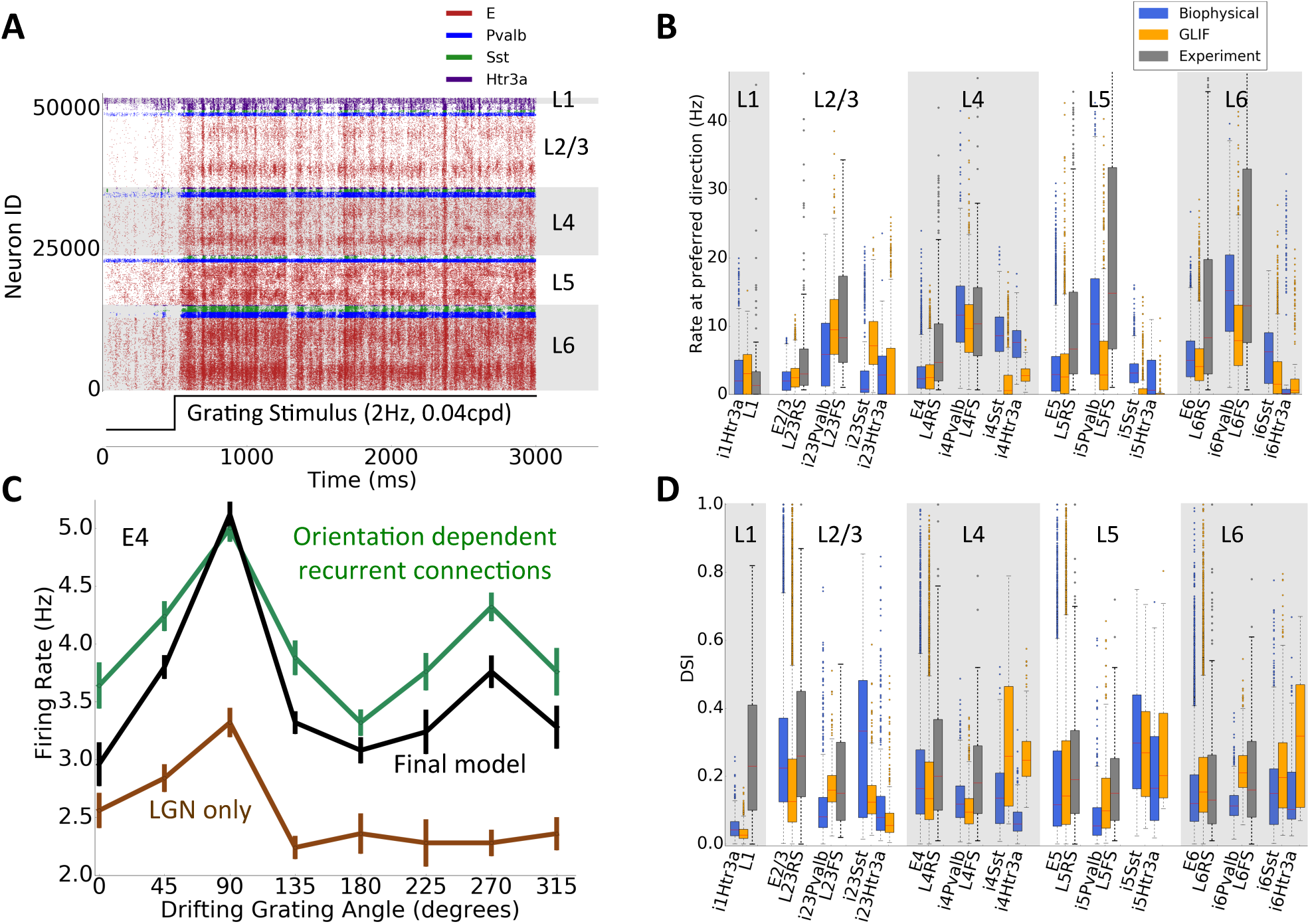
Simulated responses to drifting gratings for the final cortical recurrent connectivity rules (from Fig. 6). **A:** Raster plot in response to a drifting grating (note the horizontal stripes corresponding to strong responses of the cells that prefer the direction of the grating; neuron IDs are sorted within each class by the preferred angle). **B:** Peak firing rate boxplots compared to *in vivo* recordings. **C:** Example tuning curves (mean ± s.e.m) for an E4 neuron for the final rules, in comparison to purely orientation-based rules (Fig. 5C), and no recurrent connections (LGN only of Fig. 3D). **D:** DSI boxplots for the final V1 models and *in vivo* recordings.

Both variants maintain strong similarity with one another (see Discussion; **S** values: E-rates = 0.97, E-OSI = 0.93, E-DSI = 0.95; Table S1). We also calculated the pairwise similarity score between the twenty mice used for our experimental data and found a median similarity score (between animals) for each of the three metrics reported here to be in the range [0.81, 0.84] for excitatory (regular-spiking) neurons and [0.64, 0.66] between Pvalb (fast-spiking) neurons, providing upper limits for matches with experimental data given the variability between animals; as shown above, performance of our final models is close to these limits in most cases. Finally, our dynamics-based metrics still maintain a strong match compared to experimental data (Fig. S13).

We confirmed our predicted need of like-to-like synaptic weight rules for E-to-Sst and E-to-Htr3a neurons (Fig. S14). Indeed when this rule is replaced with uniform synaptic weights, independent of the source-target tuning angle difference, both Sst and Htr3a neurons lose their orientation and direction selectivity. This observation supports our prediction that these neurons receive their orientation and direction tuning from recurrent connections within the local circuit. We further performed a sensitivity analysis with the GLIF network model by sweeping through the major synaptic strength parameters globally (E-to-E, E-to-I, I-to-E, and I-to-I). The results (Fig. S15) demonstrate that we found a suitable, though not unique, parameter regime. Not surprisingly, we observe tradeoffs across the sampled values of parameters, where improvement in one metric comes at the detriment of another.

## Simulating the Models Using Diverse Stimuli

With this final synaptic design in place, we simulate responses to drastically different stimuli – flashes, natural movies, and a looming disk (Fig. 8). This demonstrates the utility of our model as a resource that allows for testing of any kind of visual stimuli.

**Figure 8:**
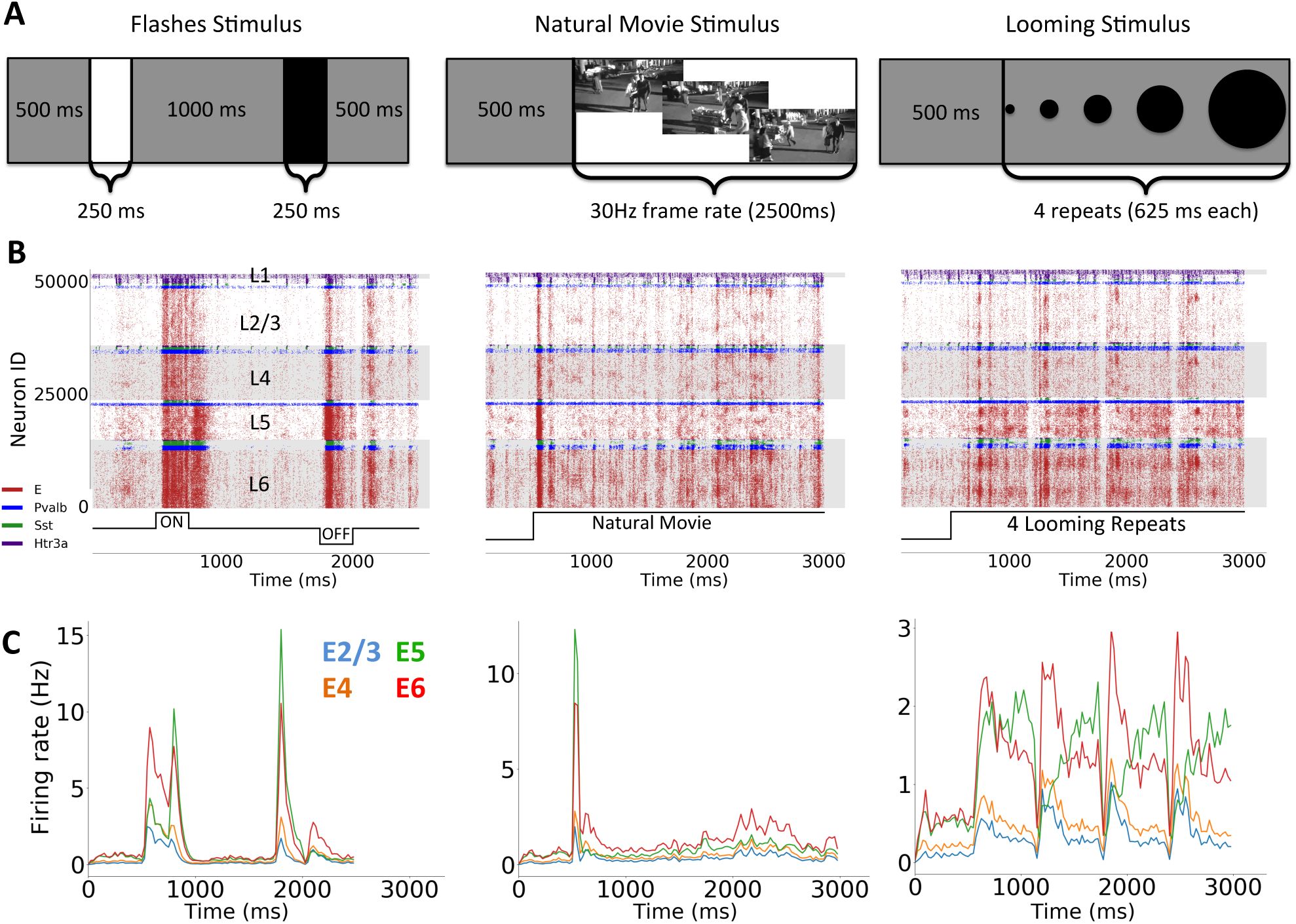
Responses of V1 models to **A:** full-field flashes, a natural movie, and a looming disk, all preceded by 500 ms of grey screen. **B:** Example raster plots for all three stimuli from the biophysical V1 model. **C:** Plots of the mean firing rate across trials for excitatory neurons separated into layers.

A full-field luminance change is one of the strongest stimuli to test the stability of the network (Fig. 8A). Our models remain stable in each of 10 trials with ON (gray screen to white, then back to gray) and OFF (gray screen to black, then back to gray) flashes (Fig. 8B). The networks show strong flash-onset and -offset responses that differ between ON and OFF flashes (Fig. 8C). The natural movie (Fig. 8A) induces varied spatio-temporal patterns (Fig. 8B) and a strong stimulus-onset response (Fig. 8C). Both the flash and natural movie stimuli are available in the publicly available Allen Brain Observatory dataset (de Vries *et al*., 2019) and a thorough comparison between the models and the experiments can be found in the SI (Fig. S16). There we compare temporal response metrics such as the time-to-peak for flashes as well as lifetime sparsity and population sparsity in response to natural movies.

As a further illustration, we simulated a situation not previously investigated - a looming disk on a gray background, (Fig. 8A; see Methods), a class of stimuli used to probe the visual system of various species (e.g., (Gabbiani *et al*., 2002; Yilmaz and Meister, 2013)). Our network responds vigorously (Fig. 8B), with differences in the time course of population firing rates between layers (Fig. 8C). The layer 5 excitatory neurons’ firing rate grows with the stimulus as though tracking the size of the disk, while excitatory neurons in other layers show a strong initial transient response that decreases as the disk grows. This response was not designed for nor expected and demonstrates the use of our models for predictive studies.

## Discussion

We here present two closely related variants of a simulated mouse cortical area V1. Both have an identical network graph, *i.e.*, connectivity, with ∼230,000 nodes of two different flavors, either biophysically-elaborate or highly simplified ones. The variants were constrained by a plethora of experimental data: the representation of the individual cells and their firing behavior in response to somatic current injections, LGN filters, thalamocortical connectivity, recurrent connectivity, and activity patterns observed *in vivo*. This work continues the trend of developing increasingly more sophisticated models of cortical circuits in general (e.g., Traub *et al*., 2005; Zhu, Shelley and Shapley, 2009; Potjans and Diesmann, 2014; Markram *et al*., 2015; Joglekar *et al*., 2018; Schmidt *et al*., 2018) and visual cortex in particular (Wehmeier *et al*., 1989; Troyer *et al*., 1998; Zemel and Sejnowski, 1998; Krukowski and Miller, 2001; Arkhipov *et al*., 2018; Antolík *et al*., 2019). Our main goal was to integrate existing and, especially, emerging multi-modal experimental datasets describing the structure and *in vivo* activity of cortical circuits into biologically realistic network models.

Our models are represented with a standardized data format SONATA (Dai *et. al* 2019, github.com/AllenInstitute/sonata) via the Brain Modeling ToolKit (BMTK, github.com/AllenInstitute/bmtk; (Gratiy *et al*., 2018)) and the open source software NEURON (Hines and Carnevale, 1997) and NEST (Gewaltig and Diesmann, 2007). The SONATA format is also supported by other modeling tools, including RTNeuron (Hernando *et al*., 2013), NeuroML (Gleeson, Steuber and Silver, 2007), PyNN (Davison *et al*., 2009), and NetPyNE (Dura-Bernal *et al*., 2019). Simulations were carried out on 384 CPU cores (12 nodes, 32 CPU cores per node) for the biophysical network and 1 CPU core for the GLIF network. The biophysical network requires ∼90 minutes for 1 second of simulation, whereas the GLIF network requires ∼4 minutes for 1 second of simulation. Thus, the biophysical model can be employed where access to substantial computing resources is available, whereas the GLIF model can be simulated with relatively minimal computing resources.

While our models are similar in size and scope to the Blue Brain Project’s network model of rat somatosensory cortex (Markram *et al*., 2015), they also differ in several ways, besides the obvious distinction of visual vs. somatosensory cortex. Because we turned our efforts primarily towards comparing simulations with *in vivo* experimental recordings, we supply biologically realistic visual stimuli to our models. Thus, we employed a large body of functional data to constrain our thalamocortical inputs, so as to allow arbitrary spatio-temporal image sequences to be simulated. In studying the recurrent connectivity, we placed substantial emphasis on the functional logics of the connectivity and synaptic strengths, which takes into account the stimulus tuning and the potential computational roles of neurons in the circuit, besides the purely structural rules. As a result of these choices, we identified a number of experimentally testable predictions about the relations between the connectional properties and functional tuning of neurons.

Recent studies (Rössert *et al*., 2016; Arkhipov *et al*., 2018) demonstrated that the conversion of a biophysical network model to a GLIF counterpart could result in good qualitative and quantitative agreement in spiking output. We here likewise observed an overall agreement between the biophysical and GLIF models of V1. Although both connectivity graphs are identical, the input-output function of every neuron are different; yet, to judge by their firing rate distributions, the two models act similarly at the population level. This reveals, yet again, the perhaps unreasonable effectiveness of point neuron models given their vastly reduced degrees of freedom (Koch 1999). This is true for both passive (Arkhipov *et al*., 2018) as well as active dendritic models (Rössert *et al*., 2016). A potential reason for this effectiveness at system level simulations originates from their effectiveness at single-cell simulation of input/output transformations. In particular, the individual GLIF (Teeter *et al*., 2018) and biophysical models (Gouwens *et al*., 2018) we use here show similar levels of explained variance when mapping a noisy current injection at *the soma* to an output spike train. It is possible that future models that reconstruct and simulate neurons using dendritic current injection data will outperform the network models presented here. Nevertheless, our current results support applicability of the computationally less expensive GLIF network models (here approximately >8,000 times faster), although ultimately the level of resolution to use should be based on the scientific question under investigation. For instance, computing the extracellular field potential requires spatially extended neurons (Rall and Shepherd, 1968; Lindén *et al*., 2011; Einevoll *et al*., 2013; Reimann *et al*., 2013; Hagen *et al*., 2019). On the other hand, for robust *in-silico* perturbation studies, the GLIF network allows for many more rapid iterations and tests. Developing our V1 simulacra at two levels of resolution enables a larger spectrum of possible studies.

In the process of building and testing the models, we made three major predictions about structure-function relationships in V1 circuits. The first addresses observations that non-Pvalb interneurons (Htr3a/VIP and Sst) show direction and orientation tuning (Liu *et al*., 2009; Kerlin *et al*., 2010; Ma *et al*., 2010), but receive connections from other V1 neurons that are distributed uniformly rather than in a like-to-like fashion (Fino and Yuste, 2011), and little to no LGN input (Ji *et al*., 2015). We thus implemented like-to-like rules for synaptic strengths between excitatory and non-Pvalb inhibitory neurons, which resulted in robust tuning of Htr3a and Sst classes in our models. This tuning was lost when the rule was removed for either of the two classes (Fig. S14).

Our second prediction extends from experimental work investigating functional connections between excitatory neurons (Bock *et al*., 2011; Ko *et al*., 2011; Cossell *et al*., 2015; Wertz *et al*., 2015; Lee *et al*., 2016), thus far primarily in L2/3. Our results suggest that synaptic weights follow rules that are different from the rules that allow two neurons to connect in the first place: whereas the latter are organized in a like-to-like orientation-dependent (symmetric) manner (Ko *et al*., 2011), the former follow direction-dependent (asymmetric) rules (Fig. 6A) with phase dependence (Fig. 6B, E). In our models, these weight rules were implemented among excitatory and inhibitory populations within and across layers (Figs. 6A) to enable realistic levels of orientation and direction tuning (Figs. 7C, D, S12). How can this be reconciled with the report (Cossell *et al*., 2015) that similarity of preferred direction is not a good predictor of synaptic strength (in L2/3)? Because our models employ additional phase-dependent rules (Fig. 6B, E), where incoming connection weights are close to zero outside of a stripe perpendicular to the target neuron’s preferred direction, many presynaptic neurons that share the target neuron’s direction preference connect very weakly to it (if they are outside of the stripe). Therefore, direction similarity by itself is not a strong determinant of weights in our models either, whereas combined with the phase-related geometric constraints it does determine the weights. Interestingly, as we were finalizing this report, a new experimental study (Rossi, Harris and Carandini, 2019) appeared, showing (in L2/3) the preferential location of presynaptic neurons to be within a stripe, as in our connectivity implementation (Fig. 6B, E), thus supporting our prediction (although the new data suggest this architecture may be realized in connection probabilities rather than in synaptic weights).

Our third prediction concerns the asymmetry in cortical retinotopic mapping between the horizontal and vertical axes (Schuett, Bonhoeffer and Hübener, 2002; Kalatsky and Stryker, 2003). This results in higher firing rates for vertical-than for horizontal-preferring neurons, which is not observed experimentally (Fig. 6C). We thus infer the existence of one or more compensatory mechanisms, which may occur at many levels, including connection probability, LGN projections, etc. Our models addressed this at the synaptic strength level (Figs. 6E, F).

These three predictions concern important relationships between the circuit structure and *in vivo* function. The first prediction is significant because mechanisms of tuning of Sst and Htr3a/VIP interneurons are likely to be critical in enabling diverse Sst- and Htr3a-mediated functions (see, e.g., (Liu *et al*., 2009; Kerlin *et al*., 2010; Ma *et al*., 2010; Adesnik *et al*., 2012; Pfeffer *et al*., 2013; Fu *et al*., 2014; Tremblay, Lee and Rudy, 2016; Muñoz *et al*., 2017)). The second prediction suggests a set of general mechanisms that apply across cortical layers and neuronal classes to shape the essential computations of orientation and direction selectivity. The third prediction illuminates the potentially widespread wiring and/or homeostatic mechanisms that equalize firing rates between vertical- and horizontal-preferring neurons. All three predictions are amenable to experimental investigation (Bock *et al*., 2011; Hofer *et al*., 2011; Ko *et al*., 2011; Cossell *et al*., 2015; Wertz *et al*., 2015; Lee *et al*., 2016; Znamenskiy *et al*., 2018; Rossi, Harris and Carandini, 2019).

These findings are a starting point for further elaborations of predictions. The GLIF V1 model, in particular, minimizes the entry barrier to biologically realistic modeling for researchers, due to the low computational demands. Our models, together with all meta-data and code, are freely accessible for download via the Allen Institute for Brain Science’s web portal at brain-map.org/explore/models/mv1-all-layers. We hope that the community will exploit these resources to investigate more biologically refined models of cortex, the most complex piece of active matter in the known universe.

## Acknowledgements

We thank Marius Pachitariu for providing spike-sorting code and assistance. We thank the Allen Institute founder, Paul G. Allen, for his vision, encouragement, and support.

## Methods

For convenience to the reader, we include Supplemental Material 1 that shows the README file from our release (as of this submission). This gives an intuition of what it takes to run the models and complements the methods descriptions here.

### Instantiating the network

The V1 neurons were instantiated and distributed through every layer with raw number estimates available in the supplemental document (document V1_structure.xlsx from our web portal available at brain-map.org/explore/models/mv1-all-layers). We considered the estimated cell densities measured in every layer based on nuclear stains (Schüz and Palm, 1989) with the assumption of an 85% and 15% fractions for excitatory and inhibitory neurons, respectively. The fractions used for the interneuron classes were based on expression levels in double in-situ hybridization experiments (Lee *et al*., 2010). The layer thicknesses were taken from the Allen Mouse Brain Atlas (see Cortical Layer Thickness Measurements). Our model incorporated inhibitory neurons in layers L2/3 through to L6 from three broad classes, Paravalbumin- (Pvalb), Somatostatin- (Sst), and Htr3a-prositive; and excitatory neurons in each layer were considered as one class (Figs. 1A, C). Layer 1 (L1) had only a single inhibitory class of Htr3a neurons (Lee *et al*., 2010; Tremblay, Lee and Rudy, 2016). L2/3 excitatory neurons (class E2/3) were reconstructed from the Cux2 Cre-line, which is almost pan-excitatory in this layer. L4 excitatory cells were reconstructed from four populations of cells studied in our slice recording pipeline (Gouwens *et al*., 2019) – the Scnn1a, Nr5a1, and Rorb Cre-lines, as well as reconstructions from non-Cre-animals. L5 excitatory neurons were sourced from two populations – the cells labeled by the Rbp4 Cre-line and unlabeled L5 neurons. Although L4 and L5 excitatory cells were reconstructed from multiple Cre-lines, it is not known whether cells labeled by these different Cre lines differ in connectivity. Furthermore, they do not appear to show substantially distinct patterns of activity *in vivo* under passive conditions (de Vries *et al*., 2019). Therefore, for all simulations and analyses we combined the L4 and L5 excitatory cells into a single class per layer (E4 and E5). L6 contained one excitatory class (E6), with neurons from Ntsr1 Cre-line only (due to availability at the time of creating the models). Altogether, we used 112 unique neuron models for the biophysical and 111 for the GLIF networks. At time of model building, there were no Htr3a reconstructions for L6 neurons and therefore we re-used the two deepest L5 Htr3a models to populate this cell class in L6. Although the Allen Cell Types Database had more cell models, not all models could fit geometrically in the V1 volume without protruding beyond the pia. This was due to Cre-lines not labeling specific layers exclusively, resulting in cases where cells from certain Cre-lines resided in adjacent layers (see **Somatic Coordinates**).

The neuron models were fit to *in-vitro* measurements (Gouwens *et al*., 2018; Teeter *et al*., 2018) and are publicly available via the Allen Cell Types Database (celltypes.brain-map.org/). All our biophysical models used passive dendrites although the Allen Cell Types Database includes neuron models with active dendritic conductances. This was due to active-dendritic models being too computationally expensive (prohibitively) for the extent of our work. Further, the somatic spike output from the active-dendrite models do not show much better performance than the models with active conductances restricted to the soma (celltypes.brain-map.org/). Therefore, we used the less computationally expensive neuron models.

#### Cortical Layer Thickness Measurements

Layer thicknesses for the model were taken from the Allen Mouse Brain Atlas (Oh *et al*., 2014 - atlas.brain-map.org/). They were calculated from a mouse common coordinate framework in which voxels were annotated with cortical areas and layers. In this framework, streamlines were calculated that connected pia to white matter using the shortest paths (Oh *et al*., 2014 - Documentation in atlas.brain-map.org/). For each voxel on the surface of V1, the thickness of each layer was calculated along the associated streamline, and the median values across all of V1 were used to construct the model.

#### Somatic Coordinates

With the number of neurons identified (V1_structure.xlsx), we needed to assign somatic coordinates for every cell and select appropriate neuron models. For the biophysically detailed neurons we also had to assign to a neuron a rotation about the depth axis (white-matter to pia). This is due to our V1 model using a fixed number of reconstructed neuron models relative to the total number of neurons simulated and hence when reusing a model, we randomly rotated the individual neurons between 0 and 2π around the depth axis. For the somatic coordinates, cells for each population were uniformly distributed within a cylindrical domain and within the specified layer depth. For the biophysical models, the depth of a neuron would affect which neuron model was assigned to it. The first condition was that a model would not be assigned to a particular cell if that model’s morphology significantly extended out of the pia when placed at the cell’s somatic location (with a tolerance of 100 μm). Once all putative cell models that pass this criterion were identified, we randomly selected a model based on a Gaussian probability density function (with standard deviation of 20 μm).

#### Visual Coordinates

Neurons’ positions are defined in the physical space, whereas visual stimuli (see **Visual Stimuli**) supplied to the models, as well as the LGN filters converting these stimuli to spike trains impinging on V1 neurons, are defined in the visual space. Thus, a mapping between the two spaces needs to be defined. The cortical plane (plane perpendicular to the depth axis) was mapped to the visual space, with the geometrical center of the model corresponding to the center of the visual space. Retinotopic mapping experiments in the mouse V1 identified how much displacement in visual cortex corresponded to displacements in visual space (Schuett, Bonhoeffer and Hübener, 2002; Kalatsky and Stryker, 2003). Using these results (Figure 3 from (Schuett, Bonhoeffer and Hübener, 2002) and Figure 4 from (Kalatsky and Stryker, 2003)), we approximated that the visual degrees traversed per mm of cortex are 70 degrees/mm in the azimuth and 40 degrees/mm in elevation. Note the asymmetry between the two directions. From this we can convert any translation of azimuth and elevation in cortex to a translation in visual space. For example, consider moving 845 μm in the azimuth (radius of the V1 model): the movement in visual space is then estimated to be 0.845 mm * 70 degrees/mm = 59.15 degrees. The somatic position of every neuron was used, via such translations, to establish the assigned neuron’s position in the visual space, which was then used in algorithms establishing connectivity from the LGN to V1 (see below).

### Thalamocortical Connectivity

#### Distributing LGN Units

We sought to create an LGN model that roughly captures the entire LGN with an estimated 18,000 neurons in the mouse. In our model, we do not explicitly model the shell and core of the LGN and simply distribute the LGN units on a 2D plane in visual space to model 240 degrees (horizontal) by 120 degrees (vertical). We imposed a lattice structure on the 2D plane by dividing it into girds (15 blocks horizontally by 10 blocks vertically of size 16×12 degrees). Each block had a total of 116 LGN units (Table 1) distributed uniformly within the block to give a total of 17,400 LGN units that can process arbitrary visual stimuli.

**Table 1:** Distribution of LGN unit numbers in every block and the receptive field sizes per class.

| LGN Class | Units per block | Spatial size range (degrees) |
| --- | --- | --- |
| *sON-TF1* | 7 | [2, 9] |
| *sON-TF2* | 5 | [2, 9] |
| *sON-TF4* | 7 | [2, 9] |
| *sON-TF8* | 15 | [2, 9] |
| *sOFF-TF1* | 8 | [2, 9] |
| *sOFF-TF2* | 8 | [2, 9] |
| *sOFF-TF4* | 15 | [2, 9] |
| *sOFF-TF8* | 8 | [2, 9] |
| *sOFF-TF15* | 7 | [2, 9] |
| *tOFF-TF4* | 10 | [2, 9] |
| *tOFF-TF8* | 5 | [2, 9] |
| *tOFF-TF15* | 8 | [2, 9] |
| *sONsOFF* | 8 | 6 |
| *sONtOFF* | 5 | 9 |

Each LGN unit is represented by a spatio-temporally separable filter, which operated on the movies in the visual space as inputs, and returned a time series of the instantaneous firing rate as output (this rate was then converted to spikes in each individual trial using a Poisson process). The spatial components of the LGN filters are spatially symmetric two-dimensional Gaussian kernels and the temporal components are a sum of weighted raised-cosine bump basis functions (Pillow *et al*., 2005). The temporal kernel was designed to have a bi-phasic impulse response:

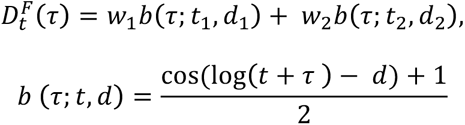

where there are six parameters: i) two time constants (*t*_1_, *t*_2_) for the basis functions, ii) two weights (*w*_1_, *w*_2_) used to linearly sum the functions and iii) offsets (*d*_1_, *d*_2_). All data and code are available through the BMTK (github.com/AllenInstitute/bmtk). The spatial and temporal filters are combined to form a 3D spatiotemporal kernel to respond to input signals that are grayscale, represented on a −1 to 1 scale (from black to white), with a time step of 1 ms.

The LGN filters were sampled from 14 classes (Table 1) that approximated the diversity observed in experimental recordings *in vivo* (Durand *et al*., 2016) (see Main Text and **Fig. 2A**). The LGN filter parameters used for every class were obtained by fitting filter responses to the mean experimental responses for every class (resulting parameter values are available in the BMTK). A ±2.5% jitter was added for every parameter when instantiating individual LGN filters. We observed that receptive field sizes of cells from most of the LGN classes in the experimental recordings (Durand *et al*., 2016) spanned a large range within class. We thus assigned every LGN unit a randomly generated spatial size within the recorded ranges drawn from a triangular distribution defined as follows: zero at lower bound, peak at the lower bound plus 1 degree, and then zero again at the upper bound (to approximate the experimental distributions).

#### Thalamocortical Architecture Impact on Direction Selectivity

The major guiding purpose for creating thalamocortical connections in our V1 models was to enable direction selectivity, which was proposed to arise due to integration of sustained and transient LGN inputs by V1 cells (Lien and Scanziani, 2018). Before instantiating such rules for the full-scale model, we performed a simplified theoretical analysis to investigate how combinations of transient and sustained pools of LGN inputs, using biologically realistic parameters, would create direction-selective responses in target V1 cells. For this analysis we approximated the LGN input to a V1 cell using a sustained ON and a transient OFF subfields.

For the thalamocortical projections to a V1 neuron in our full models (see **Forming Thalamocortical Connections**), we would first identify all suitable LGN filters that have overlapping retinotopic positions with the V1 cell. This pool of filters was then split into a sustained subfield ellipse in one half of the receptive field and a transient subfield ellipse in the other half (Fig. 2C). The orientation of the ellipses would depend on the assigned preferred angle of the V1 neuron. The ellipses’ major axis would be perpendicular to the preferred orientation of the V1 neuron and the sustained subfield would be positioned such that it is activated first in the case of a bar moving in the preferred direction of the V1 neuron (Fig. 2D). We would then randomly select filters from within these ellipses from the population of sustained or transient LGN filters (Figs. 2C, 3A). For the simplified theoretical analysis here, we consider the sustained ON and transient OFF subfields, represented by a single elliptical filter each, approximating contributions from all LGN cells within a subfield.

The synaptic input current from one of the subfields (labeled as *F* = *ON* or *F* = *OFF*) to the V1 cell in response to a stimulus is then described by

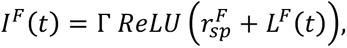

where Γ is the constant determining the magnitude of the current (assumed to be the same for both subfields), 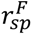 is a baseline (spontaneous) firing rate, and *ReLU*(*x*) is a rectified linear unit function that is zero below a threshold (here set at zero) and linear above the threshold. The response is dependent on the stimulus *S*(*x*, *y*, *t*):

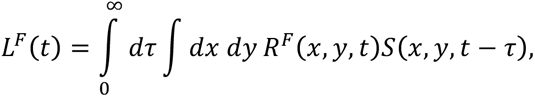

We consider the case where the two subfields are offset along the x-axis, so that each subfield is described as:

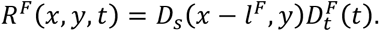

The assumption used here is that each kernel is spatio-temporally separable.

The temporal kernel used here is a sum of weighted raised-cosine bump basis functions as used above (Pillow *et al*., 2005; see **Distributing LGN units**). The spatial kernel is described by an elliptical Gaussian profile:

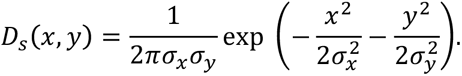

with the standard deviations *σ_x_*, *σ_y_*, respectively. We will study a special case of subfields separated by a distance *d* along the x-axis using *l^ON^* = *d*/*2* and *l^OFF^* = −*d*/*2*:

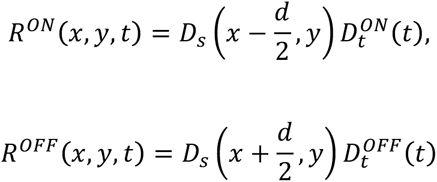

Let us examine the response of a cell to moving grating stimuli having maximum luminance *S_max_* and a contrast *c*:

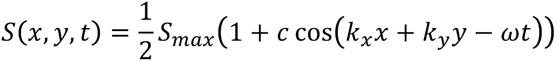

where ***k*** = (*k_x_*, *k_y_*) defines the direction of the grating wave front: *k_x_* = *k_cos_*(*θ*), *k* = 2*πSF*, *ω* = 2*πTF* and *SF* (cpd) and *TF* (Hz) are the spatial and temporal frequencies of a grating, respectively.

It is more convenient to work in the complex space:

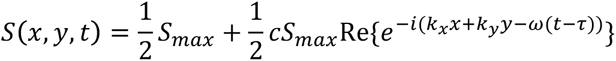

The input current from each subfield is 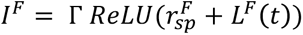 where 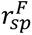 is independent of stimulus and 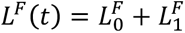 is a stimulus dependent response:

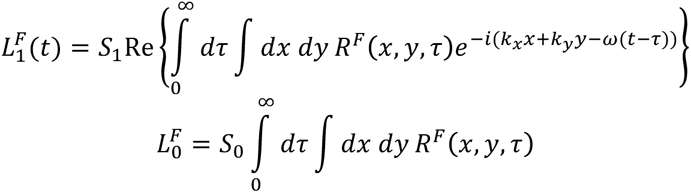

Here we use a short hand notation 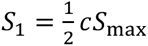 and 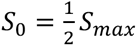.

Substituting *R^F^* (*x*, *y*, *t*) we find:

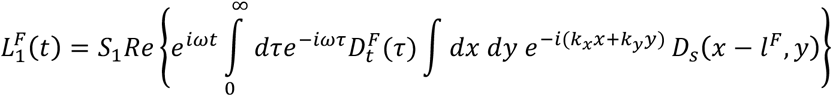

Since the temporal kernel 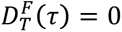 when *τ* < 0, we can simply extend the integration to negative infinity over *τ*.

The temporal integral in 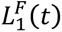 is the Fourier transforms over time:

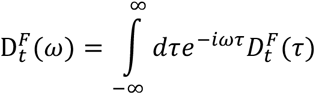

that could be expressed using the magnitude 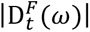 and phase *ψ^F^*(*ω*):

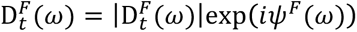

The spatial integral 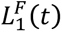 in is the spatial Fourier transform:

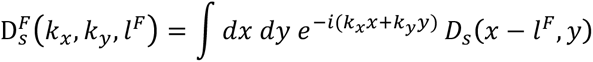

Thus, we can express 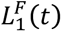 as

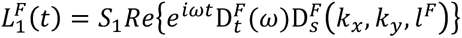

Thus, the response to a drifting grating with temporal angular frequency *ω* is determined by the Fourier component at that frequency only. We can compute the temporal components (raised cosine bumps) Fourier transforms numerically.

We can compute the spatial transform analytically to find:

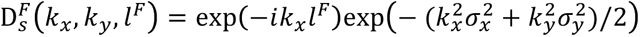

which has an amplitude:

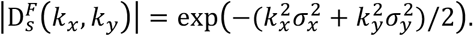

so that:

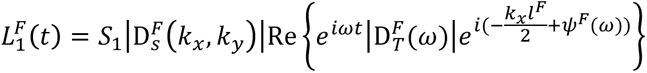

The total input current to a cell is the sum from the two subfields:

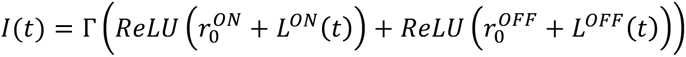

Using these equations, we can estimate both the direction selectivity index (DSI) and the orientation selectivity index (OSI) of the F0 and F1 components for a variety of filter parameters: subfield separation *d*, ellipse aspect ratio or width (determined by *σ_x_*, *σ_y_*), and temporal parameters. The F0 response is a commonly used metric that calculates the cycle average mean of the response to a drifting grating while the F1 component computes the modulation response at the input temporal frequency (Movshon, Thompson and Tolhurst, 1978).

We used filter parameters from sON-TF8 and tOFF-TF8 as well all other default values: *d* = 5 degrees (Lien and Scanziani, 2013), SF = 0.025cpd, TF = 8Hz, ellipse aspect ratio = 3.0, ellipse minor axis = 4.0 degrees. For a set of fixed stimulus (drifting grating), we changed one parameter at a time and observed the impact on OSI and DSI. For the distance between the elliptical sustained and transient subfields (*d*; Fig. S1A), we note that the F1 component switches direction preference (i.e., its DSI changes sign) as *d* grows, due to a shifting phase difference between the subfields. The DSI of the F0 component is always zero as the net input remains constant for the preferred and null directions, consistent with experimental recordings (Lien and Scanziani, 2013, 2018). On the other hand, the OSI of the F0 component is constant but non-zero due to the elliptical structure of the subfields that biases the net input per grating cycle for specific orientations (but not directions). The OSI of the F1 component is positive even when *d* = 0 due to the elliptical shape of the subfields (and temporal properties). Second, by varying the sustained time-to-peak parameter (starting from the transient subfield’s time-to-peak of 30ms, Fig. S1A), we observe, as expected, that asymmetry in the temporal properties of the subfields is essential for producing direction selectivity. There is no direction selectivity in the F1 component when both filters are identical temporally; but as the time-to-peak of the sustained subfield increases, there is a quick rise in F1 DSI. This is followed by a reversal in the direction preference for very high (non-biological) time-to-peak values. The F1 OSI shows a sharp monotonic decrease with the sustained time-to-peak while the F0 OSI is non-monotonic but roughly constant. Other changes investigated in the subfield parameters were the aspect ratio of the ellipses and the size of the ellipses that both showed relatively constant F1 DSI as both ellipse sizes were altered together (Fig. S1A). On the other hand, the OSI values showed a monotonic increase with both illustrating the contribution of the elongated structure for endowing orientation selectivity. An aspect ratio of one still showed some orientation selectivity due to the temporal offsets of the filters giving slight orientation selectivity (our OSI metric is based on circular variance, see Orientation Selective Index below).

We next investigate the effect of changing the spatial frequency of the drifting grating (Fig. S1B). As before, the F0 DSI always remains zero. As the spatial frequency increases, we again observe a reversal in the preferred direction for the F1 component as observed experimentally in mouse cortex (Billeh *et. al* 2019). For orientation selectivity, the F1 OSI shows a sigmoidal increase as spatial frequency increased while the F0 OSI shows a peak with a fast decay due to reduced responsiveness of the LGN ellipses to high spatial frequencies. On the other hand, the F1 OSI is relatively flat while the F0 OSI shows a peak response as a function of temporal frequency, albeit with a slower decay, again due to the reduced responsiveness of the LGN subfields to high temporal frequencies (Fig. S1B). For our choice of subfield parameters, the F1 DSI does not switch sign as we varied temporal frequency, but such switching can occur as observed experimentally and with different filter properties and time constants (Billeh *et. al*, 2019).

In summary, these simplified calculations confirm that the overarching model of the integration of sustained and transient LGN responses (Lien and Scanziani, 2018) indeed enables directionally selective input currents into V1 cells when biologically realistic parameters are used. Given this reassuring result, the next step was to create a similar architecture of connections to the V1 model from the thousands of filters representing LGN cells in the visual space.

### Forming Thalamocortical Connections

The connections from the LGN to V1 neurons followed an approach similar to previous work (Arkhipov *et al*., 2018). The first step was to establish shared retinotopy between the V1 neurons and the LGN units. The coordinates of the LGN units were in visual space (degrees) while the V1 neurons’ coordinates were in regular 3D space mapped to the cortical surface and white-matter-to-pia depth (see **Somatic Coordinates**). By imposing that the center of the V1 model mapped to the center of the visual space, the location of each V1 neuron was converted to visual space using the cortical magnification factor, as described in section **Visual Coordinates**. This procedure assigned each V1 neuron a position in visual space, which may be expected to correspond approximately to the center of that neuron’s RF in the complete model. We then identified which LGN units would project to every V1 neuron (from the classes to receive LGN inputs; see Main Text and **Table 2**), as follows.

**Table 2:** Properties of the subfields in the visual space used to select LGN neurons projecting to V1 neurons (for every cell class receiving LGN inputs; the remaining classes are assumed to receive no LGN input (Ji *et al*., 2015)). The connection probability refers to probability a neuron receives input from the LGN (Ji *et al*., 2015). The mean LGN input current corresponds to the mean excitatory LGN current the neuron class receives (Lien and Scanziani, 2013; Ji *et al*., 2015) when voltage clamped at the inhibitory synapse reversal potential (see **Thalamocortical Synaptic Weights**). The V1 TF column represents the preferred temporal frequency of the V1 neuron class (Niell and Stryker, 2008; Durand *et al*., 2016). The S_ON_ Ratio refers to the probability the sustained component will be ON instead of OFF (Lien and Scanziani, 2013) – the transient component was always OFF. The Separation Range refers to the distance between the sustained and transient subfield ellipses – E4 estimated from Lien and Scanziani, 2013. The Width Range refers to the minor-axis width of the ellipses (diameter). The Aspect Ratio refers to the length of the major-axis relative to the minor-axis. Note the aspect ratio is relative to neurons’ visual space center and once sizes of LGN receptive fields are incorporated, the results match experimental measures (Lien and Scanziani, 2013) more accurately as shown previously (Arkhipov *et al*., 2018). The final column refers to the number of synapses an LGN neuron makes to a V1 neuron if a connection exists. This was extrapolated from experimental work (Morgenstern, Bourg and Petreanu, 2016) as discussed in **Thalamocortical Synapse Estimate**.

| V1 Neuron Class | Connection Probability | Mean LGN current (pA) | V1 TF (Hz) | S<sub>ON</sub> Ratio | Separation Range (deg) | Width Range (deg) | Aspect Ratio Range | Number of Synapses |
| --- | --- | --- | --- | --- | --- | --- | --- | --- |
| *i1Htr* | 0.588 | 29.0 | 2.0 | 0.75 | [6, 10] | [8.5, 11] | [2.2, 2.4] | 10 |
| *E2/3* | 0.789 | 20.3 | 1.5 | 0.90 | [4, 6] | [7.5, 9.5] | [3.4, 3.6] | 15 |
| *i2/3Pvalb* | 0.824 | 50.8 | 2.0 | 0.75 | [6, 10] | [10, 13] | [1.6, 1.8] | 15 |
| *E4* | 1.000 | 46.0 | 2.0 | 0.90 | [4, 6] | [7.5, 9.5] | [3.4, 3.6] | 80 |
| *i4Pvalb* | 1.000 | 119.8 | 2.0 | 0.75 | [6, 10] | [10, 13] | [1.6, 1.8] | 75 |
| *E5* | 1.000 | 20.3 | 1.5 | 0.50 | [8, 12] | [12, 16] | [1.6, 1.8] | 15 |
| *i5Pvalb* | 1.000 | 63.7 | 2.0 | 0.50 | [6, 10] | [10, 13] | [1.6, 1.8] | 20 |
| *E6* | 0.778 | 17.1 | 1.5 | 0.90 | [3, 4] | [9, 11] | [3.4, 3.6] | 15 |
| *i6Pvalb* | 0.818 | 44.1 | 2.0 | 0.75 | [6, 10] | [10, 13] | [1.6, 1.8] | 10 |

Given the directionally selective architecture to be imposed, every V1 neuron was assigned a preferred angle of stimulus motion to determine the placement of the elliptical subfields from which LGN units would be sampled (Figs. 2C, 2D, 3A). There was always a transient OFF subfield and a sustained subfield that was either ON or OFF (this choice was made based on the relative abundance of the different classes of LGN cells in our experimental recordings (Durand *et al*., 2016), as summarized in Fig. 2A). The two subfields were identically oriented and offset by certain distance; the offset and the short axes of both ellipses were co-aligned with the assigned preferred direction of the target V1 neuron. The position of the target neuron was at the middle of the line connecting the centers of the two subfields (Fig. 3A). The subfields were positioned along the vector of the preferred direction of the target neuron in such a way that the vector pointed from the sustained subfield to the transient one (Figs. 2C, 2D). Note that the assigned angle was also used for the recurrent connectivity (see below) and was set such that every V1 neuron class represented every angle in the range [0, 360°) with even spacing. The dimensions of the subfields and their separation varied based on the V1 neuron’s class (**Table 2**); these choices were made according to estimates of the expected metrics – such as the OSIs and DSIs – for the class, based on experimental reports (see details and references in V1_parameter_estimate.pptx). The subfield parameters for the E4 target population were informed by our previous model of L4 (Arkhipov et. al, 2018), and parameters for the other populations were chosen following the assumption that V1 cell classes with stronger orientation/direction selectivity would utilize smaller and more elongated LGN subfields. Importantly, we chose these subfield parameters once and did not vary them to tune the model for target OSI/DSI values. The good agreement with the experiment observed for the final model (**Fig. 7**) suggests that our initial choice of these subfield parameters was appropriate (and, to the best of our knowledge, it is consistent with available experimental observations); however, it is possible that the agreement could be further improved by tuning the subfield parameters.

As reported previously, a linear angle approximation was used (Arkhipov et. al, 2018). Further, every V1 neuron was assigned a preferred temporal frequency drawn from a Poisson distribution with a mean as measured experimentally (**Table 2**, (Niell and Stryker, 2008; Durand *et al*., 2016)). This determined the probability of selecting LGN units preferring particular temporal frequencies. Given that there was a discrete number of LGN filters for every class (sON, sOFF, tOFF), the probability of selecting a particular subclass (i.e. a particular TF) was based on the distance of the V1 neuron’s temporal frequency from the LGN unit’s preferred temporal frequency, divided by the total possible distance for that class.

Once the subfields were established, the LGN units to be connected to the target cell were selected among the units that had the centers of their spatial kernels within the subfields (and of the LGN class matching to each subfield, see **Fig. 3A**). From this total pool, LGN units were connected randomly based on the probability of connections (given their temporal frequency as mentioned above). Thus, not every LGN unit in the subfield formed a connection with the target V1 cell (Figs. 2C, 3A). Finally, for the ON/OFF filters, a restriction was set that required the axis of the ON/OFF subfield to be within 15-degrees relative to the assigned orientation preference angle of the V1 neuron (Arkhipov et. al, 2018). With all these choices, the suitable LGN units were selected probabilistically to project to each target V1 cell. Based on these rules, the average number of LGN units connecting to a V1 cell for excitatory neurons is: 19.3 ± 6.0 (mean ± SD), median = 19, min = 2, max = 47. For inhibitory neurons: 15.0 ± 4.4 (mean ± SD), median = 15, min = 2, max = 32. The mean number of LGN projecting units to V1 neurons is below the recently reported estimates (Lien and Scanziani, 2018); although the authors themselves acknowledge their measurements are likely overestimates. Nevertheless, the most important parameter is the total synaptic current that every population receives (see **Thalamocortical Synaptic Weights**) which was matched to experimental measurements (Lien and Scanziani, 2013, 2018) and could compensate for the differences we have in this version of the model.

#### Thalamocortical Synapse Estimate

For the biophysical model we estimated the number of synapses impinging on different V1 neurons. The exact numbers of synapses are only estimates as the more critical step was ensuring the total excitatory current received from the LGN matched experimental measurements (see below). Should the number of synapses be incorrectly estimated, this was compensated for by the final synaptic weights.

Our calculation and formalism for the number of thalamocortical synapses per neuron is described below; we also provide a supplementary document (Num_TC_synapses.xlsx) where all the calculations were done. As the field advances, in particular with electron-microscopy technology, we would need fewer assumptions and simply use the available data. In the model, synapses were placed along the dendrites up to 150 μm away from the soma but excluding the soma, as done in a previous model of the layer 4 of V1 based on experimental reports (Schoonover *et al*., 2014; Arkhipov *et al*., 2018)

One key resource we used was the fluorescence measurements of the density of thalamocortical axons across cortical depth (Morgenstern, Bourg and Petreanu, 2016). We used this work to determine the fraction of fluorescence across cortical layers as an estimate of the fraction of LGN projections to different layers. The full calculation is in the accompanying supplemental document (Num_TC_synapses.xlsx) and here we explain our technique and assumptions. In particular we assume the Fluorescence Signal (FS) is a function of the following factors:

1. Number of cells in a layer (Schüz and Palm, 1989)
2. Percentage of cells that actually get innervated in a layer from the LGN (Ji *et al*., 2015)
3. At a specific depth (layer), the proportion of dendrites from cells in different layers that extend to other layers

a. For inhibitory neurons, dendrites where assumed to stay within their layers and not extend to other layers.
4. The fraction of LGN synapses on a stretch of dendrite is the same whether that dendrite is from an E or Pvalb cell.

a. Assumption includes that, out of all interneurons, Pvalb cells are the only ones to receive significant innervation except for layer 1 (Ji *et al*., 2015).

From here, for a specific layer, the below calculation was used to approximate the fluorescence signal (FS) from labeled thalamocortical axons. This example is for layer 4:

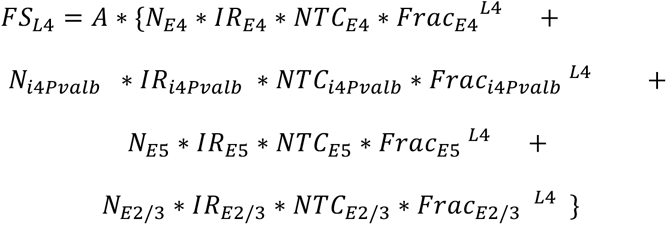

In the accompanying document, this is found by summing the rows for the gray matrix. The different notations mean:

- *FS_L_*_4_ = Fluorescence signal in layer 4
- *A* = Constant factor converting fluorescence signal to biological innervation numbers. We assume fluorescence is a linear function of axon density, and so *A* is constant for every layer. We will need to solve for *A* (see below)
- *N_E_*_4_ = Number of excitatory cells in L4 (Schüz and Palm, 1989)
- *IR*_*E*4_ = Innervation ratio of LGN onto L4 pyramids (Ji *et. al, 2015*)
- *NTC*_*E*4_ = Number of synapses that are thalamcortical for every L4 excitatory cell – the numbers we are seeking for every layer. From (4) above, it is assumed that *NTC*_*i*4*pvalb*_ = *NTC*_*E*4_.
- *Frac*_*E*4_^*L*4^ = The fraction of excitatory cells’ dendrites in L4 that is contributed from L4 cells (from assumption (3) above). See the light green matrix in the accompanying excel sheet.
  - Note that *Frac*_*E*4_^*L*4^ + *Frac*_*E*2/3_^*L*4^ + *Frac*_*E*5_^*L*4^ = 1.
  - Note that we assumed *Frac*_*E*6_^*L*4^ = 0 and thus that is not included in the above example of L4.
  - Note that *Frac*_*i*4*Pvalb*_^*L*4^ = 1 is assumed for all layers for Pvalbs (assumption (3.a) above).

We note that the document had a finer division of every layer (split in two: upper (A) and lower (B) components) and the idea of single layers here is just used for explanatory purposes.

All these assumptions can be written in a matrix form as follows:

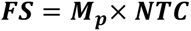

Where ***FS*** is an Nx1 matrix of the fluorescence signal across layers and ***NTC*** is the Number of thalamocortical synapses that is also Nx1. ***M_p_*** holds the properties described above and is a matrix of dimensions NxN (contributions from all layers). We can thus solve for ***NTC*** by taking the inverse:

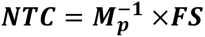

Since the constant factor *A* is not known, the values of ***NTC*** are not the actual numbers of synapses. To account for this, we use the experimental finding that, in the mouse visual cortex, the number of thalamocortical synapses on L4 excitatory cells is approximately 1200-1500 (Schoonover *et al*., 2014; Arkhipov *et al*., 2018). This gives us the scaling factor to account for *A* and hence allows us to estimate NTC for all layers.

For the supplemental document, which was divided into finer divisions, 1200 was used as the average of all the L4 divisions (see scaling factor). The final numbers of synapses are shown in **Table 2**.

#### Thalamocortical Synaptic Weights

Various studies have identified the thalamic innervation pattern into the visual cortex across laminae (Lien and Scanziani, 2013, 2018; Kloc and Maffei, 2014; Schoonover *et al*., 2014; Ji *et al*., 2015; Morgenstern, Bourg and Petreanu, 2016; Bopp *et al*., 2017). We used these results to identify the total current that different cell classes should receive from the LGN. One study, already published during building of the model, measured that the net current into layer 4 excitatory cells responding to drifting gratings at their preferred angle was on average 46 pA (Lien and Scanziani, 2013). Other work using optogenetic stimulation identified the cell classes that are innervated by the thalamus, for both the probabilities and relative strengths (Ji *et al*., 2015). Assuming linear scaling to layer 4 excitatory neurons, we estimated the target mean current for every cell class in response to a grating at a neuron’s preferred direction (**Table 2**).

To attain the target currents, for the biophysically detailed model, we created networks that had 100 cells from every model, all preferring a single direction, that receive LGN innervation as described above (but no other connections). A grating at 2Hz, full contrast, full field with a spatial frequency of 0.04 cycles per degree (to match the experimental work precisely (Lien and Scanziani, 2013)) was shown to these networks. Further, the neurons were clamped at the reversal potential of the inhibitory (GABA) synapses in our model (again as performed experimentally). The net mean current during exposure was measured and the synaptic weights iteratively adjusted until the target current was reached with 2% tolerance. For surrounding LIF neurons, for the same stimulus, we matched the firing rates that were observed with purely LGN input in the biophysically detailed core neurons of the same class. As mentioned in the Main Text, during optimization of the full V1 model the weights of synapses from LGN to excitatory layer 4 cells were not adjusted at all, given that the measurements we used as targets in the procedure described here were of high precision and obtained *in vivo* (which is the condition we were aiming to match in our full model). Weights of all other synapses from LGN were adjusted, but the adjustment was allowed to be no more than by a factor of 2 for the mean input current (Table 2).

Finally, the GLIF V1 model used the same strategy to attain the same target mean currents using the same grating LGN stimulus. However, as the GLIF models employed in the V1 model were using post-synaptic current based synapses (see Synaptic Characteristics), the weights were initially set as the target currents and no voltage clamping was required. However, the average rheobase (minimal current step amplitude to elicit an action potential) of the GLIF models in the model are bigger than experimental measurements (Fig. S17), except for Pvalb neurons that had smaller rheobase values. To match closely to the experimental data, the established weights from LGN to V1 were scaled by the average ratio between average rheobase of GLIF model and experiment data i.e., 0.81 for Pvalb population and 1.36 for other populations.

### Background Connectivity

A second source of input to the V1 models was a background to coarsely represent the “rest of the brain”. This was modeled as a single input unit that fired at 1 kHz with a Poisson distribution. All neurons received connections from this unit, and the weights were optimized (at the same time with the optimization of weights for the recurrent connectivity) to ensure the V1 spontaneous firing rates matched target experimental rates (see below).

### Recurrent Connectivity

The cortico-cortical connection probabilities for different cell-class pairs were estimated based on an extensive and systematic survey of the existing literature and curated into a resource that we make publicly available (Fig. 4, see details and notes regarding assumptions and the literature used in Connection_probabilities.pptx). It is important to note that in many cases the values reported in the literature do not take into account two effects that strongly influence connection probabilities. The first is distance dependence: cells closer to each other typically have a higher chance of being connected than cells further apart. The second is that connection probabilities can be affected strongly by differences or similarities in functional preferences of cells, such as preference for orientation. Pyramidal cells in L2/3 of mouse V1, for instance, have a higher chance of being connected with one another if they prefer similar orientations, compared to orthogonally tuned cells (Ko *et al*., 2011; Cossell *et al*., 2015; Wertz *et al*., 2015; Lee *et al*., 2016). Based on these two factors, the adjustments described below were made.

It is reasonable to assume, for the mouse visual cortex, that both these factors are independent (given the “salt and pepper” arrangement of orientation tuned cells in the mouse (Harris and Mrsic-Flogel, 2013; Seabrook *et al*., 2017)) and thus the total probability of connection for a cell-class pair is a product of the distance-dependent and preferred-angle-dependent factors (functions of *r* and *Δϕ*, respectively):

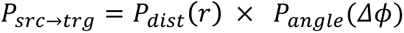

First we will discuss each of the components separately, and the final section will illustrate our approach for combining the two.

#### Distance dependent adjustment

We noted that the majority of the experimental literature reporting probability of connections tended to consider inter-somatic distances that were within approximately 0 − 50 μm to 0 − 100 μm. Since we aimed to have a Gaussian profile for distance dependence (Levy and Reyes, 2012), the probability at the origin had to be adjusted to account for these measurements. Since measurements were made in the approximate range of 50 – 100 μm for the upper bound, we chose to consider the mid-point of 75 µm as our reference point for such upper bound. Note the distance is only measured in a plane and is independent of cortical depth in our calculations.

For the Gaussian probability distribution:

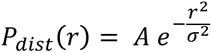

Given our assumptions, the integral of this probability from 0 to *R*_0_ = 75 μm, divided by the area within the radius *R*_0_, should be equal to the reported measured probability, *P_rep_*:

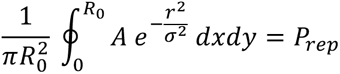

Converting to polar coordinates:

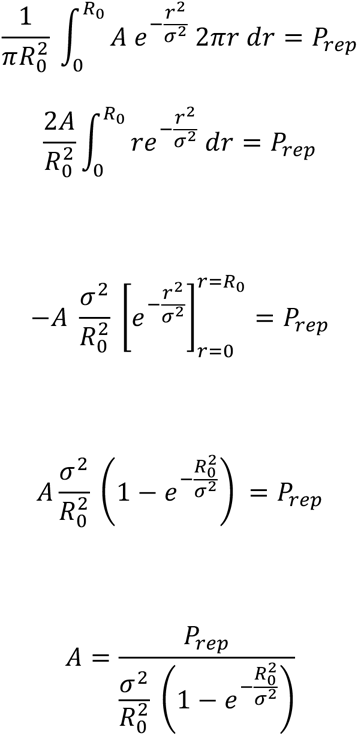

This establishes the relationship between the values reported in the literature and our distance-dependent formula for connection probability.

From work in the mouse cortex (Levy and Reyes, 2012), the standard deviations were estimated to be (Fig. 4):

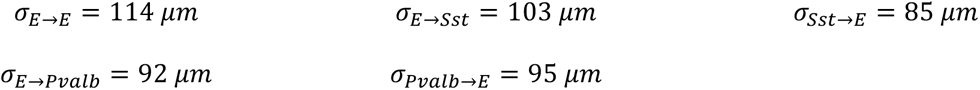

From internal data at the Allen Institute during model building:

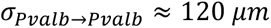

In the absence of data for other connection classes, we assumed that connections between excitatory neurons and Htr3a neurons follow the same dependence as between excitatory and Sst neurons (bidirectionally). Finally, we also assumed that connections among all inhibitory classes have the same distance dependence (i.e., same as *σ_pvalb→pvalb_*).

#### Orientation tuning adjustment for excitatory-to-excitatory connections

For orientating tuning dependence, our system is modeled such that pairs containing cells with similar preferred orientation angles have higher connection probabilities than pairs of orthogonally tuned cells, when the presynaptic neuron is excitatory (like-to-like connectivity) (Ko *et al*., 2011; Cossell *et al*., 2015; Wertz *et al*., 2015; Lee *et al*., 2016). Here we assume the dependence is linear (Fig. 4D) as a function of the orientation tuning difference (*Δϕ*):

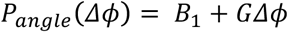

Since we considered orientation selective tuning for connectivity (not direction selective), the difference of preferred angles of any two cells can be compressed to be between 0° and 90°. For this model, we can see that the intercept occurs at (0°, *B*_1_). At the other extreme of the model, we set the point to be (90°, *B*_2_). The relative strength of the dependence can be described by a ratio *Q* = *B*_2_/*B*_1_. As can be seen, for like-to-like, *Q* < 1 (i.e., *G* < 0).

Our model is developed such that the integral of the function *P_angle_*(*Δϕ*), normalized by the range of *Δϕ*, is always equal to 1. This was implemented because this function is used as a multiplier with the distance dependence function *P_dist_*(*r*), and since we assume that experimentalists measuring *in-vitro* probability of connections sample equally from cells preferring all possible orientation angles *in vivo*. This does restrict the ratio *Q* one can select, based on the distance dependence and measured connection probabilities from experimental literature. As will be discussed below, if the ratio is outside of a suitable range, we rescaled it to reach the correct range.

Because *B*_1_= *QB*_1_,

the gradient can be expressed as:

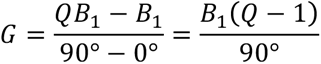

Integral of *P_src→trg_*(*Δϕ*) (with normalization for the angle range) should be set to 1 to determine the scaling factor:

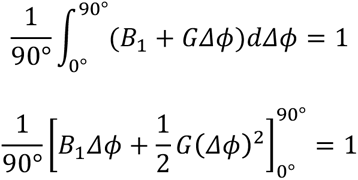

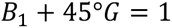

Substitute *G*:

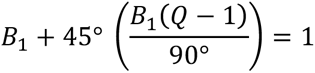

Solving for *B*_1_:

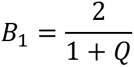

And thus:

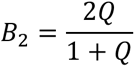

The value of *Q* for layers 2/3, 4, and 6 was set to 0.5 given the high orientation selectivity (Niell and Stryker, 2008; Durand *et al*., 2016). For layer 5, it was set at 0.8 for the excitatory-to-excitatory connections due to lower orientation selectivity in this layer (Niell and Stryker, 2008; Durand *et al*., 2016).

#### Combining distance-dependent and orientation-dependent adjustments

As can be observed from the above, the scaling can increase the measured connection probability and to ensure our probabilities were never greater than 1, we forced the following condition:

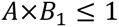

Thus, we used the following algorithm:

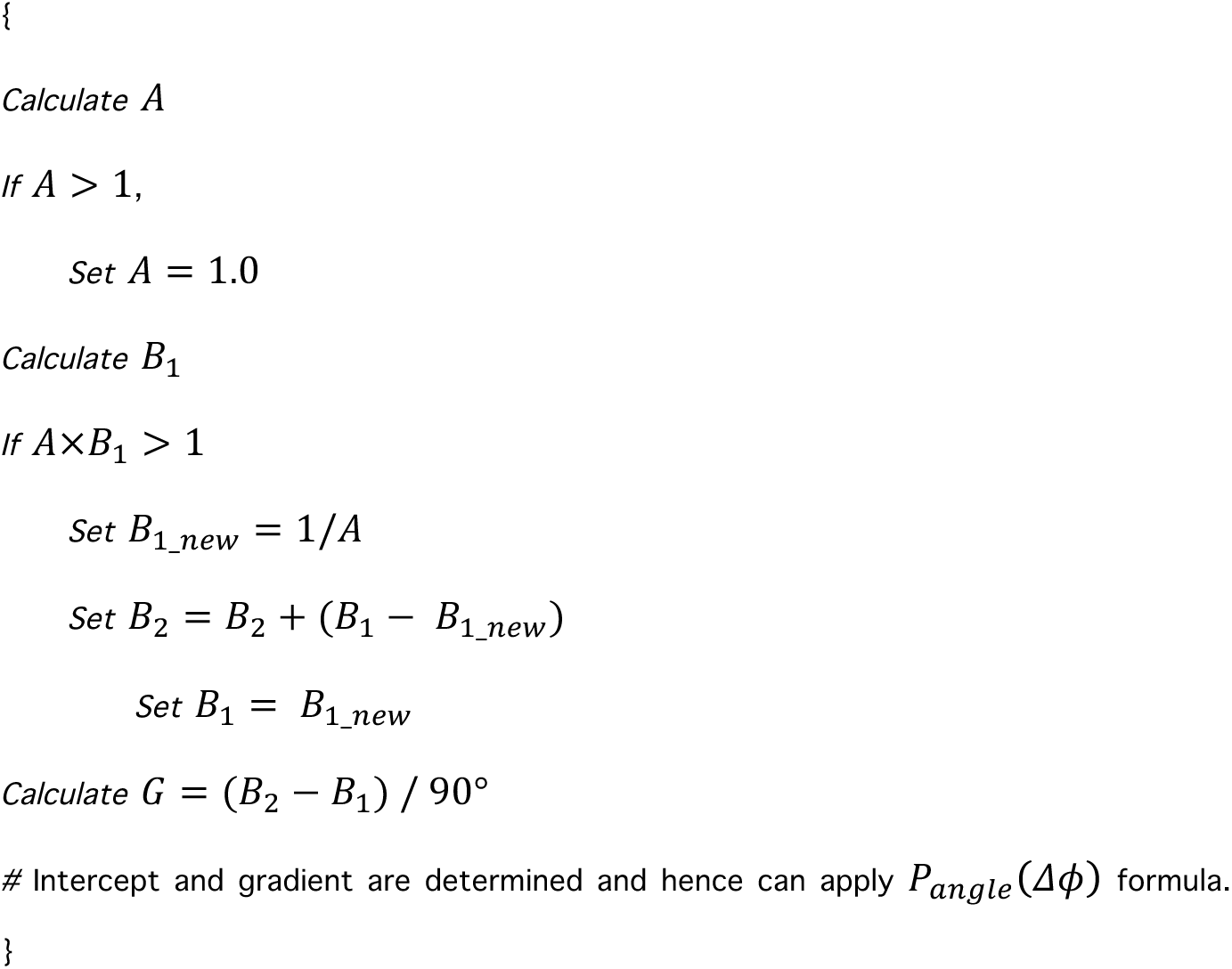

In this formalism (pseudo-code above), if one selects a specific value of *Q* that happens to push the probability values above 1, the worst-case scenario would be that *Q* is rescaled to 1.0 and hence there is no orientation tuning dependence. The trend will never reverse. And this scenario will only occur if there already exists a very high connectivity probability between two cell classes.

With this approach, we have accounted for distance dependence and functional connectivity between the different cell classes in our model. Our next step was to determine the dendritic targeting rules for the biophysically detailed model.

#### Dendritic Targeting for the Biophysical Model

The location of synapses between connected neurons has been demonstrated to have different patterns depending on the neuronal classes (Thomson and Lamy, 2007; Egger *et al*., 2015; Narayanan *et al*., 2015). Although, unfortunately, the available information is sparse, it does delineate trends that may be generalizable, and thus we used these data to implement the rules described below.

##### Excitatory-to-Excitatory Connections

All excitatory-to-excitatory connection avoided the soma and targeted the apical and basal dendrites. For layers 2/3, and 4, the connections were within 200 μm from the soma while for layers 5 and 6, the synapses could form anywhere along the dendrites (Thomson and Lamy, 2007; Egger *et al*., 2015; Narayanan *et al*., 2015). Note that the literature sources are mostly measurements from rat somatosensory cortex. The cortex depth in the rat is approximately 2 mm while our model it is 0.9 mm, and hence we scaled values accordingly.

##### Excitatory-to-Inhibitory Connections

For excitatory-to-inhibitory synapses, both the soma and dendrites could be targeted with no distance limitations (Thomson and Lamy, 2007). This was implemented for all layers and the values were again approximations from the relevant sources.

##### Inhibitory-to-Excitatory Connections (Inhibitory-to-Inhibitory Connections)

For inhibitory-to-excitatory connections we again depended on the data form rat cortex (Thomson and Lamy, 2007). Synapses from the Pvalb class were placed on the soma and dendrites within 50 μm from the soma of any target neuron. Synapses from Sst neurons were placed on the dendrites, 50 μm or further from the soma. Finally, synapses from Htr3a neurons were placed on the dendrites, from 50 μm to 300 μm from the soma. These rules also considered the morphology of neurons in the mouse visual cortex from reconstructions of axons and dendrites (Jiang *et al*., 2015). We assumed for these purposes that Pvalb neurons correspond to basket cells, Sst neurons to Martinotti cells, and Htr3a neurons to Bitufted and Bipolar cells described by (Jiang *et al*., 2015).

Due to the lack of information on inhibitory-to-inhibitory connections, for this class of connections we used rules identical to the inhibitory-to-excitatory connections described above.

##### Layer 1

Finally, for layer 1 neurons, which are Htr3a only in our V1 model, we used the rules below that heavily depended on data from rat neocortex (Jiang *et al*., 2013) and neuron morphology from mouse V1 (Jiang *et al*., 2015), and are similar to other layers due to lack of references with explicit measurements. Our original goal for the model was to project i1Htr3a-to-E2/3 to apical dendrites (no somatic connections) from 50 μm and greater (see below). This is based on distance estimates from the bottom of L1 to upper L2/3 that are approximately 50 μm. This was decided by observing the extent of axonal arbors of L1 (according to Jiang *et al*., 2015, reconstructions). Similarly: i1Htr3a-to-E4 projected to apical dendrites that are 200 μm away from the soma; i1Htr3a-to-E5 projected to apical dendrites that are 300 μm away from the soma and greater; i1Htr3a-to-E6 projected to apical dendrites that are 500 μm away from the soma; i1Htr3a-to-i1Htr3a projected everywhere including soma; i1Htr3a-to-i2/3 projected to basal dendrites from 50 μm and greater. For the other inhibitory layers that project to layer 1, the same rules were used as for within-layer i-to-Htr3a. Finally, excitatory projections to layer 1 were placed on the soma and dendrites with no distance limitations.

Note, however, that during our post-synaptic-potential optimization (see below), we had to change the rules of synaptic placement when L1 was the source onto excitatory cells. Our optimization methodology would create 100 target cells of a specific cell model that receive 1 spike at 0.5 seconds and we would record the generated postsynaptic potential (PSP). The weight would be scaled until we were within 1% of the target PSP. We observed that the when L1 was the source impinging on excitatory cells, the targets sections were so far away that the somatic PSP would reach a maximum and never match the target PSP regardless of how strongly the weight was scaled. This was due to the most distal compartments reaching their maximum membrane deviation that is equal to the reversal potential of the synaptic drive. With these distal compartments being at their maximum, and the attenuation that occurs due to dendritic filtering (recall dendrites in our model are passive), the soma would reach a maximum PSP that did not match our target values.

Thus, to address this issue, we changed the synaptic placement rules for all L1-to-Excitatory neurons so that synapses were placed on dendrites at 50 μm or further from the soma. This is just a highly simplified approximation, but, in terms of reaching closer to the soma than our original rules, it is reasonable since L1 neurogliaform cells are known to bulk release GABA into large volumes and not form well-targeted synapses with post-synaptic cells (Szabadics, Tamás and Soltesz, 2007; Oláh *et al*., 2009; Tremblay, Lee and Rudy, 2016). In the future, employing neuronal models with active dendritic conductances will help alleviate such problems. While a number of such “all-active” biophysical neuronal models are already available in our Allen Cell Types Database (celltypes.brain-map.org/), they are much more computationally expensive than the “perisomatic” biophysical neuronal models with passive dendrites, used here. Furthermore, even the active dendrite models in the Allen Cell Types database have spatially uniform conductances, whereas to avoid the above described problem of distal inputs driving the dendritic voltage to synaptic reversal potential, one would have to include experimentally observed somatodendritic gradients of ion channels (leak, voltage-dependent potassium, HCN) to reduce the input impedance of small diameter dendritic branches. Developing reliable and accurate all-active neuronal models incorporating these gradients for a variety of cell types is thus an important avenue for future work.

Finally note that in our optimization we always let the cells relax to their baseline. Since the resting potential is lower than the reversal potential of the synapses, the single spike at 0.5 seconds would always cause a depolarization. We still used this depolarization level to optimize weights for excitatory PSPs and inhibitory PSPs.

#### Orientation Rule for Synaptic Strength

##### Matching Target Post Synaptic Potentials

The first version of our V1 model (**Figs. 4, 5**) used an orientation-dependent like-to-like rule for synaptic weights of all connection classes: E-to-E, E-to-I, I-to-E, and I-to-I (see Main Text). Since neurons had pre-assigned preferred angles, the connection strength was a function of the difference between the assigned angles of two connected neurons, defined within 90°. The synaptic strength between two cells was then defined as:

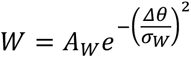

where *Δθ* is the difference between the assigned angles of two neurons and *σ_w_* is the standard deviation set to 50° for all connection classes. Finally, *A_w_* is the weight constant that needed to be determined for every connection class to be matched to Post Synaptic Potential (PSP) targets.

For the biophysical model the units of *W* are in *μS* (defined as the peak conductance), and for GLIF model, in *pA* (see **Synaptic Characteristics**). Since most of the studies used to construct our PSP resource (Connection_strengths.pptx) employed *in vitro* patch-clamp experiments, the data do not distinguish a neuron’s functional preferences, such as preferred angle. Therefore, we assumed the neurons were targeted uniformly and, thus, for optimization we created 100 target cells from every model that were assigned tuning angles with equidistant spacing in the range [0, 360°). We then created a virtual source node for every connection class using the rules described above. The source node would emit 1 spike every 0.5 seconds. We then averaged the post-synaptic responses over all 100 target cells and iteratively updated the weight value (the factor *A_w_* in the equation above) until the mean PSP was within 1% of the target value.

For scaling the weights when the target was a LIF neuron, 1000 source cells were created, each firing at 1Hz from a Poisson distribution. These cells would first target every *biophysical* cell model, using the synaptic weights that were already optimized as described above, and the resulting firing rates due to this input would be calculated. The target firing rate for the LIF neurons were then estimated as the weighted average rate (relative to the proportion of times a model would appear as part of a population). The same source cells (with identical spike times) would then be connected to LIF targets and the firing rate would be matched to within 5% of the desired firing rate.

For inhibitory connections onto the target LIFs, we used the same scaling factors as calculated for their excitatory counterparts. Although not ideal, we chose this route after checking our previous Layer 4 model (Arkhipov *et al*., 2018) and observing that indeed in that previous work the scaling ratios for LIFs for inhibitory input were approximately equal to the scaling ratios of excitatory inputs.

Finally, for the GLIF model, the weights could be calculated analytically based on connection strengths (i.e., PSPs) between the source and target populations (shown in Connection_strengths.pptx) and the mathematical model of the postsynaptic current (i.e., alpha function, see **Synaptic Characteristics**), together with the GLIF model membrane potential dynamics (Teeter *et al*., 2018). Namely, the weights were computed by solving the following equation that describe dynamics in the GLIF model after one spike injection.

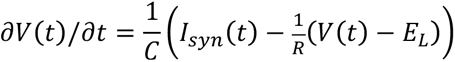

where *V*(*t*) is the membrane potential, *C* is the capacitance of the target neuron, *I_syn_*(*t*) is the alpha-shaped post-synaptic current function with weight *W_GLIF_* (definition in **Synaptic Characteristics**), *R* is the resistance of the target neuron, and *E_L_* is the resting potential. Note that weights in the GLIF model are current based while they are conductance based for the biophysical model. The steps for computing the weight W*_GLIF_* based on the above GLIF model voltage dynamics are:

1. Solving the above dynamic equation to get the analytical solution of membrane potential *V*(*t*);
2. Computing the derivative of the solution of *V*(*t*), i.e., *∂V*(*t*)/*∂t*;
3. Setting *∂V*(*t*)/*∂t* to zero and solving the equation to get the optimal time point *t_max_* at which *V*(*t*)
4. its maximum;
5. Substituting *t_max_* for *t* and the target PSP for *V*(*t*) to the solution of *V*(*t*);
6. Solving the equation generated in 4) to get the weight *W_GLIF_*.

The resultant solution for the weight *W_GLIF_* is

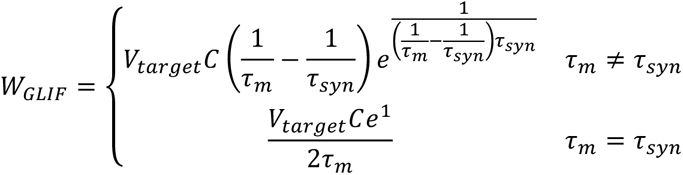

with *V_target_* being the target PSP, *τ_syn_* being the synapse time constant, and *τ_m_* being the membrane time constant.

##### Optimization of Full V1 Models

As described in the Main Text, running simulations after the above optimization did not yield suitable network behaviors in either of our V1 models. Thus, we used an iterative grid search method (Arkhipov *et al*., 2018), where weights were uniformly scaled for every class (e.g. scaling weights of excitatory layer 4 to excitatory layer 5 connections all by the same amount, as one iteration). We searched in discrete increments weight changes across connection classes and selected the best result before moving on to the next connection class (although there was still a need to revisit connections classes during this process). The optimization employed a small training set consisting of two 0.5 second-long simulations: one of gray screen, and the other of a single drifting grating. We aimed to satisfy three criteria: (i) match spontaneous firing rates (gray screen stimulus) to experimental observations, (ii) match peak firing rates for the drifting grating, and (iii) avoid epileptic-like activity where the network would ramp up to have large global bursts and then enter a period of silence until the next very rapid burst. The weight adjustments were kept in a strict range where, for example, the LGN to L4 excitatory weights were not adjusted at all given that they were fit to direct *in vivo* experimental measurements (Lien and Scanziani, 2013). Other LGN connections were restricted to be scaled only in the range [0.5, 2] from the target net input current as those were scaled from optogenetics experiments (Ji *et al*., 2015). The optimization was performed starting from L4 only and adding successive layers one by one (Figs. 4G, S6). First, all interlayer connections were set to zero and only the intra-layer connections in L4 were optimized. Once our criteria were met, we added L2/3 to the optimization, including the interactions between the two layers. This procedure simplified the optimization process even though weights optimized at one step had to be readjusted at the next step (typically minor). This process was continued for layer 5, followed by layer 6, and finally layer 1. During our optimization, the weight scaling was restricted in the range of [0.2, 5]. In the deeper layers (layers 5 and 6), this rule had to be expanded to reach the net adjustment range of [0.12, 18] for the biophysical model and [0.17, 6.0] for the GLIF model. Note that adjusting the synaptic weights in the biophysical model did not translate directly to scaling the PSP (see the Layer 1 description in **Dendritic Targeting for the Biophysical Model**).

#### Optimization with the Direction-Based Rule and Phase Dependence for Synaptic Strength

As described in the Main Text, the next version of our V1 models used a rule for synaptic strengths that was asymmetric with respect to the reversal of direction and included phase dependence, such that the strongest synaptic inputs were sourced from a stripe perpendicular to the preferred direction of the target cell (Figs. 6A, 6B). Once this rule was introduced, the weights needed to be optimized further, as the balance in the network was affected. As a first step, we scaled the recurrent synaptic weights so that the net current (area under the curve, Fig. 6A) became the same as in the previous version of the model (Fig. 4D) for every connection class. However, this was not sufficient, and, thus, we further performed another round of optimization as described in the above section. It turned out that because of the scaling to match the area under the curve, the weights were already close to the correct solution, and we found that these new optimizations required only a few iterations before converging to meet our criteria. For the same reason, here it was not necessary to optimize the models layer-by-layer, and instead the optimization was performed with the full recurrent connectivity. The weight scaling was not constrained to tight limits, however, due to the new synaptic strength profiles that deviated substantially and in a non-linear fashion from those used before.

#### Correcting for Biases Between Horizontal- and Vertical-Preferring Neurons

After finalizing the optimization using the rules above, we noticed biased firing rates in our models, in that vertical drifting gratings evoked higher firing rate relative to horizontal gratings (Fig. 6C). Since this was not observed experimentally and was a result of extra excitatory synaptic drive into vertically preferring neurons (Fig. S11), we adjusted incoming synaptic weights to maintain equal net synaptic drive. The adjustment depends on the cortical magnification factors in the azimuth and elevation dimensions. As described in **Visual Coordinates**, the physical dimensions of each V1 neuron was converted to visual space by a conversion factor of 70 degrees/mm in the azimuth (x-dimension) and 40 degrees/mm in elevation (z-dimension), estimated from experimental reports (Schuett, Bonhoeffer and Hübener, 2002; Kalatsky and Stryker, 2003). To adjust for this asymmetry, we collapsed every neuron’s preferred angle to the quadrant *θ* = [0, 90] and scaled synapses to neurons that preferred horizontal motion (0-degrees) by

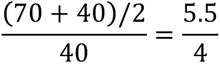

whereas synapses to neurons preferring vertical motion (90-degrees) were scaled by:

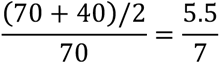

Given these two points, we then fit a linear function to estimate the weight scaling for every intermediate value, resulting in

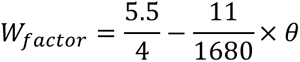

This weight adjustment fixed the bias (Figs. 6C, S11) and resulted in horizontal-preferring neurons having a heavier tail of the incoming synaptic strength distribution than vertical-preferring neurons (Fig. 6E). Finally, due to our highly non-linear V1 models, this adjustment resulted in deviations from our target optimization firing rates. Thus, a small amount of grid search tuning was needed again to match our target criteria.

#### Synaptic Characteristics

The synaptic mechanisms used for the biophysical model were as in the L4 model (Arkhipov *et al*., 2018). The synapses were bi-exponential (using NEURON’s Exp2Syn mechanism) with a reversal potential of −70 mV for inhibition and 0 mV for excitation. The weights’ units are in *μS* (peak conductance). The tau1 and tau2 constants for the mechanism were 2.7 ms and 15 ms for inhibitory-to-excitatory synapses, 0.2 and 8 ms for inhibitory-to-inhibitory synapses, 0.1 ms and 0.5 ms for excitatory-to-inhibitory synapses, and 1 ms and 3 ms for excitatory-to-excitatory connections. Note that these are not the somatic temporal characteristics, but time constants at the synaptic location; the PSP shape at the soma depends on dendritic location of the synapse and membrane dynamics.

For the GLIF model, postsynaptic current-based synaptic mechanisms were used with dynamics described by an alpha-function:

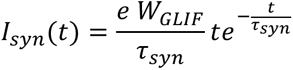

Where *I_syn_* is the postsynaptic current, *τ_syn_* is the synaptic port time constant, and *W_GLIF_* is the input connection weight. This function was normalized such that a post-synaptic current with synapse weight *W_GLIF_* = 1.0 has an amplitude of 1.0 *pA* at the peak time point of *t* = *τ_syn_*. The *τ_syn_* constants for the mechanisms were 5.5 ms for excitatory-to-excitatory synapses, 8.5 ms for inhibitory-to-excitatory synapses, 2.8 ms for excitatory-to-inhibitory synapses, and 5.8 ms for inhibitory-to-inhibitory connections, which were extracted from LIF models in the L4 model (Fig. S2B of (Arkhipov *et al*., 2018)).

## Visual Stimuli

The visual stimuli used in our simulations were identical to those used for the experiments we compare to (except the looming disk that had no experimental counterpart). Each simulation included a 500 ms interval of gray screen in the beginning, which was then followed by a single trial of presentation of the stimulus.

### Drifting Gratings

For the drifting grating stimuli, we used sinusoidal gratings with a spatial frequency of 0.04 cycles per degree with a temporal frequency of 2Hz (for 2.5 seconds after the gray-screen). All stimuli were run for 10 trials for every direction of motion (8 sampled directions with increments of 45 degrees) at 80% contrast (for both the experiments and the models). Although the experimental data from mice (see below) included more temporal and spatial frequencies, we restricted our analysis to match the drifting gratings used in our simulations.

### Flashes

The flash stimuli (10 trials) consisted of: 500 ms of gray screen, followed by 250 ms of white screen (ON-flash), returning to a gray screen for 1000 ms, then another 250 ms of black screen (OFF-flash), and a final gray screen for 500 ms). The contrast was at 80% (to match experiments). We also conducted simulations with full-contrast flashes (100%), and the models were stable and produced results very similar to the 80% contrast case (data not shown).

### Natural Movies

We tested our models on a clip (10 trials) from one of the natural movies (*Touch of Evil*, directed by Orson Welles) used in the Allen Brain Observatory (de Vries *et al*., 2019). The 2.5 seconds shown were matched between the model and experiment.

### Looming Disk

The looming stimulus is a growing black disk (circle) on a gray background. When the maximum circle size (radius of 25 degrees) is reached, the circle disappears and grows again. This is repeated four times throughout the 2.5 second stimulus presentation (625 ms duration for every repetition).

### Data Analysis

#### Firing Rates

The firing rates were estimated from all trials of a simulation. Since all simulations started with a 500ms gray-screen period followed by the stimulus, the firing rate is estimated using the stimulus duration without these first 500 ms (that is, 2500 ms for a drifting grating or a natural movie). Thus, the firing rate for a neuron in a trial was calculated by dividing the total number of spikes after the gray screen by the stimulus duration (2500 ms). Some metrics required time-dependent firing rates that are described below. For the OSI and DSI metrics, to avoid noise from very sparsely firing neurons that could yield spurious OSI/DSI values of 1.0, we required that neurons’ firing rates at their preferred drifting grating direction be greater than 0.5 Hz.

#### Orientation Selectivity Index (OSI)

The OSI metric computed is also referred to as the global Orientation Selectivity Index, as it takes into account the response of a neuron in all directions tested (not just the preferred and orthogonal). The OSI is calculated as:

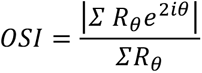

where *R*_*θ*_ is the mean firing rate response to a drifting grating of angle *θ*.

#### Direction Selectivity Index (DSI)

Similar to the OSI metric, the DSI also considered responses in all directions of drifting gratings shown (sometimes referred to as the global Direction Selectivity Index). The DSI is calculated as:

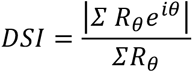

where *R*_*θ*_ is the mean firing rate response to a drifting grating of angle *θ*.

#### Response at Preferred Direction

The plots quantifying neurons’ response at their preferred direction report the mean firing rate values based on the largest mean response (across trials) over all 8 directions tested.

#### Signal Correlations, Noise Correlations, and Correlation of Signal and Noise Correlations

We computed the signal correlation as the Pearson correlation coefficient between the trial-averaged spike counts for each pair of neurons (Arkhipov *et al*., 2018). For natural movies, we computed the correlation for binned spike counts in non-overlapping windows of length 50 ms. For gratings, the correlation was computed over the spike counts in 8 different orientations. The noise correlation was computed as the Pearson correlation coefficient between single-trial spike counts for each pair of neurons, and then averaged over stimuli conditions (8 orientations for gratings and non-overlapping 50 ms windows for natural movies). To compute the correlation of signal and noise correlations for a single experimental mouse, we calculate the Pearson correlation coefficent between the noise correlation and signal correlation metrics already calculated. Since we have many mice (20 for drifting gratings, 7 for natural movies), we subsample neurons within 150 μm from the center mini-column of the models to match the number of neurons per mouse. The subsampling is without replacement. We restricted the sampling near the center of the models to match experimental Neuropixels recording as much as possible.

#### Lifetime and Population Sparsity

Lifetime sparsity for each neuron was computed using the following definition (Vinje and Gallant, 2000):

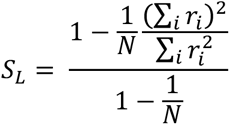

where N is the number of stimulus conditions and r_i_ is the trial-averaged spike count for stimulus condition i (de Vries *et al*., 2019). To compute the population sparsity, we used the same equation, but where N is the total number of neurons in the population and r_i_ is the average spike-count of neuron i over all stimulus conditions (de Vries *et al*., 2019).

#### Similarity Score

A similarity score was developed to compare the distribution of all excitatory neurons in the models with all regular spiking neurons recorded experimentally as well as for Pvalb neurons in the models with fast-spiking neurons from the same Neuropixels recording. The score compares any two distributions and does not require a normality assumption nor that both distributions have an equal number of samples. Moreover it can be applied to any metric and we use it here to compare OSI, DSI, and the firing rate distributions of the models with experiments. The metric uses the *D* statistic from a Kolmogorov–Smirnov test that calculates the distance between the cumulative distributions of two samples and is bounded in the range [0, 1]. Since we are interested in similarity in this work and matching distributions, this was converted to a similarity score, **S** = 1 − *D*. Fig. S5 illustrates how for two different distributions **S** is close to 0, whereas for two similar distributions it approaches 1.

#### Electrophysiological Recordings

##### Animal preparation

All experimental procedures were approved by the Allen Institute for Brain Science Institutional Animal Care and Use Committee. Five weeks prior to the experiment, mice were anesthetized with isoflurane, and a metal headframe with a 10-mm circular opening was attached to the skull with Metabond. In the same procedure, a 5-mm-diameter craniotomy and durotomy was drilled over left visual cortex and sealed with a circular glass coverslip. Following a 2-week recovery period, a visual area map was obtained through intrinsic signal imaging (Juavinett *et al*., 2017). Mice with well-defined visual area maps were gradually acclimated to the experimental rig over the course of 12 habituation sessions. On the day of the experiment, the mouse was placed under light isoflurane anesthesia for ∼40 min to remove the glass window, which was replaced with a 0.5 mm thick plastic window with laser-cut holes (Ponoko, Inc., Oakland, CA). The space beneath the window was filled with agarose to stabilize the brain and provide a conductive path to the silver ground wire attached to the headpost. Any exposed agarose was covered with 10,000 cSt silicone oil, to prevent drying. Following a 1-2 hour recovery period, the mouse was head-fixed on the experimental rig. Up to six Neuropixels probes coated in CM-DiI were independently lowered through the holes in the plastic window and into visual cortex at a rate of 200 µm/min using a piezo-driven microstage (New Scale Technologies, Victor, NY). When the probes reached their final depths of 2,500–3,500 µm, each probe extended through visual cortex into hippocampus and thalamus. Only data obtained from V1 was included in this study. In total, data from 20 mice were used for the drifting gratings analysis (one experiment per mouse) and 7 mice for the natural movie and flash analysis.

##### Data acquisition system

Recordings were performed in awake, head-fixed mice allowed to run freely on a rotating disk. During the recordings, the mice passively viewed a battery of visual stimuli, including local drifting gratings (for receptive field mapping), full-field flashes, drifting gratings, static gratings, natural images, and natural movies, with the same parameters as those from the Allen Brain Observatory (de Vries *et al*., 2019). All spike data were acquired with Neuropixels probes (Jun *et al*., 2017) with a 30-kHz sampling rate and recorded with the Open Ephys GUI (Siegle *et al*., 2017). A 300-Hz analog high-pass filter was present in the Neuropixels probe, and a digital 300-Hz high-pass filter (3rd-order Butterworth) was applied offline prior to spike sorting.

##### Data preprocessing

Spike times and waveforms were automatically extracted from the raw data using Kilosort2 (github.com/mouseland/kilosort2). Kilosort2 is a spike-sorting algorithm developed for electrophysiological data recorded by hundreds of channels simultaneously. It implements an integrated template matching framework for detecting and clustering spikes, rather than clustering based on spike features, which is commonly used by other spike-sorting techniques. After filtering out units with “noise” waveforms using a random forest classifier trained on manually annotated data, all remaining units were packaged into Neurodata Without Borders format (Teeters *et al*., 2015) for further analysis.

##### Neuronal Classification

Regular spiking (RS) neurons and fast spiking (FS) neurons were determined by the duration of the spike (time between trough and peak of the waveform). The duration of the spikes showed a bimodal distribution (Hartigan dip test, p=0.004), with a dip at 0.4 ms. We classified a neuron as RS if its duration was > 0.4 ms, and otherwise FS (Fig. S4). In total we had 328 L6 RS neurons, 72 L6 FS neurons, 419 L5 RS neurons, 80 L5 FS neurons, 294 L4 RS neurons, 49 L4 FS neurons, 251 L23 RS neurons, 49 L23 FS neurons, and 81 L1 neurons.

