## Supplemental Information for "Systematic Integration of Structural and Functional Data into Multi-Scale Models of Mouse Primary Visual Cortex"

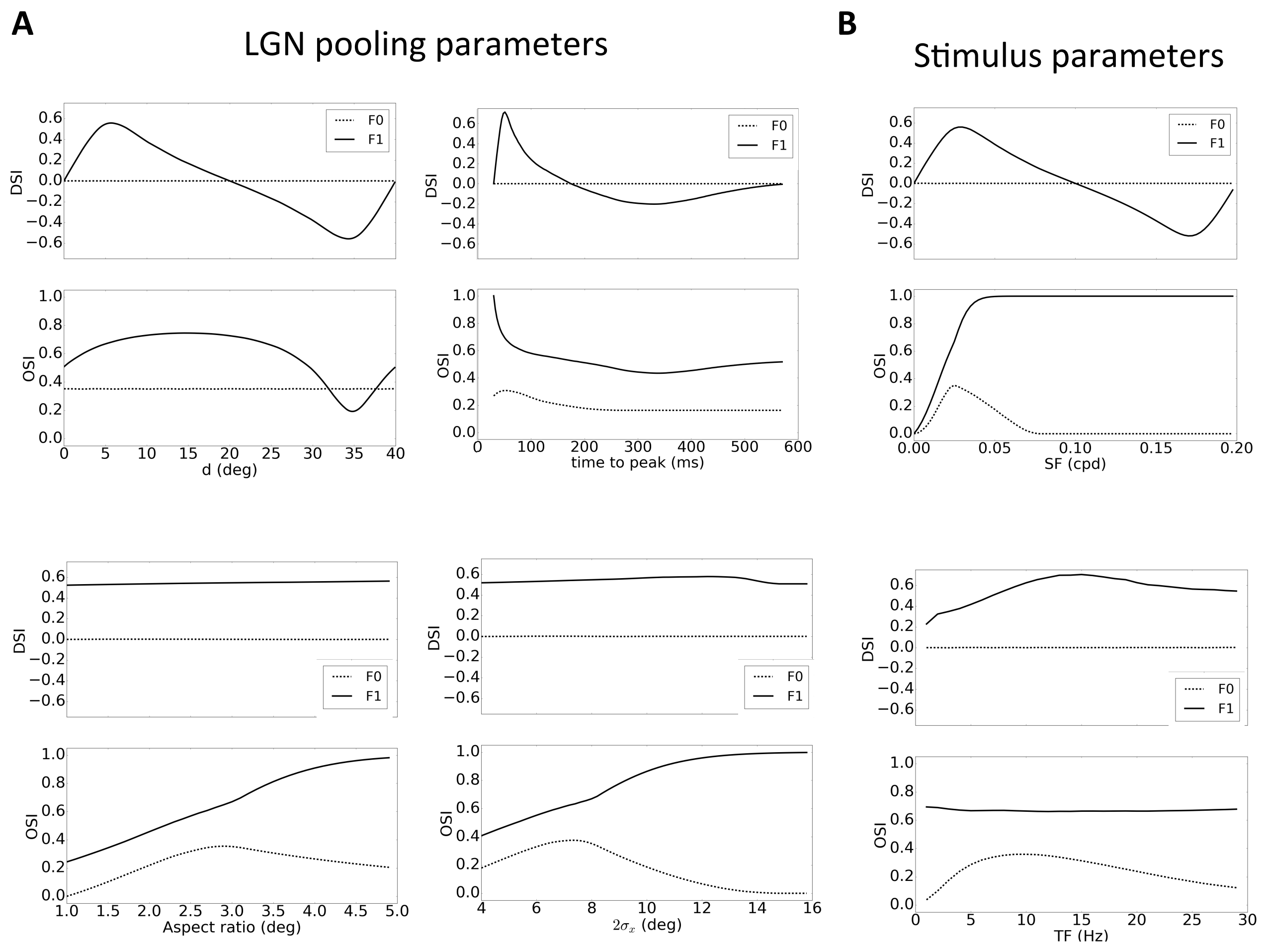


**Figure S1:** The OSI and DSI values for the F1 and F0 components of the theoretically estimated aggregate LGN input into a V1 cell, in response to a drifting grating stimulus as a function of the LGN pooling parameters and stimulus parameters. **A:** Influence of the following LGN pooling parameters on OSI and DSI: distance separation, $d$ (top-left), time-to-peak of the sustained subfield (top-right; time-to-peak of the transient subfield is fixed), ellipse aspect ratio (major axis to minor axis) (bottom-left), and size of the subfields (widths with constant aspect ratio) (bottom-right). **B:** Stimulus parameters: the spatial frequency (top) and temporal frequency (bottom) of drifting gratings. The filters used in (A) and (B) for the two LGN subfields are a sustained ON preferring TF=8 Hz and a transient OFF preferring TF=8 Hz; the remaining parameter values (unless varied on the panel) are: $d$ = 5 degrees, ellipse aspect ratio = 3.0, ellipse minor axis = 4.0 degrees. The grating parameters (unless varied on the panel) were SF = 0.025cpd and TF = 8Hz.

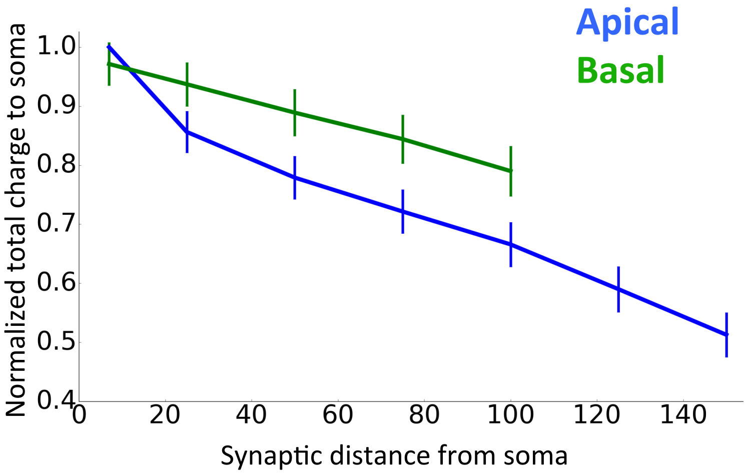


**Figure S2:** Total charge conductance to soma as a function of dendritic placement. Charge was calculated as the integral of total current in response to a drifting grating. All the plots are averaged over all 37 excitatory models used in layer 4 of the V1 biophysical model. The models, individually, show an exponential decay with distance due to the passive dendrites as expected.

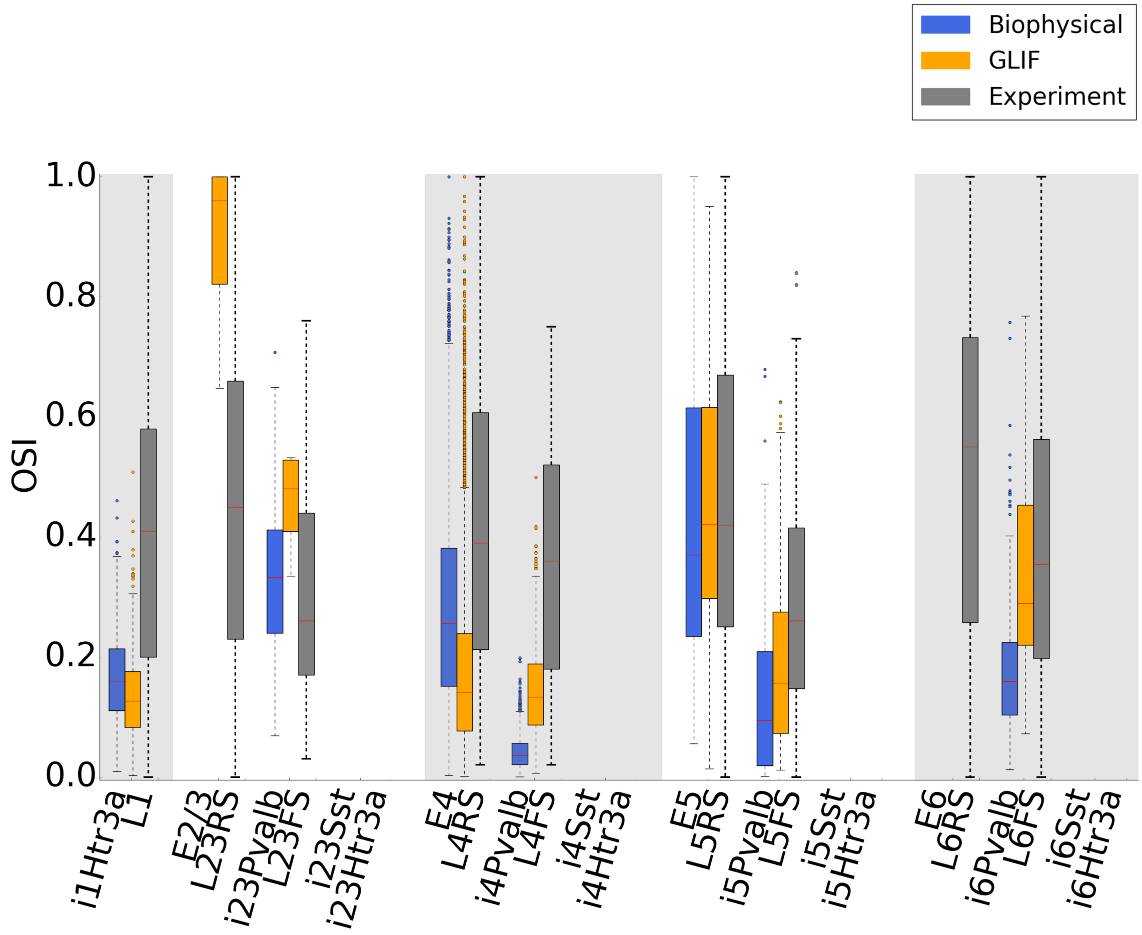


**Figure S3:** OSI values for both V1 models (biophysical and GLIF) for the LGN-only case (i.e., only LGN inputs are active, and all other connections are removed) across neuronal classes, compared to *in vivo* measurements (in awake mice without perturbations, i.e., with intact connectivity). The experimental values are included only as a reference. Note that OSI values can be high in the models as a consequence of very low firing rates due to lack of recurrence (Figure 3).

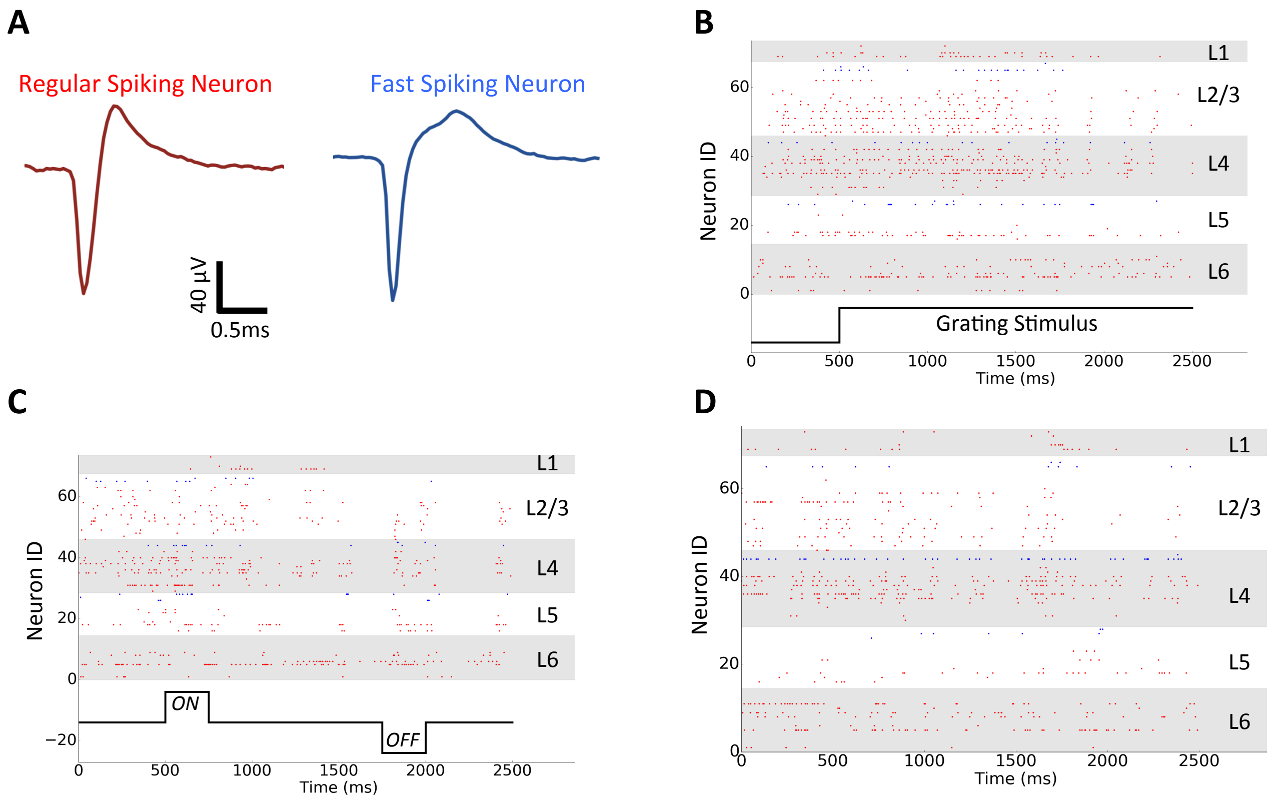


**Figure S4:** Examples of electrophysiological recordings from mouse V1 using a Neuropixels probe from a single experiment (see Methods). In total 27 experiments were aggregated for our analysis. **A:** Spike waveforms from a regular-spiking neuron and a fast-spiking neuron. **B:** Raster plot example in response to a drifting grating from a single recording. **C:** Raster plot example in response to ON and OFF flashes. **D:** Raster plot example in response to a natural movie. The movie portion used in the experiments was 120 seconds long, whereas for simulations we used a 2.5-second clip from the experimental stimulus, to reduce computing expense. For the analysis comparing responses between the models and experiment, we used the activity recorded only for this 2.5-second clip (i.e., exactly matching the frames between the models and experiment). Here, responses to the clip are shown for a single experimental trial (since the clip is a part of the bigger movie, no gray screen period is present).

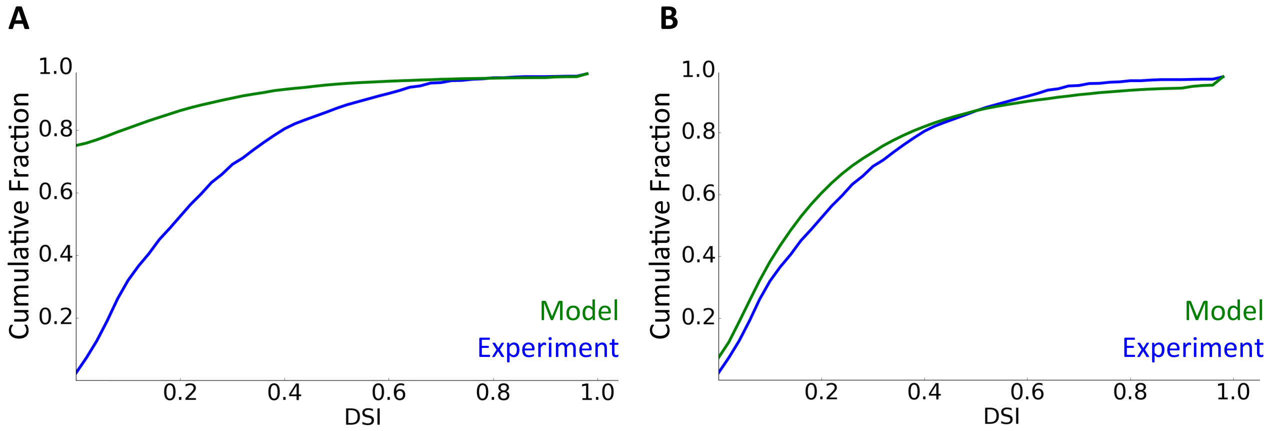


**Figure S5:** Two example distributions to demonstrate the similarity score, $\mathbf{S}$, used. **A:** DSI distribution between the biophysical V1 model that only has LGN input (no recurrent connections) and the experimental data. Here the similarity score is $\mathbf{S}$ = 0.24. **B:** DSI distribution between the final V1 biophysical model and the experimental data. Here the similarity score is $\mathbf{S}$ = 0.92.

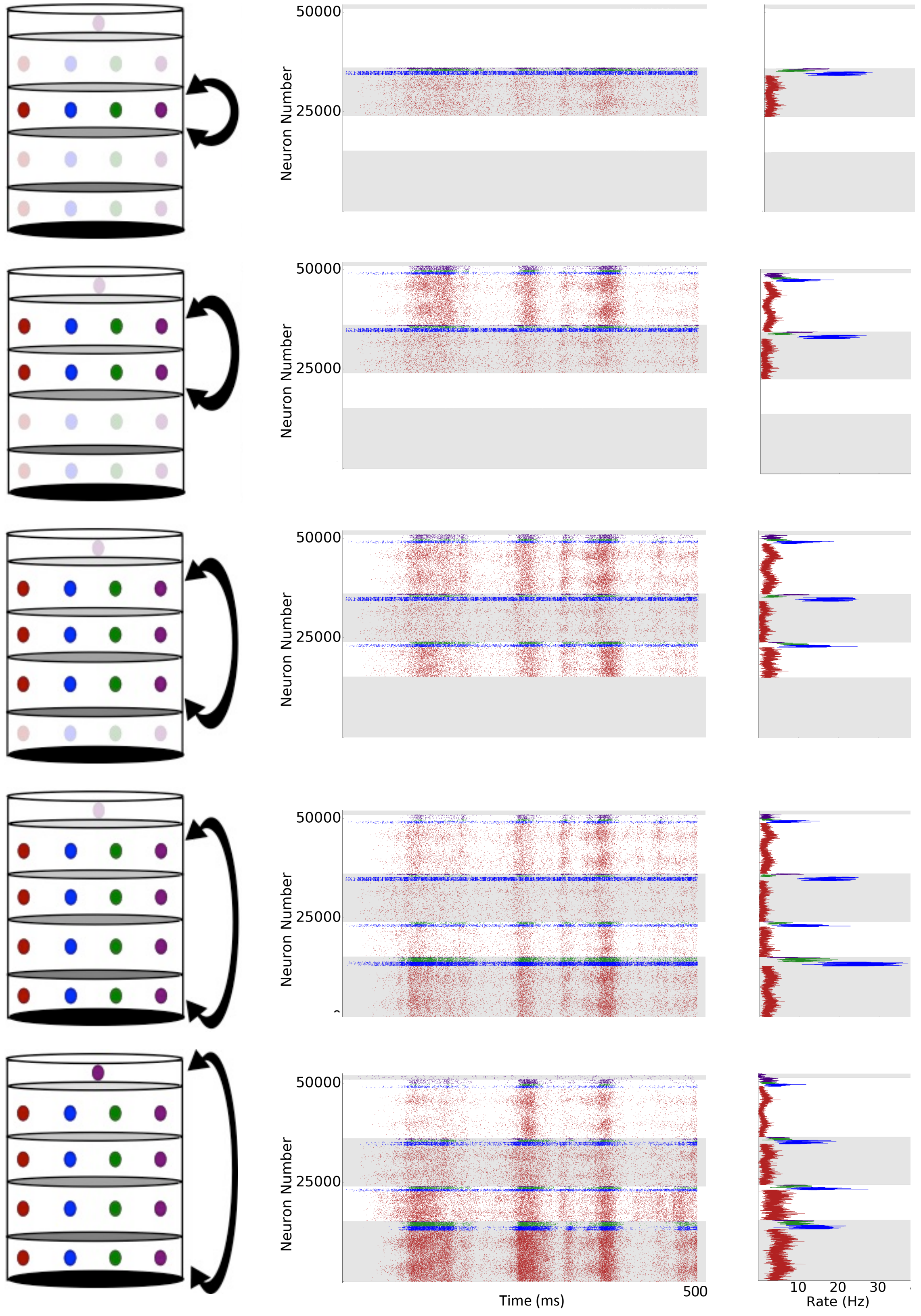


**Figure S6:** Schematic of the optimization procedure with example raster plots and firing rates at every stage. The stimulus is a 500ms drifting grating that was used for the optimization process.

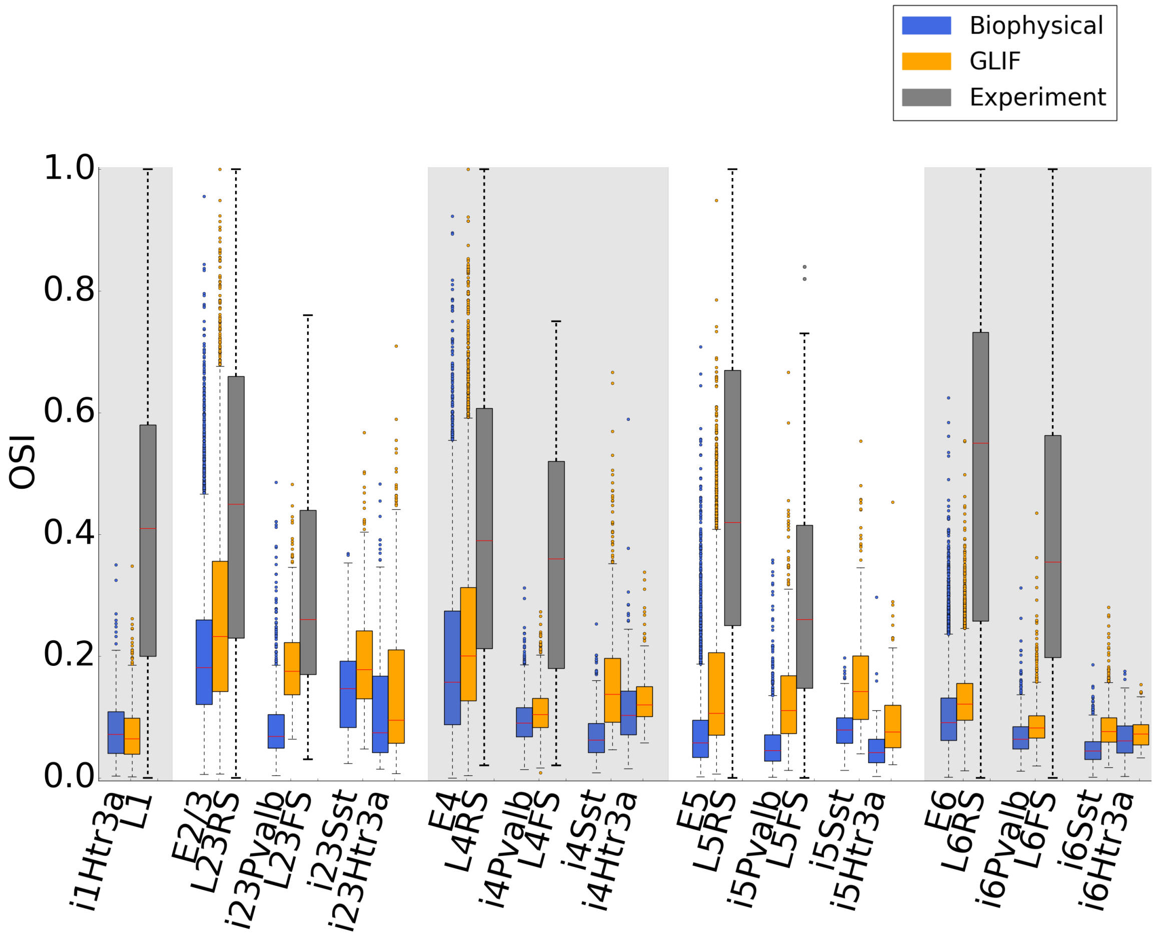


**Figure S7:** OSI values for both V1 models (biophysical and GLIF) across neuronal classes, compared to *in vivo* measurements, for the models with full connectivity using the orientation dependent like-to-like rules for both connection probability and synaptic strengths (Figure 5).

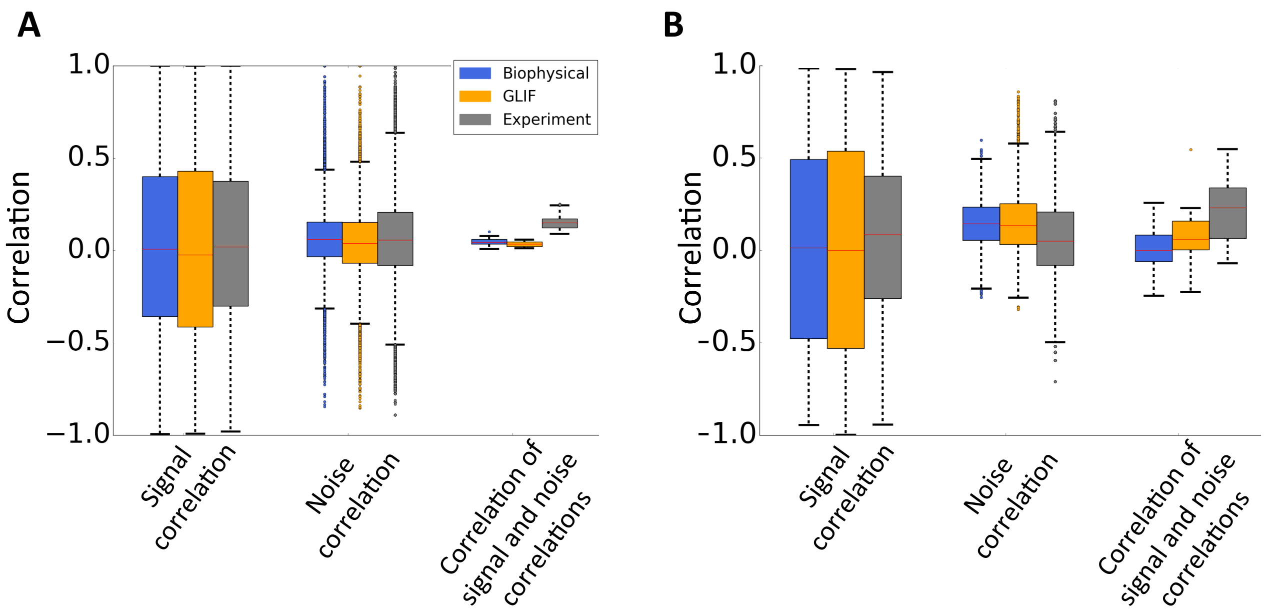


**Figure S8:** Quantification of signal correlation, noise correlation, and correlation of signal and noise correlation for both V1 models (biophysical and GLIF) compared to *in vivo* measurements for the models with full connectivity using the orientation dependent like-to-like rules for both connection probability and synaptic strengths (Figure 5). **A:** For excitatory neurons (selected as regular spiking from experimental data). **B:** For parvalbumin neurons (selected as fast spiking from experimental data).

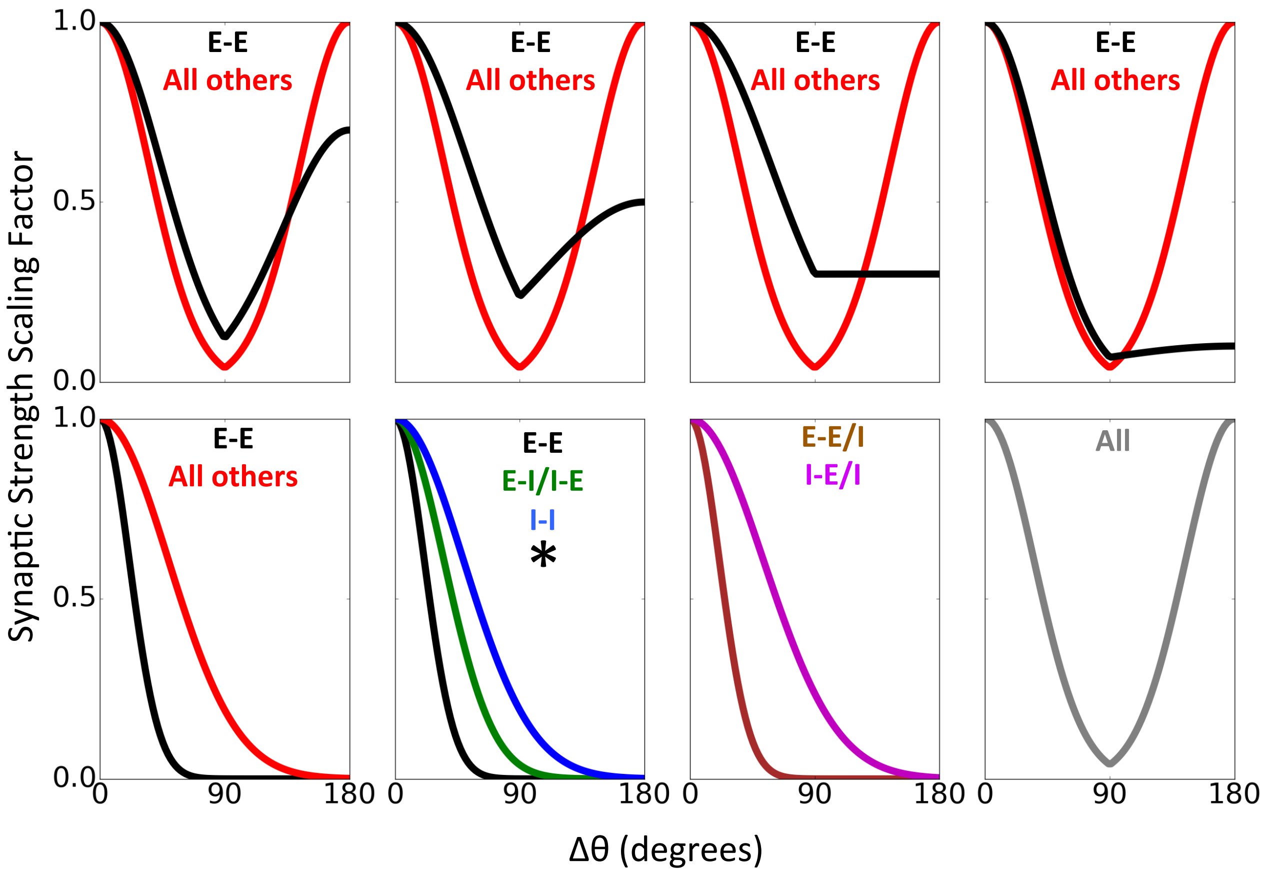


**Figure S9:** The eight synaptic strength rules tested before finalizing the models (Figures 6, 7). Note that multiple sets of parameters were tested for each rule, primarily using the GLIF model (over 100 variations). The final rule used is marked with an asterisk (see Fig. 6A for details). Given that multiple different values of parameters (and rules) result in networks with robust levels of direction selectivity, we cannot reliably choose a single 'optimal' set. The set (Fig. 6A) we use should be considered a representative example among possible solutions.

The connection groups varied are E-to-E, E-to-I, I-to-E, and I-to-I. For all red curves that indicate "all others", they refer to the latter three connection groups. For the top row from left to right, the rules for the E-to-E connections are:

$$\text{Scaling factor}_1 = \begin{Bmatrix} e^{-\left(\frac{\Delta\theta}{62^\circ}\right)^2} & \Delta\theta \leq 90^\circ \\ 0.7 * e^{-\left(\frac{\Delta\theta}{75.7^\circ}\right)^2} & \Delta\theta > 90^\circ \end{Bmatrix}$$

$$\text{Scaling factor}_2 = \begin{Bmatrix} e^{-\left(\frac{\Delta\theta}{75^\circ}\right)^2} & \Delta\theta \leq 90^\circ \\ 0.5 * e^{-\left(\frac{\Delta\theta}{104.1^\circ}\right)^2} & \Delta\theta > 90^\circ \end{Bmatrix}$$

$$\text{Scaling factor}_3 = \begin{cases} e^{-\left(\frac{\Delta\theta}{82^\circ}\right)^2} & \Delta\theta \leq 90^\circ \\ 0.3 * e^{-\left(\frac{\Delta\theta}{125.8^\circ}\right)^2} & \Delta\theta > 90^\circ \end{cases}$$

$$\text{Scaling factor}_4 = \begin{cases} e^{-\left(\frac{\Delta\theta}{55^\circ}\right)^2} & \Delta\theta \leq 90^\circ \\ 0.1 * e^{-\left(\frac{\Delta\theta}{63.9^\circ}\right)^2} & \Delta\theta > 90^\circ \end{cases}$$

The scaling factors for "All other" connection groups in the top row of Fig. S9, as well as bottom-right panel in Fig. S9, follow the equation:

$$\text{Scaling factor} = \begin{cases} e^{-\left(\frac{\Delta\theta}{50^\circ}\right)^2} & \Delta\theta \leq 90^\circ \\ e^{-\left(\frac{\Delta\theta-180}{50^\circ}\right)^2} & \Delta\theta > 90^\circ \end{cases}$$

The scaling factors in the remaining three panels in the bottom row of Fig. S9 follow basic Gaussian profiles with different values for the standard deviation (values tested in the range [10°, 90°]).

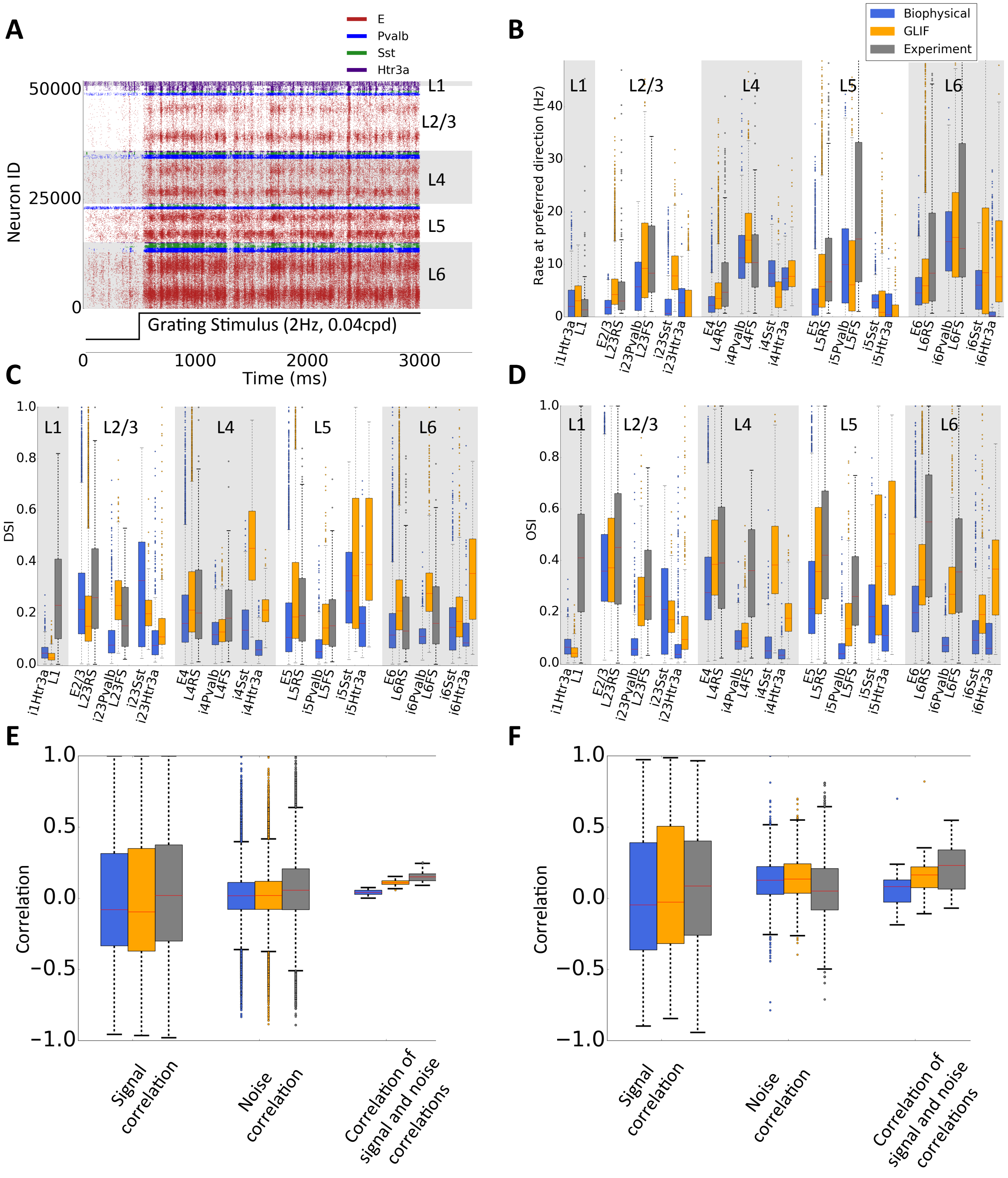


**Figure S10:** Simulated responses to drifting gratings for the V1 model using rules from Fig. 6A, 6B, before correction for the cortical magnification factor asymmetry. **A:** Example raster plot in response to a drifting grating. **B:** Peak firing rates; **C:** DSI; **D:** OSI for V1 models and *in vivo* recordings. **E,F:** Quantification of signal correlation, noise correlation, and correlation of signal and noise correlation for excitatory neurons (**E**) and parvalbumin neurons (**F**).

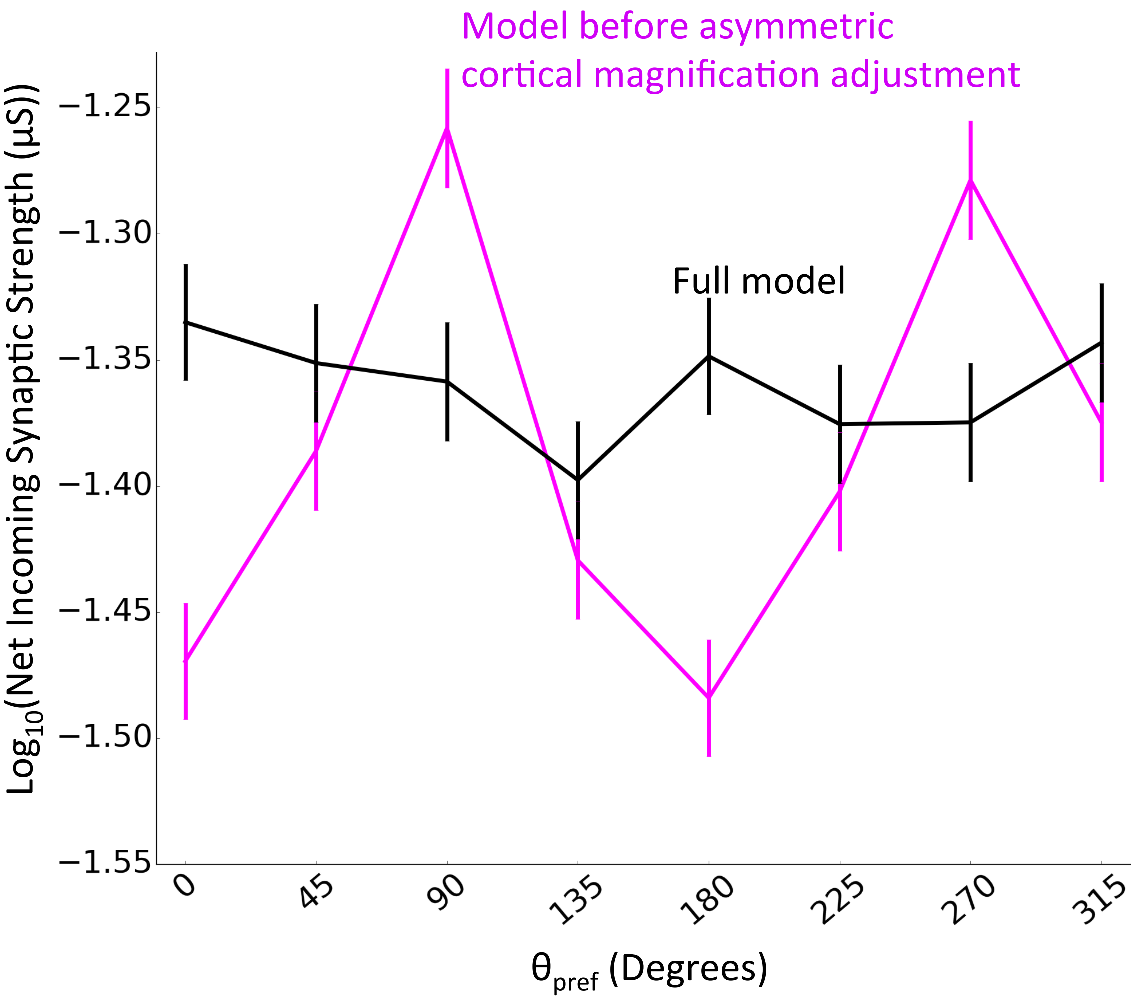


**Figure S11:** Net incoming synaptic strength for excitatory neurons that are ± 3-degrees of their assigned preferred angle (median and s.e.m) in the V1 model that uses rules from Figs. 6A, B, before (Fig. S10) and after (Fig. 7) correction for the cortical magnification factor asymmetry.

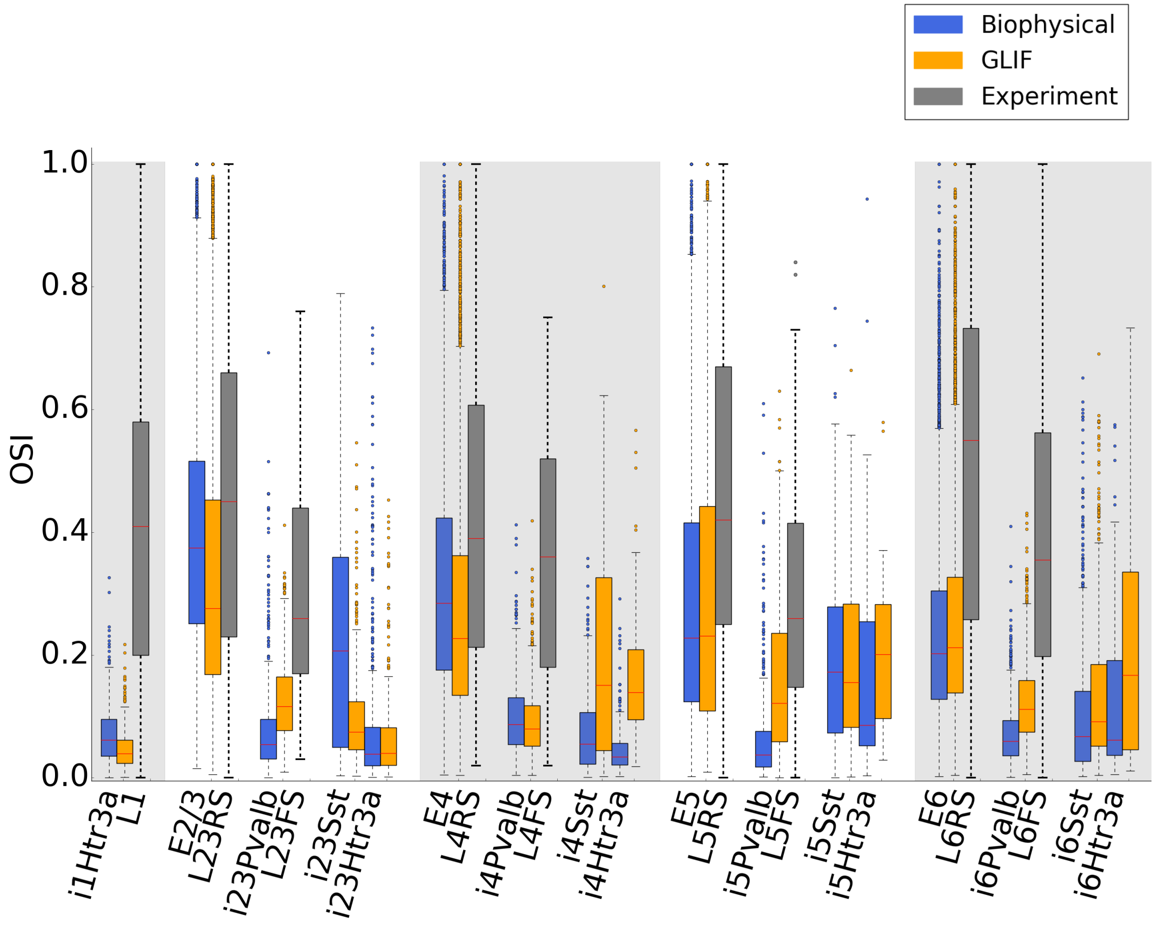


**Figure S12:** OSI values across neuronal classes for the final versions of both V1 models, compared to *in vivo* measurements.

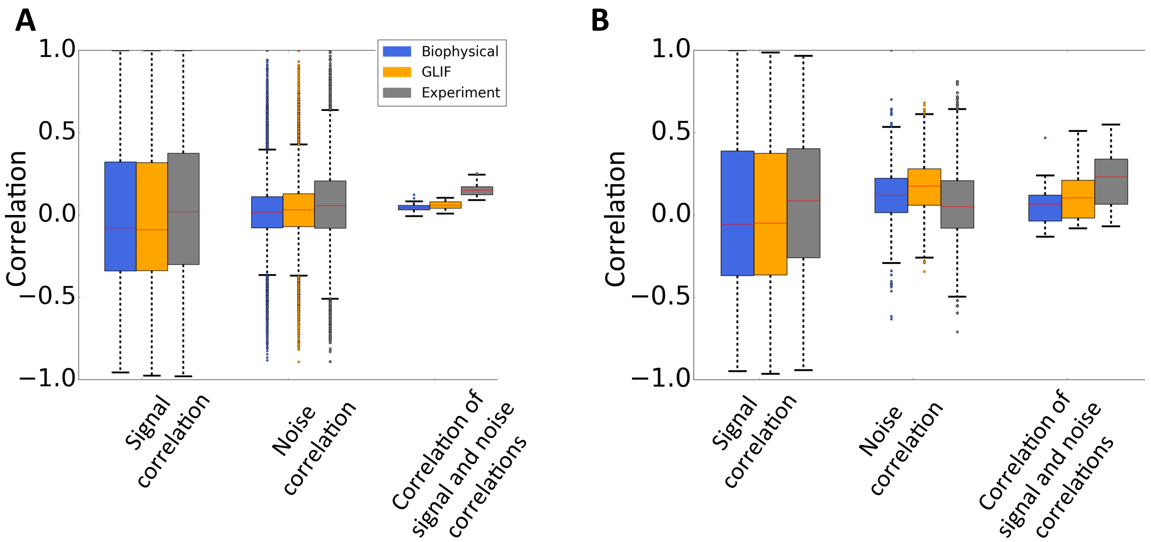


**Figure S13:** Quantification of signal correlation, noise correlation, and correlation of signal and noise correlation for both V1 models (biophysical and GLIF) compared to *in vivo* measurements for the final versions. **A:** For excitatory neurons (selected as regular spiking from experimental data). **B:** For parvalbumin neurons (selected as fast spiking from experimental data).

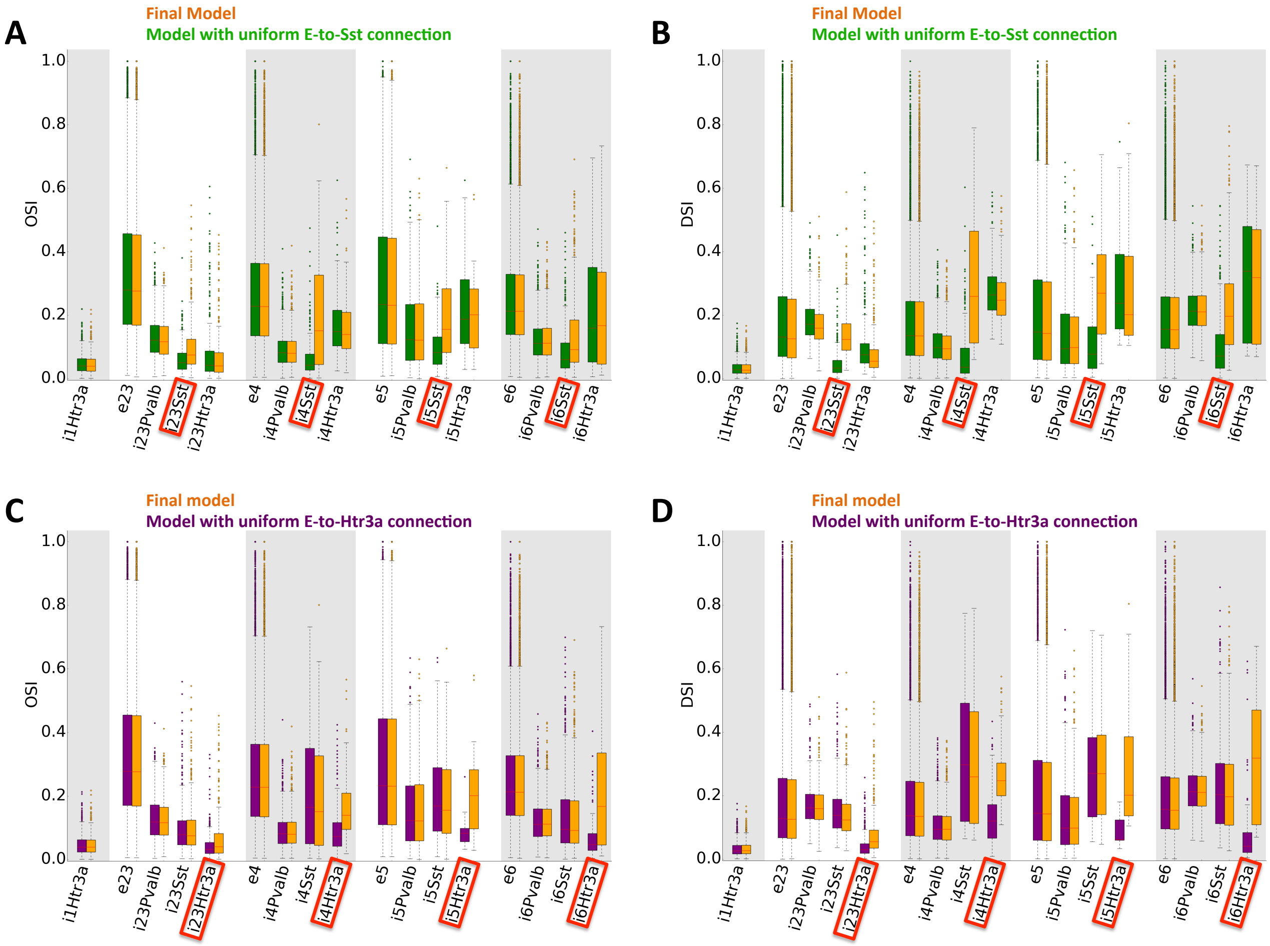


**Figure S14:** Removing the functional connectivity rules of E-to-Sst or E-to-Htr3a results in a loss of direction and orientation selectivity of the target Sst and Htr3a neurons. **A,B:** Simulations results for when the E-to-Sst connections are uniform (no dependence on direction preference of source and target neurons). A prominent decrease in OSI (A) and DSI (B) across all layers is observed for the Sst neurons (red boxes). **C,D:** Simulations results for when the E-to-Htr3a connections are uniform. A prominent decrease in OSI (C) and DSI (D) is seen for the Htr3a neurons (red boxes). All simulation results are from the GLIF V1 network model.

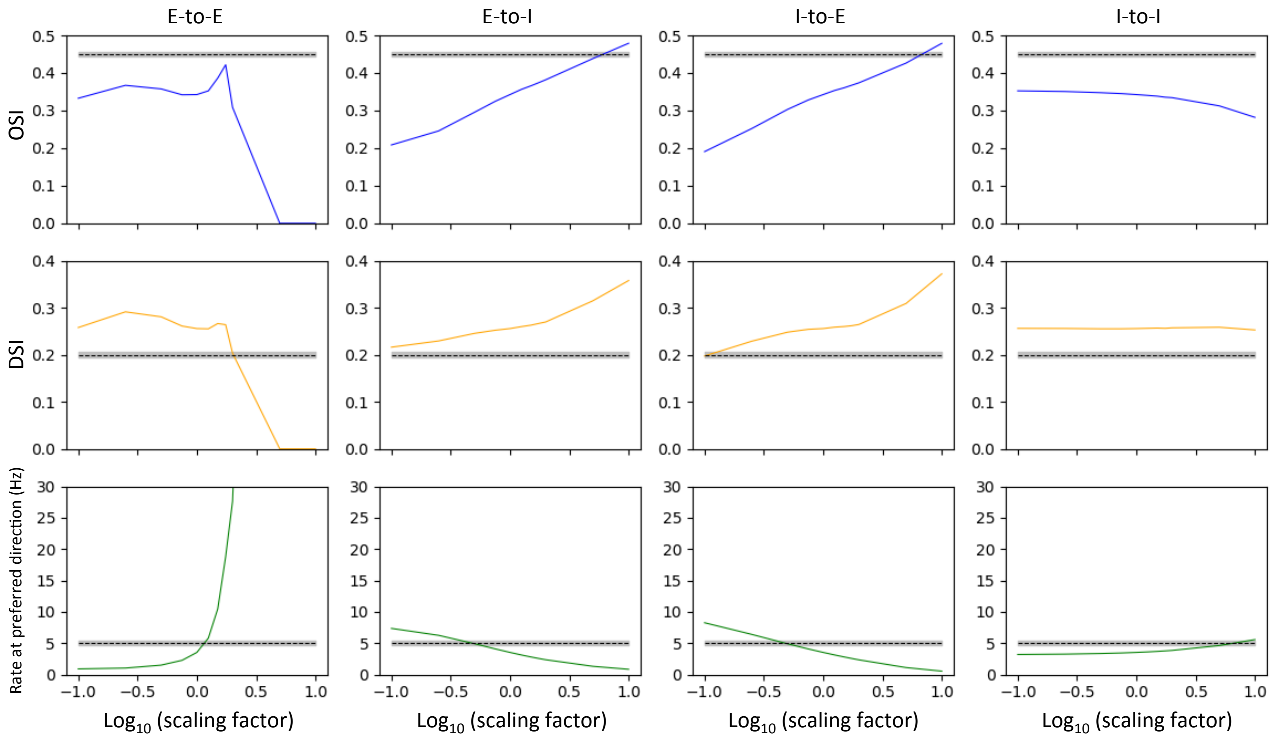


**Figure S15:** Sensitivity analysis for the final model quantifying the responses of excitatory neurons. The four major classes of connections (E-toE, E-to-I, I-to-E, and I-to-I) were scaled uniformly in the GLIF model for a range of values (from 0.1 to 10). These scaling factors are used as the x-axes of all plots. Every column represents one class of connections scaled and every row shows a different metric (OSI, DSI, or Rate at the preferred drifting grating direction). The horizontal lines are median experimental values with the shaded region showing the standard error of the mean.

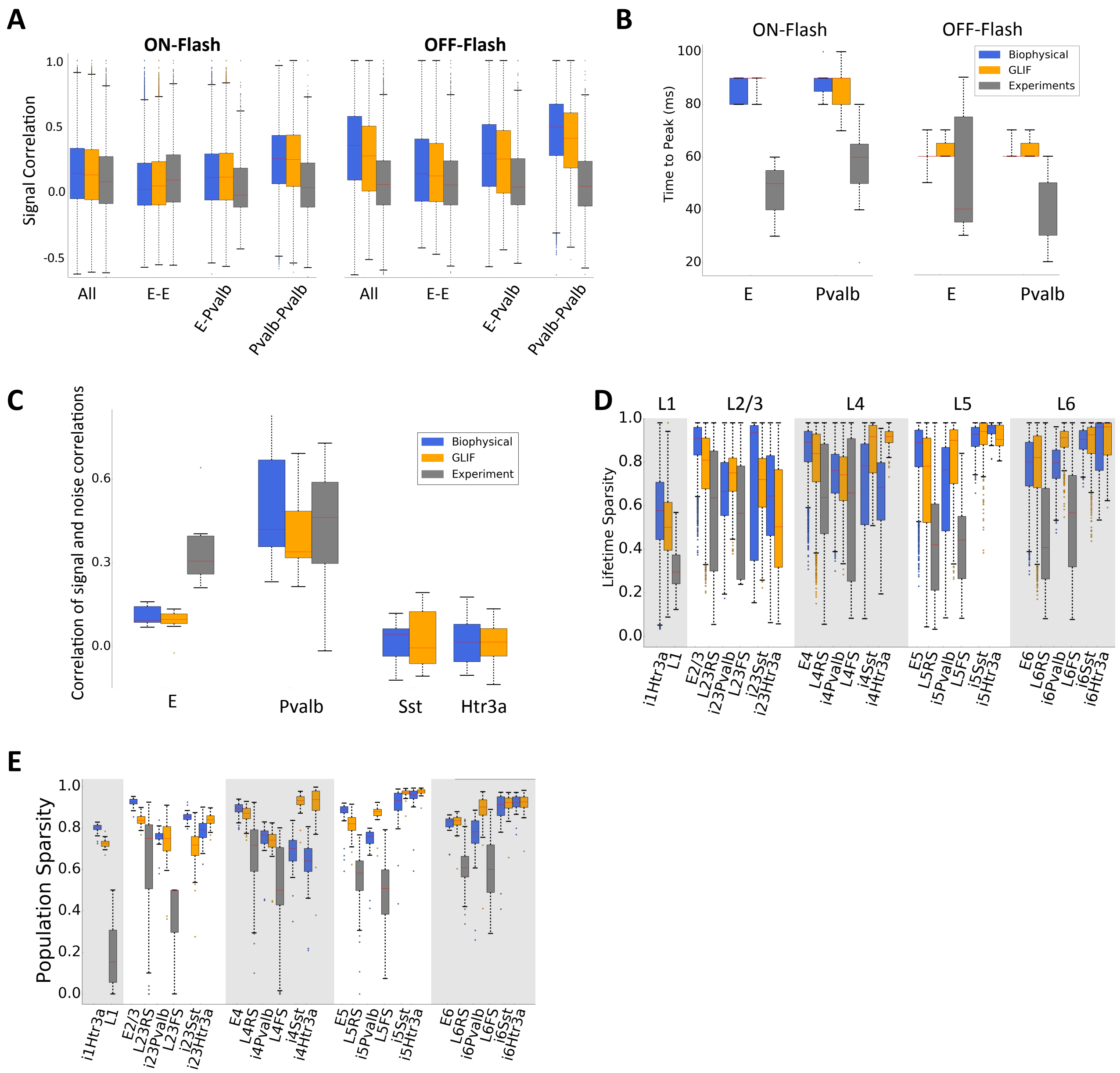


**Figure S16:** Response of the final V1 models to full-field flashes and a natural movie compared to experimental *in vivo* extracellular recordings. **A:** Signal correlations in responses to ON and OFF flashes for the models and Neuropixels experiments. **B:** Time-to-peak of responses to flashes for excitatory and Pvalb neurons in the models and experiments (regular-spiking and fast-spiking respectively). **C:** Correlation between signal and noise correlations (see Methods) for responses to a natural movie. **D:** Life-time sparsity, averaged over trials, for responses to a natural movie. **E:** Population sparsity of responses to a natural movie.

Given the availability of experimental data to the same flash stimuli and the same natural movie, we here discuss the performance of our model with these stimuli. To quantify the degree to which neurons follow the same time course in response to a global flash stimulus, we compute the signal correlations (see

Methods) between neurons. Despite the highly correlated structure of the input, neurons tend to have low correlations with each other in both models and the experimental data (Fig. S16A). The signal correlations in the models are slightly higher than, but otherwise overlap closely with the experimental ones; the Pvalb-Pvalb correlations deviate the most from experimental measurements, but are still well below 1. We speculate that could be due to a lack of heterogeneity of a background input from the rest of the brain. We also observe that the time-to-peak values for responses to full-field flashes are comparable to Neuropixels experimental recordings (Fig. S16B). This is an important indication that the dynamics of initial transformation of the visual signal is captured relatively well. The network, however, shows less variability in time-to-peak compared to the experiments.

In comparison with artificial stimuli like gratings or flashes, natural stimuli exhibit distinct statistical features and evoke highly heterogenous responses. We test our models on a clip from one movie shown to mice in the Allen Brain Observatory, which used $Ca^{2+}$ imaging to quantify responses of many neuronal populations across most layers of visual cortex (de Vries *et al.*, 2019). We compute the correlation between the signal correlations and noise correlations for spiking responses of neuron pairs from our models and *in vivo* electrophysiology recordings (for direct comparison) and find similar, almost all positive, values for models and experiment alike (Fig. S16C). As reported previously (de Vries *et al.*, 2019), this metric exhibits rather small (positive, but close to zero) values for excitatory neurons, although the spiking experimental data result in somewhat higher values than what we observe in the models. Interestingly, the fast-spiking interneurons in our experimental data exhibit substantially higher values of this metric than do excitatory neurons. This trend is captured by the models (Fig. S16C). Another major characteristic of responses to natural stimuli is the high lifetime sparsity that quantifies the selectivity of each neuron across the movie stimulus (Vinje and Gallant, 2000; de Vries *et al.*, 2019). This finding is also reproduced by our models (Fig. S16D, see Methods) albeit with higher values than our electrophysiology data. A similar, yet bigger, difference exists between the models and experiments for population sparsity, which quantifies selectivity across activity patterns in the population (Fig. S16E, see Methods). These differences could arise due to unique reasons based on specific cell classes; for example, we do not model long-range intracortical input to Layer 1 neurons that could explain the substantially higher levels of both sparsity measures observed in L1. Given the use of the natural movie in the Allen Brain Observatory, we can compare the VIP (subclass of Htr3a) neurons in L2/3 and L4 that exhibit reduced sparsity compared to excitatory and SST classes (that survey did not include Pvalb). Htr3a (or VIP) neurons are not readily identifiable in electrophysiological recordings and are arguably less well parameterized in the models (due to lower data availability) than excitatory and Pvalb classes. Nevertheless, in the biophysical model the Htr3a class does exhibit reduced sparsity in L2/3 and L4. The model shows high sparsity for Htr3a in L5 and L6 (mostly non-VIP in these layers), an observation that has not been yet tested experimentally. Investigating these similarities and differences further, as shown in this work for direction and orientation selectivity (Figs. 3 – 7), will yield insights into the mechanisms and function of cortical circuits.

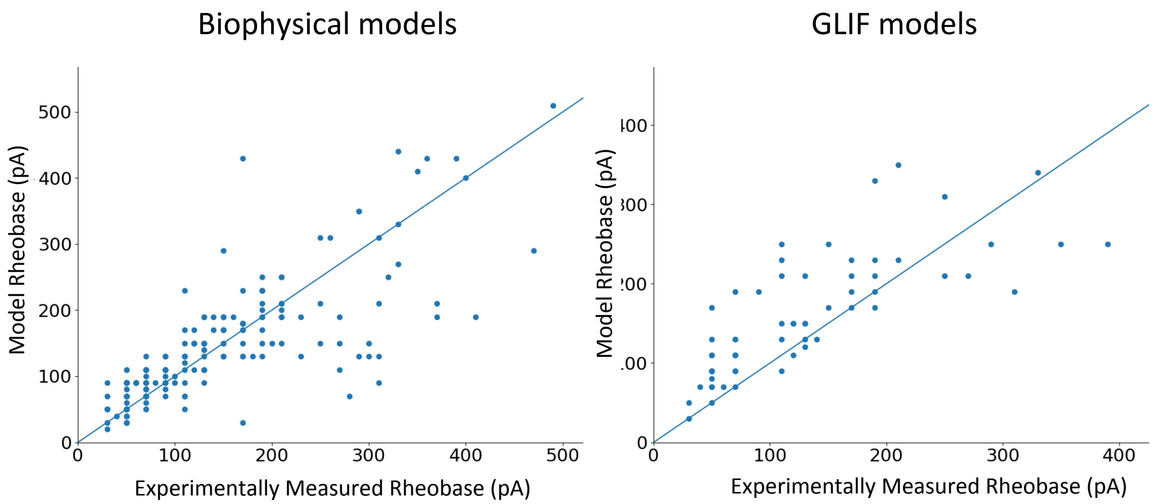


**Figure S17:** Model rheobase vs. experimentally measured rheobase for individual neuron models.

| Similarity Score | Rate | OSI | DSI |
|---|---|---|---|
| E: Biophysical-Experiment | 0.72 | 0.75 | 0.92 |
| E: GLIF-Experiment | 0.72 | 0.72 | 0.87 |
| Pvalb: Biophysical-Experiment | 0.80 | 0.33 | 0.71 |
| Pvalb: GLIF-Experiment | 0.76 | 0.44 | 0.83 |
| E: Biophysical- GLIF | 0.97 | 0.93 | 0.95 |
| Pvalb: Biophysical- GLIF | 0.82 | 0.78 | 0.79 |

**Table S1:** The similarity score values ($\mathbf{S}$) between the final V1 network models (Biophysical and GLIF) and the experiments. We use E to represent excitatory neuron comparisons (E in model with RS from experimental data) and Pvalb to represent Pvalb comparisons (Pvalb in model with FS from experimental data). Almost all comparisons show strong agreement between the models and the experiments except for OSI in Pvalb neurons. Note the high $\mathbf{S}$ values between the two model resolutions across metrics (final two rows).

**Supplementary Material 1:** To give a sense of using and running the models, the README file from our release is shown here for convenience to the reader (Ver. Oct. 14, 2019).

All data, code, and metadata are available under the Allen Institute terms of use:
https://alleninstitute.org/legal/terms-use/

================================================

1. INTRO

================================================

The data, code, and metadata for two V1 networks models are made available here.

All data is accessible with Python and ran on Python 2.7. To run simulations with the code here,

it is expected that users have NEURON and/or NEST installed depending on the application.

Please see the Brain Modeling ToolKit page for more information.

All software needed is provided here. See below for further details.

The paper describing the models is:

SYSTEMATIC INTEGRATION OF STRUCTURAL AND FUNCTIONAL DATA INTO MULTI-SCALE MODELS OF MOUSE PRIMARY VISUAL CORTEX

Yazan N. Billeh, Binghuang Cai, Sergey L. Gratiy, Kael Dai, Ramakrishnan Iyer, Nathan W. Gouwens, Reza Abbasi-Asl,

Xiaoxuan Jia, Joshua H Siegle, Shawn R. Olsen, Christof Koch, Stefan Mihalas, Anton Arkhipov

We developed data-driven models of the mouse primary visual cortex (area V1) for in silico visual

physiology studies with arbitrary visual stimuli. The models, constrained by experimental measurements have

the same network graph of 230,978 nodes of two different levels of granularity: biophysically detailed compartmental

models and Generalized Leaky Integrate and Fire (GLIF) point-neuron models. Otherwise, the two V1 models

are identical.

In total, the two V1 networks contain 17 cell classes, represented by 112 unique individual neuron models

for the biophysical and 111 for the GLIF network, copied and distributed in layers according to the best data

available. The final network contains 230,924 cells, of which 51,978 are in the core.

===============================================

2. DIRECTORY STRUCTURE

===============================================

-- Resources (meta-data) and Outputs

resources_meta_data: contains interactive files for all meta-parameters used in model building

simulation_outputs: the spiking files for drifting gratings, flashes, and natural movies

-- Biophysical_network: Directory for the biophysical V1 network resolution

biophys_components: the mechanisms and components of the biophysical network such as cell models and synaptic dynamics. See the BMTK documentation for more detail.

inputs: example LGN and background inputs to the model in a single directory (but same as GLIF below)

network: the networks files all in a single directory. Some of these files are identical to the GLIF network but are saved twice for user convenience.

output: example output for one drifting grating simulation run. If you run the network, this file is overwritten unless you change the file name.

config: configuration file for running the network

run_bionet.py: python scripts that runs the network using BMTK and the config.json file (see below for more details).

-- GLIF_network: Directory for the GLIF V1 network resolution

models: the GLIF neuron models for the simulation

inputs: example LGN and background inputs to the model in a single directory (but same as biophysical)

network: the networks files all in a single directory Some of these files are identical to the biophysical network but are saved twice for user convenience.

output: example output for one drifting grating simulation run. If you run the network, this file is overwritten unless you change the file name.

config: configuration file for running the network

run_pointnet.py: python scripts that runs the network using BMTK and the config.json file (see below for more details).

- For creating the nodes, connections, and inputs for the models

build_model: instantiates the neurons of the network

LGN: directory with the code to instantiate an LGN module, create spike-trains, and connect the LGN units to the V1 neurons

create_recurrent_connections: code to form the recurrent connections of the V1 models (recall they are identical for both models)

background: the files for the basic background Poisson input targeting all nodes

===============================================

3. META DATA (RESOURCES) TO BUILD THE MODELS

===============================================

The directory resources_meta_data contains meta-data used to constrain the building of the models

V1_structure: an xlsx file of how number of neurons and proportions in every layer were calculated

Num_TC_synapses: an xlsx file of how we determined the number of synapses each LGN cells makes on a V1 neuron in different layers

Connection_probabilities: Interactive PowerPoint with all connection probabilities (at 75 micrometers). You can click on every number to see the references used and assumptions (if any).

Connection_strengths: Interactive PowerPoint with all connection strengths (unitary PSP in mV). You can click on every number to see the references used and assumptions (if any).

V1_parameter_estimate: The guideline of parameters of V1 neuron response properties for all cell-types. This was used as a guide for optimization before the neuropixels data became available.

===============================================

4. SIMULATING THE V1 MODELS

===============================================

Both models will run with BMTK using the same API. We provide scripts for both model resolutions. Therefore this is sub-divided into two sections for biophysical and GLIF V1 network models. Note the biophysical was simulated with NEURON 7.4 and the GLIF network model was simulated with NEST 2.14. To run these networks, go to the relevant directory and run the following commands:

1. Biophysical V1 network model (note we used 384 processes - see paper for more details)

-----------------------------------------------------------------------------------------------

```
mpirun -np 384 nrniv -mpi -python run_bionet.py config.json
```

2. GLIF V1 network model (this runs on a single CPU)

-----------------------------------------------------------------------------------------------

```
mpirun -np 1 python run_pointnet.py config.json
```

================================================

5. ANALYSING SIMULATION OUTPUTS FROM THE V1 MODELS

================================================

The direction simulation_outputs contains the spike train outputs from the simulations shown in the paper for the final V1 model

simulation_outputs/biophysical: the spike-train outputs from the biophysical model for 80 drifting gratings (8 directions, 10 trials each), flashes (10 trials), the natural movie (10 trials), and the looming stimulus (10 trials).

simulation_output/GLIF: the spike-train outputs from the GLIF model for 80 drifting gratings (8 directions, 10 trials each), flashes (10 trials), the natural movie (10 trials).

plot_raster.py: Scripts that will plot the raster plot for any of the spike-trains as shown in the paper with boxes delinating the layers

calculate_metrics.py: script that will calculate the firing rates of every neuron and calculate the DSI and OSI. Outputs are saved as a dataframe when this code is run.

*.ipynb: jupyter notebooks for the analysis done for the natural movies and flashes.

================================================

6. REQUIREMENTS

================================================

The above code is run on Python 2.7 with standard scientific packages (numpy, scipy, matplotlib, hdf5).

The below requires installation of specific software not provided here (but all are open source)

--NEURON: for the biophysical model, all simulations were run on NEURON 7.4 as the simulation engine with our python code (BMTK) as the wrapper interface.

--NEST: for the GLIF model, all simulation were run on NEST 2.14.0 as the simulation engine with our python code (BMTK) as the wrapper interface.

—-BMTK: The Brain Modeling ToolKit must be installed

--allenSDK: Only needed for building the models. The AllenSDK is a python API required to build the models (instantiate neurons from the Allen Cell Types Database) as described in section 8. This is available via brain-map.org

================================================

7. FILES FROM THE MODELS

================================================

The model files that describe the network - neurons, connections, LGN input, synaptic weights, etc. are also provided in this package.

These files for LGN and Background below are share between both the biophysical and GLIF model and are not saved twice to save on memory.

This section described those file. The code to generate these files is also provided and described individually below.

1. LGN

-----------------------------------------------------

LGN/LGN/lgn_full_col_cell_models_3.csv: contains all the LGN filter models used (Fig. 2A in the paper)

LGN/LGN/lgn_full_col_cells_3.csv: contain very LGN unit (17,400) with their coordinates and parameters

LGN/trains/grating_3.0sec_SF0.04_TF2.0_ori90.0_c80.0_gs0.5_spikes.nwb: LGN incoming spike-train for 10 trials of a drifting grating moving in the 90-degree direction. The first 0.5 seconds is grey-screen before the grating appears for 2.5 seconds (3 seconds total)

LGN/trains/fullField_250ms_1000msISI_c80_spikes.nwb: Full field flash incoming LGN spike-train for 0.5 grey screen, 250ms ON flash, 1 second grey screen, 250ms OFF flash, and a final period of grey screen. There are 10 trials.

LGN/trains/naturalmovie_graycorr_1000Hz_spikes.nwb: Natural movie LGN spike-train with 10 trials. The first 0.5 seconds is grey-screen before the grating appears for 2.5 seconds (3 seconds total).

Note the .nwb files can be opened with an HDF5 viewer (see section 9.3).

This directory also contains the script for generating the LGN filter units, spike-trains, and connecting them to the V1 network. This is described in section 9 below.

2. Background

-----------------------------------------------------

background/BKG/bkg_nodes.csv: contains the single background node that projects to all V1 neurons.

background/BKG/bkg_node_types.csv: A node_types file is needed for BMTK and this basic csv file describes it for the single background node.

background/trains/bkg_spikes_n1_fr1000_dt0.25_100trials.nwb: Spike-train with 100 trials of the background node firing at 1kHz. The first 80 trials are used for the drifting gratings, then for the natural movie and finally for the flashes. Note the .nwb file can be opened with an HDF5 viewer (see section 9.3).

3. V1 Neurons

-----------------------------------------------------

Two files describe the network. A nodes file and a node_types file. This is because we have >100 models but > 230,000 neuron models and so avoid saving multiple parameters. The nodes file contains every neuron with its unique attributes (e.g. spatial coordinates). The nodes file also contains a model-ID that points the node_types file. This will indicate the values for that neuron's model. See the BMTK or SONATA manuscript for more details. Finally, since we have different neuron models in both resolutions, we provide both files here

V1 - biophysical: files can be found in Biophysical_network/network/

V1 - GLIF: files can be found in GLIF_network/network/

4. Connectivity

-----------------------------------------------------

Both models have the same adjacency matrix (graph), though they need different synaptic weights and hence we provide both model files separately here Biophysical_network/network and GLIF_network/network

===============================================

8. BUILDING THE  V1 MODELS

===============================================

This directory described the build_model directory. Note that the code creates the biophysical instantiation. This is then converted to the GLIF files since both models are identical. Below is the description for generating the biophysical model node and node_types files.

Constructing the nodes

======================

Directory:

    build_model/cells_peri

Script:

    construct_bio_models_prop.ipynb

Inputs:

    processing_cell_models_perimodels_props.csv

    processing_cell_models_peritaus.csv

Outputs:

./biophysical/morphology - contains swc files

./biophysical/electrophysiology      - contains json files with fitted parameters

./biophysical/bio_models_prop.csv - contains comprehensive properties for biophysical models

Procedure:

Manually remove the rows with missing data from models_prop.csv and save as models_prop_nomissing_data.csv

Run construct_bio_models_prop.ipynb in order to combine model properties in models_prop.csv with those obtained from allensdk.core.cell_types_cache, lims_utils and taus.csv. Compute the depth range [detal_ymin, delta_ymax] spanned by each specimen's morphology along the depth axis. The distances [delta_ymin, delta_ymax] are measured relative to the soma. For convenience copy the swc morphologies and json parameter fits into the directories:

./morphology - contains swc files

./electrophysiology   - contains json files with fitted parameters

Manually copies the output data into cell_models/biophysical to use for network building.

Constructing the table of properties for intfire models

=======================================================

Directory:

build_model/V1

Script:

average_model_lifs.py

Inputs:

pop_queries.json - high-level description of the network populations of cells to be build.

Outputs:

build_model/cells_peri/intfire/lif_models_prop.csv - properties of lif models

./bio2lif_mapping.csv  - mapping between biophysical and lif models

Procedure:

Construct the mapping file bio2lif_mapping.csv from biophysical to lif models. The script average_model_lifs.py averages the time constants of the biophysically detailed cells and creates an average tau that is used as a parameter for the lif model. Currently we build a single lif model per population.

Building network nodes (cells)

==============================

Directory:

build_model/V1

Script:

build_nodes.ipynb

Inputs:

pop_queries.json - high-level description of the network populations of cells to be build.

Outputs:

v1_nodes.csv - positions and models of the cells created based on net_description.

v1_node_types.csv - properties of the models for needed by the simulator.

Procedure:

Run script build_nodes.ipynb

1. Setting up a high-level description of populations

-----------------------------------------------------

The populations in net_description.json are defined as e.g.:

```
"i4Sst":   {"ncells":2384,
```

```
                    "labels": {"ei":"i", "location":"VisL4"},

                    "depth_range":[310,430],

                    "criteria":{"ei":"i", "location":"VisL4", "cre_line":["Sst"],
"express":True}},
```

where "criteria" are used to choose models from the available models in the bio_models_prop.csv. The number of cells in each population is proportional to our expectation about the relative occurrence of each cre-line in mouse V1. The numbers of cells are pre-calculated in the file column_structure.xls. For the domain we use a right circular cylinder with a total radius of 845 um. The core cylinder within 400 um is populated with the biophysical cell models and the remaining annulus with the lif models.

2. Assign somatic coordinates

-----------------------------

Cells for each population are uniformly distributed within a cylindrical domain and within the specified "depth_range".

3. Assign rotation_angle_yaxis

------------------------------

Uniformly draw rotational_angle_yaxis for each cell.

4. Assign cell models

---------------------

Assign model to cells based on their y_soma position. A model may be assigned to a particular cell if that model's morphology does not significantly stick out of the pia when placed at the cell's somatic location. Currently, we allow for dendrites to stick out of the pia with a tolerance = 100 um because there are not enough short L23 excitatory cells.

Determine the acceptable yrange = [ymin,ymax] for each model based on its dendritic span [delta_ymin,delta_ymax].

For a given cell position, find models for which y_soma is in yrange.

Randomly choose from the allowed models with a Gaussian (prob = np.exp(-(y_soma-y_spec)**2/length_sigma**2/2)) probability density function (length_sigma = 20 um). The probability density function is chosen such that we preferentially choose a model, which has a y_spec closest to the y_soma of a particular cell.

For the periphery we substitute the biophysical models with the lif models according to the mapping bio2lif_mapping.csv

5. Test and visualize the node construction

------------------------------------------

* Build a network with one cell per model, construct and save segment coordinates of each morphology to build_model/cells_peri/biophysical/morph_segs

* analyze/plots the depth distributions of the models and their morphologies with build_model/V1/plot_model_distr.ipynb

===============================================

9. PROVIDING LGN INPUT TO THE  V1 MODELS

===============================================

This section described the scripts for generating the LGN input. The files are described in section 7 of this README file. Since the LGN is shared between both model resolutions, there is a single directory.

The scripts are in LGN/scripts

1. Create the LGN module

-----------------------------------------------------

Script:

generate_lgn_full_col.py

Inputs:

high-level description of the network populations and number of cells desired. This is input directly in the script.

Outputs:

two files that descrive the LGN nodes and the LGN node_types. As provided, the save commands are commented.

Procedure:

run the script and all sub-directories are already pointed too such as lgnmodel/ and network2/. The script lgn_functions.py contains many utility functions that are called on by this script.

2. Connect the LGN module to the V1 neurons

-----------------------------------------------------

Script:

connect_LGN_to_V1.py

Inputs:

the lgn_models and dimensions of the LGN visual space need to match the generate_lgn_full_col.py function.

Outputs:

an hdf5 file that described which LGN units connect to which V1 neurons.

Procedure:

run the script and all sub-directories are already pointed too such as lgnmodel/ and network2/. The script lgn_functions.py contains many utility functions that are called on by this script that also contains a call to the parameter dictionary.

3. Generate spike trains from the LGN model

-----------------------------------------------------

Script:

simulate_drifting_gratings.py

Inputs:

the LGN module files created by generate_lgn_full_col.py. This is input directly in the script.

Outputs:

two hd5f files. One for the deterministic firing rate and the other for the spike trains. The spike train extensions are .nwb from the first version of the Neurodata Without Borders. The most recent SONATA format are provided in the input-subdirectories of the Biophysical_network and GLIF_network directories.

Procedure:

run the script and all sub-directories are already pointed too such as lgnmodel/ and network2/. The script lgn_functions.py contains many utility functions that are called on by this script.

===============================================

10. PROVIDING BACKGROUND INPUT TO THE V1 MODELS

===============================================

The background in a single node that connects to all neurons that fires with a Poison process at 1kHz. As described in section 7, the necessary files are:

background/create_poisson_spike_train.py: script that creates the spike train. Note the path need to point to the GLIF directory also to call the nwb module from version 1.0.

background/BKG/bkg_nodes.csv: contains the single background node that projects to all V1 neurons.

background/BKG/bkg_node_types.csv: A node_types file is needed for BMTK and this basic csv file describes it for the single background node.

background/trains/bkg_spikes_n1_fr1000_dt0.25_100trials.nwb: Spike-train with 100 trials of the background node firing at 1kHz. The first 80 trials are used for the drifting gratings, then for the natural movie and finally for the flashes. Note the .nwb file can be opened with an HDF5 viewer (see section 9.3).

===============================================

11. RECURRENTLY CONNECTING THE V1 MODELS

===============================================

This section describes the scripts needed to create the connectivity between V1 neurons (i.e. recurrent connections).

1. create V1-to-V1 connections

-----------------------------------------------------

Script:

build_CC_connections.py

Inputs:

the V1 nodes files to be connected.

Outputs:

v1_edges.h5 file that contain the graph of connectivity. This was described in section 7.4

Procedure:

run the script and all sub-directories are already pointed too such network2/. The script connect_cells.py contains all the functions that create the connectivity such as distance dependence, like-to-like connectivity and the python dictionaries with all the relevant meta-parameters described in section 3

===============================================

12. CONTACT

===============================================

We can be contacted at
